# Phenotypic differentiation between highland and coastal quinoa under cold stress conditions

**DOI:** 10.64898/2026.01.23.701318

**Authors:** Niharika Rakasi, Lydia Kienbaum, Katharina B. Böndel, Jan-David Wiederstein, Naveen Kumar Ganga Raju, Sandra M. Schmöckel, Karl J. Schmid

## Abstract

Quinoa (*Chenopodium quinoa* Willd.) is a genetically diverse Andean crop valued for its nutrition and adaptability to varied agro-climatic conditions with potential for cultivation in European and Mediterranean, particularly on marginal lands. Low temperatures during early sowing can impair germination, while delayed sowing increases the risk of poor maturation due to unfavorable autumn weather. To assess the adaptation of quinoa to low temperature conditions, that reflect cold stress, we evaluated germination and phenotypic variation in 60 accessions from highland and coastal ecotypes across three sowing dates in South-Western Germany: late winter (S1), early spring (S2), and spring (S3). Early sowing under low temperature conditions in S1 delayed seedling-emergence and reduced emergence percentages, yet these plants produced the highest average seed yield per plot (64 g) compared to S2 (46 g) and S3 (35 g). Highland accessions showed earlier seedling-emergence and with higher emergence percentages, while coastal types matured earlier and gave higher yields across sowing dates. A complementary laboratory experiment assessed germination under cold (4.4 °C) and control (18.3 °C) conditions, using both manual scoring and image analysis via a Mask R Convolutional Neural Network, to track seedling growth. This confirmed the beneficial germination performance of highland accessions under low temperature conditions, with strong agreement between manual and automated scoring. Our findings suggest that quinoa demonstrates resilience to cold stress with highland quinoa exhibiting superior germination traits, and early sowing, despite reduced emergence, can lead to higher yields. We conclude that combining favorable traits such as faster maturity and higher yield of coastal ecotypes with superior germination traits of highland accessions is a promising avenue for breeding improved quinoa varieties for cold climatic regions.

## INTRODUCTION

As the global population continues to grow, the need for increased food production becomes even more pressing. Furthermore, in recent decades environmental changes have intensified significantly, posing an imminent threat to the global food supply as it affects the crop yield, production and quality (Myers et al., 2017). Most crop species are adapted to past climate conditions and may experience decreased productivity and even extinction as we move towards the future marked by an escalating climate change (IPCC, 2023). Therefore, exploring and embracing minor and underutilized crops in agriculture as a strategic response to the challenges posed by a changing climate holds great value for future food security (J. González et al., 2015; Kamenya et al., 2021).

Quinoa (*Chenopodium quinoa* Willd.) is an annual grain crop, native to the Andean region of South America, and has the potential of an alternative crop in various regions across the globe. It is assumed that the domestication of quinoa happened over 7,000 years ago (Bazile et al., 2016). The original cultivation area of quinoa ranges from 2°N latitude in Southern Colombia to 40°S latitude in Chile, where it is cultivated in wide range of altitudes from sea level up to 4,000m above sea level (Jacobsen, 2003; Jacobsen et al., 2015; Jacobsen & Stølen, 1993; Risi & Galwey, 1989). In further consecutive studies quinoa was classified into two main types: the Andean highland type and the Chilean coastal type using morphometric, electrophoretic and genetic data (Christensen et al., 2007; Wilson, 1988; Zhang et al., 2017; Zurita-Silva et al., 2014). Quinoa is commonly classified as bitter or sweet depending on the presence or absence of saponins, chemical compounds that are primarily concentrated in the seed hull (Troisi et al., 2015).

The climatic conditions of the Andean regions are characterized by several adverse factors, including drought, salinity, frost, wind, hail, flooding, high UV irradiation, and heat (Jacobsen, 2003). Quinoa evolved under these harsh climatic conditions, and therefore demonstrates high levels of tolerance to several abiotic stresses (Jacobsen et al., 2003; Trognitz, 2003). Multiple studies have reported that quinoa shows tolerance to frost (Jacobsen et al., 2005, 2007), salinity (Adolf et al., 2012, 2013), and drought (Jacobsen et al., 2015; Razzaghi et al., 2011, 2012). The nutritional profile of quinoa seeds consists of a high protein content, abundant essential amino acids, oil content, starch, dietary fiber, antioxidants, minerals, and vitamins (Graf et al., 2015; Repo-Carrasco-Valencia et al., 2003; Repo-Carrasco-Valencia & Serna, 2011). It is also naturally gluten-free and has a low glycemic index (Bastidas, et al., 2016). In recognition of its potential to contribute to global food and nutritional security, the Food and Agriculture Organization (FAO) of the United Nations declared 2013 the International Year of Quinoa (United Nations, 2011).

Most of the quinoa production originates from Bolivia and Peru with small contributions from Ecuador, Chile, Argentina, and Columbia. Due to its beneficial properties, the popularity of quinoa has increased in recent years and attempts are made to acclimatize quinoa to North American, European and Asian countries (Bazile & Baudron, 2015; Jacobsen, 2003). The number of countries growing quinoa for either experimental or production purposes increased from just seven Andean countries in 1980 to more than 125 in 2020 (Bazile et al., 2021). Quinoa might contribute to diversified cropping systems as an alternative crop for marginal lands and low-input farming in Europe and other parts of the world. Previous research evaluated the adaptation of quinoa to environments outside its geographic origin (Jacobsen, 2003). Under European conditions, a short growth period is regarded advantageous because the harvesting stage of late maturing genotypes may coincide with unreliable autumn weather, which affects the seed quality and crop yield. A potential solution to achieve early maturity is to sow early and extend the growing season by which the productivity could also be increased (Christiansen et al., 1999). In Europe, the recommended time for sowing of quinoa is April or May when soil temperatures are about 8-10 °C. For the early establishment of the crop, seeds should be sown during March, but the low temperatures could affect germination (Jacobsen, 2003; Jacobsen & Stølen, 1996). Furthermore, in the Mediterranean region, which is characterized by an extremely variable climate with hot, dry summers and cold, wet winters (Ceccarelli et al., 2007; Jacobsen et al., 2012), the planting season for quinoa ranges from October to April. Temperature during winter and early spring will be colder and the crop will be exposed to frost and low temperatures during early stages, which can impact its growth and yield. Given that both main types of quinoa are cultivated in very different climatic environments, the investigation of differences local adaptation to cold temperatures may identify useful genotypes for future breeding programs. Previous work on cold stress in highland quinoa cultivars indicated a germination ability at very low temperatures of 2°C (Bois et al., 2006). Furthermore, quinoa is tolerant to frost during its early growth stages, but the final yield of plants exposed to frost is reduced (Jacobsen et al., 2005). So far, most cold stress experiments were conducted either in lab or under greenhouse conditions and it needs to be validated whether these observations are relevant for field cultivation. Although some experiments were performed to investigate the effects of early and late sowing dates on seed yield (Hirich et al., 2014; Iliadis et al., 1999; Öktem et al., 2020; Risi & Galwey, 1991), studies on the effects of low temperature on germination of quinoa seeds are lacking.

Therefore, the aim of this research was to investigate the effects of low temperature on quinoa germination and to assess how different sowing times influence crop growth and yield under field conditions. We hypothesized that lowland and highland accessions differ in their adaptation level to low temperature conditions during the early stages of development, reflecting the environmental conditions of their respective cultivation regions. To test this, we evaluated a representative sample of 60 quinoa accessions from coastal and highland ecotypes for cold stress tolerance. Experiments were conducted under field conditions at three different sowing dates on the same field in South-Western Germany and under controlled laboratory conditions. We chose this dual approach to enhance the robustness and practical relevance of our research: field trials offer realistic insights into plant stress responses in natural environments, while laboratory experiments allow for precise control of variables and reduced environmental noise. For the laboratory experiments, we developed an automated image analysis pipeline using convolutional neural networks (CNN), as they have been previously shown to be very useful for plant phenotyping. Image analysis employing deep learning techniques has become state of the art in high-throughput phenotyping, addressing the bottleneck created by rapid advances in genomic data generation (Katal et al., 2022; Murphy et al., 2024; Song et al., 2021; Yang et al., 2020). The automatic analysis of germination experiments has several advantages: It is highly reproducible compared to manual counting of germinated seeds, cost-efficient and offers possibilities beyond a simple counting (e.g., measurement of surface area of seedlings as a proxy for seedling growth). Due to its computational efficiency and general performance in multitask settings, and a high number of successful use cases in plant phenotyping (Fan et al., 2023; Mostafa et al., 2023; Seki & Toda, 2022; Y.-H. Wang & Su, 2022), we applied Mask R-CNN convolutional neural network in this study for the analysis of germination traits in laboratory under cold stress conditions in quinoa. The results from our field trials and laboratory experiment reveal clear phenotypic differences between low- and highland accessions and a high congruency of phenotypic evaluation under field and laboratory conditions indicating differential adaptation of coastal and highland quinoa.

## MATERIALS AND METHODS

### Plant material and genetic data

We analyzed 60 quinoa (*Chenopodium quinoa* Willd.) genebank accessions obtained from the United States Department of Agriculture (USDA; https://npgsweb.ars-grin.gov/gringlobal/search) and the Leibniz Institute of Plant Genetics and Crop Plant Research (IPK; https://www.ipk-gatersleben.de/en/genebank/), accessed on 12 August 2025. The accessions span a wide altitudinal range and collectively represent the diversity of quinoa cultivation areas across South America (Data S1). Seeds were produced during the 2018 growing season at the Heidfeldhof research station near the University of Hohenheim, Germany (48°51′ N, 7°88′ E; 389 m a.s.l.). To prevent cross-pollination, given an outcrossing rate of up to 18% (Murphy et al., 2016) panicles were bagged before flowering to ensure self-pollination. Sixty seeds per accession were used in the experiments.

Whole-genome resequencing data for 55 of these accessions are publicly available in the NCBI Sequence Read Archive (BioProject PRJNA673789). Details on DNA extraction, sequencing, read mapping, and SNP calling are provided in Patiranage et al., (2022). Variant data were extracted using vcftools v0.1.17 (Danecek et al., 2011) with a minimum read depth of 5 and ≤20% missing data per site. To minimize linkage disequilibrium, SNPs were pruned using PLINK v2.0 (Purcell et al., 2007) with a 50-SNP window, step size 5, and *r²* threshold 0.5. As shown previously (Patiranage et al., 2022), quinoa accessions form two major genetic clusters corresponding to highland and coastal origins. To verify this structure, we conducted principal component analysis (PCA) using the R package SNPRelate v1.34.1 (Zheng et al., 2012), which enabled assignment of the previously genotyped 55 accessions to either cluster for subsequent analyses. The final dataset comprised 45 highland and 15 coastal accessions (Data S1). Although this resulted in unequal group sizes, all analyses were conducted using mixed-model approaches that accommodate unbalanced designs.

### Field experiment

#### Experimental design

The field experiment was conducted from March to October 2021 at the Heidfeldhof research station near the University of Hohenheim, Germany (48°51′ N, 7°88′ E; 389 m a.s.l.), on slightly pseudogleyed brown earth (uL) with a soil value 50-60. A randomized complete block design (RCBD) with three replicates was used (Figure S1). In each replicate, 60 accessions were randomly assigned to 60 plots, and seeds were sown in three rows corresponding to three sowing dates that represented different temperature regimes. The first sowing (4 March) occurred at a mean soil temperature of 5 °C, followed by sowings on 1 April and 30 April at mean soil temperatures of 8 °C and 11 °C, respectively (Table S1). Climatic data for the entire period were obtained from the agrometeorological station located on-site (https://www.wetter-bw.de; accessed 12 August 2025; Table S2). Sowing was performed manually with 10 seeds per accession at each time point and replication. At S3, a modified sowing strategy was implemented to ensure establishment under late sowing conditions, to ensure comparable establishment success across treatments. Following emergence, phenotypic observations were taken from single selected plant per hill (Table S1). Plots were spaced 1 m apart, with 75 cm between rows and 25 cm between plants; each row measured 2.5 m in length, and sowing dates were separated by 75 cm. Weeds were controlled manually and mechanically as required, and plants were harvested by hand between 1 September and 31 October upon reaching maturity.

#### Data collection

Nine agronomic traits were recorded following standard quinoa descriptors (Bioversity International et al., 2013; Table 1). Germination was defined as seedling emergence when cotyledons were visible above ground. Emergence counts were taken daily for 30 days after sowing and used to calculate mean emergence time and emergence percentage. Panicle emergence, flowering, and maturity were recorded when at least 50% of plants within a plot reached the respective stage. Plant height, panicle length, and panicle width were measured at physiological maturity using a ruler. At maturity (defined by hard seed texture), plants were harvested manually 2–3 cm above ground, dried at 60 °C for 48 h, threshed, cleaned, and weighed to determine seed yield per plot. Data from individual plants (Data S2) were averaged per plot for each sowing date and replication (Data S3).

**Table 1.**
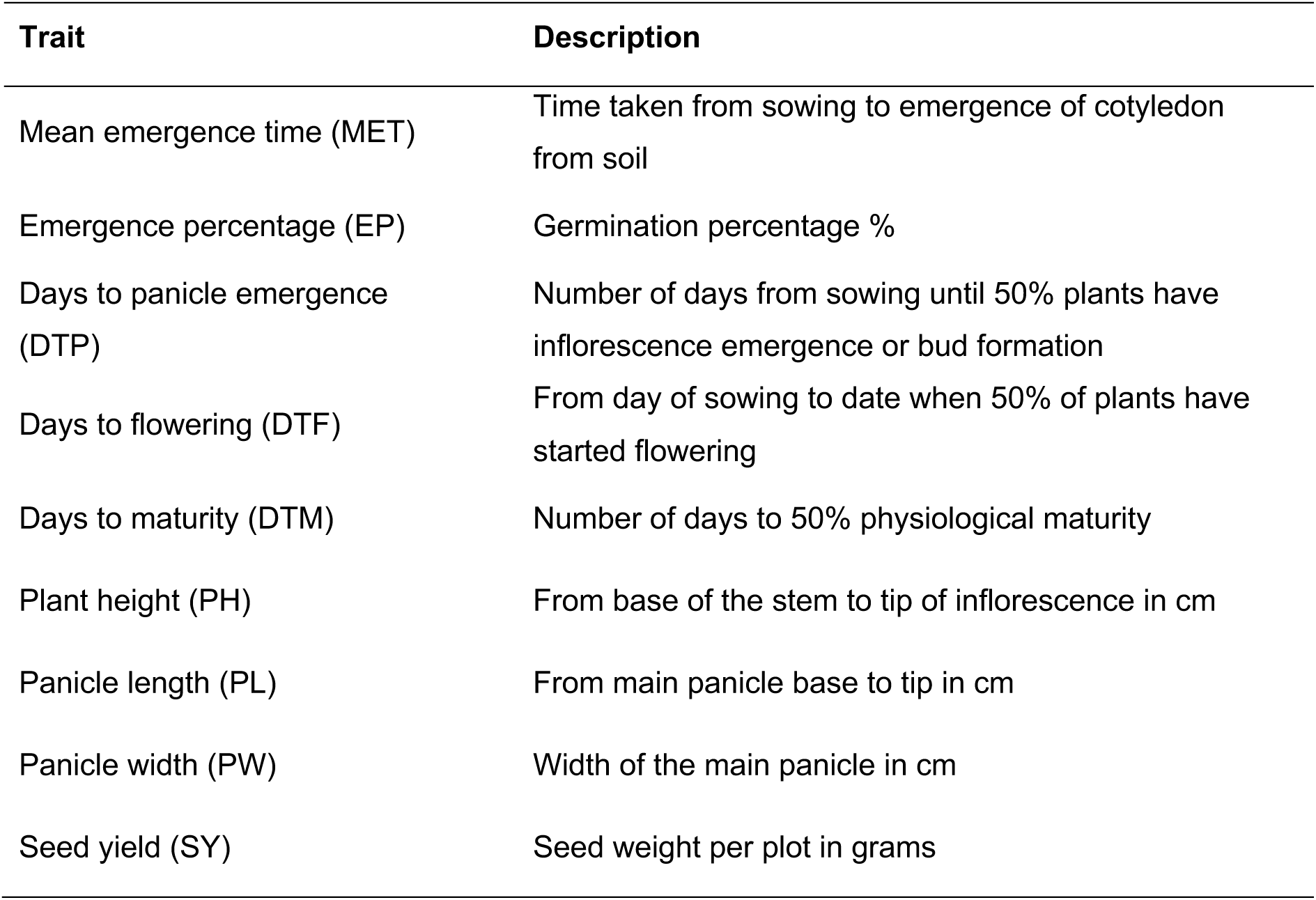
Traits recorded in the field trial for each quinoa accession.

Thermal time was quantified as growing degree days (GDD) to compare developmental stages across temperature regimes (Miller et al., 2018). GDD was calculated as:

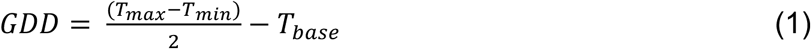

where 𝑇_max_ and 𝑇_min_ are the daily maximum and minimum air temperatures, and 𝑇_base_ (3 °C for quinoa; Jacobsen & Bach, 1996) is the threshold below which growth ceases. GDD values were accumulated daily until each developmental stage was reached.

### Laboratory germination experiment

#### Setup and manual scoring

To control for environmental factors that may have influenced field germination (e.g. humidity, soil, day length, or seed predation), a laboratory experiment was conducted to compare germination under cold (4.4 °C) and standard (18.3 °C) conditions. Seeds were placed in 9 cm Petri dishes and kept in the dark. Each treatment was replicated three times over time, using ten seeds per accession per replicate (30 seeds per treatment). To prevent mold, seeds were surface-sterilized at 50 °C for 10 min by submerging 2 ml tubes containing seeds in a 50 °C water bath. After sterilization, seeds were distributed on filter paper moistened with 6 ml autoclaved water, dishes were sealed with Parafilm® M, and incubated under either cold or control conditions. Germination was monitored manually and via photographic imaging every 24 h for 29 days (cold) and 8 days (control), based on cumulative germination patterns from the first replication (Figure S2). Manual counts recorded the number of germinated and viable seedlings, with viability defined by visible root, shoot, and leaf development. Germination data for all 60 accessions are provided in Data S4.

### Automated phenotyping using image analysis

#### Development of a deep learning image analysis pipeline

To automate germination monitoring and seedling growth measurement, we developed a deep learning image analysis pipeline based on a standardized imaging setup (Figures S3a, b). The dataset comprised a total of 6,937 images representing 60 accessions, three replications and two treatments (cold and room temperature). A total of 5,459 images were obtained under cold treatment across 31 days, while 1,478 images were taken under room temperature conditions over 12 days. Overall this resulted in at least 115 images with 10 seeds each per accession over all replications and treatments. The pipeline used a Mask R-CNN model for image segmentation and object detection (Kaiming et al., 2017); model development and evaluation are detailed in the Supplementary Text. Twelve candidate models were trained on annotated images following the configurations of Kienbaum et al., (2021) and Lozano-Isla et al., (2025), varying only hyperparameters affecting small-object detection (Table S3). Each configuration was trained five times, and performance was evaluated using mean average precision (mAP@[0.5:0.95]) and a custom penalty score for misclassification. Based on these metrics, models 8-C and 11-D performed best and were applied to the full dataset of 6,937 images to quantify seeds, seedlings, and surface area using ruler elements for scale. Model 8-C produced fewer misclassifications and was selected for subsequent analyses. Because overlapping seedlings in later growth stages caused double-counting, only images with ≤10 detected objects were retained for calculating days to germination and germination percentage (Data S5). To avoid overestimation caused by overlapping seedlings and duplicate detections, images containing more than ten detected objects were excluded from downstream analyses. This filtering primarily affected later developmental stages, where increased seedling overlaps reduced segmentation accuracy, and may therefore influence estimates of later growth dynamics more strongly than germination parameters. To test model robustness, we applied model 8-C and an updated version to a separate set of germination images from an independent experiment (Figure S3e).

#### Growth rate estimation

Temperature can influence not only germination but also post-germination growth. To assess differences among accessions, we calculated maximum seedling growth rates for each replicate and temperature condition. Growth rate was estimated as the slope of a linear regression between log-transformed average seedling surface area and time, using data extracted from the image analysis pipeline. For room temperature and cold treatments, growth rates were calculated up to day 3 and day 15, respectively, when growth curves began to plateau due to seedling overlap and reduced segmentation accuracy.

#### Seed size measurements

Seed size was assessed for 59 of the 60 accessions using two approaches: (1) the MARViN Seed Analyzer System (MARViTECH GmbH, Wittenburg, Germany) and (2) the automated image analysis pipeline. For MARViN, seeds from three successive generations (*G₀, G₁, G₂*) were measured (Data S6). Values were standardized by subtracting generation means to obtain relative seed size. For image-based measurements, seed surface area was extracted from day 1 images of the laboratory germination experiment.

### Data analysis

The data analysis was conducted with R version v4.2.2, (R Core Team, 2022).

#### Field experiment

To identify factors influencing phenotypic responses, we fitted a mixed model treating sowing dates as repeated measures (Piepho et al., 2004):

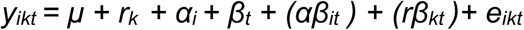

where *y_ikt_* is the response of the *i*-th genotype in the *k*-th replication at the *t*-th sowing date; μ is the overall mean; *α_i_* and *β_t_* are the effects of replication, genotype, and sowing date, respectively; *(αβ_it_)* and *(rβ_kt_ )* are interaction terms; and *e_ikt_* is the residual error. All effects except *e_ikt_* were treated as fixed. Temporal correlations among sowing dates within the same plot were modeled using a first-order autoregressive [AR(1)] structure:

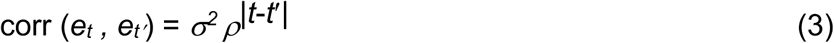

where ρ is the autocorrelation parameter (Piepho, 2019).

Wald F-tests were used to evaluate fixed effects. Best linear unbiased estimates (BLUEs) were calculated for each trait assuming fixed genotypic effects, and best linear unbiased predictions (BLUPs) were estimated by treating genotypes as random effects. Analyses were performed in ASReml v4.1.0 (Butler, 2018). Broad-sense heritability 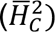 was estimated following Cullis et al., (2006):

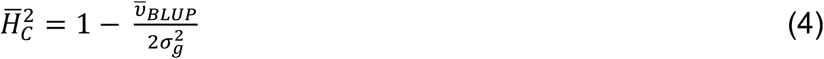

where 𝜐̅_BLUP_ is the mean variance of BLUP differences and 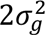 is the genotypic variance (Piepho & Möhring, 2007).

Pearson’s correlations among traits were calculated using BLUEs from PerformanceAnalytics v2.0.8 (Brian G. & Peter, 2024). Genotypic stability was assessed using the Multi-Trait Stability Index (MTSI), Weighted Average of Absolute Scores (WAAS), and WAASBY indices, implemented in metan v1.19.0 (Olivoto & Lúcio, 2020). To explore trait relationships and variation among accessions, a principal component analysis (PCA) was conducted using FactoMineR v2.11 (Husson et al., 2008).

#### Laboratory experiment

The laboratory experiment followed a split-plot design with two temperature treatments (cold and control) as the main-plot factor and 60 quinoa accessions as randomized subplots. Germination data were analyzed using the mixed model:

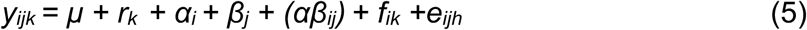

where *y_ikt_* is the response of genotype *j* under treatment *i* in replication *k*; μ is the overall mean; *r_k_* is the effect of replication; *α_i_* , *β_j_* and *(αβ_ij_)* represent treatment, genotype, and their interaction; *f_ik_* is the main-plot error; and *e_ijh_* is the residual error (Piepho, 2019). Wald F-tests assessed fixed effects, and best linear unbiased estimates (BLUEs) were computed for all traits assuming fixed genotypic effects. Analyses were performed in ASReml-R v4.1.0 (Butler, 2018). Mean differences between treatments and between coastal and highland ecotypes were evaluated using *t*-tests. Pearson correlations based on BLUEs were used to compare germination traits between field and laboratory experiments, and between manual and image-based scoring.

#### Seed size measurements

Seed size variation was analyzed using three mixed models. MARViN data, which included three generations (*G₀–G₂*), were fitted with:

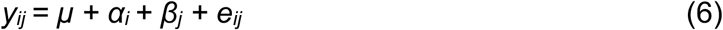

where *y_ij_* is the response of genotype *i* in experiment *j*; *α_i_* and *β_j_* denote genotype and experiment effects, respectively. BLUEs were calculated assuming fixed genotypic effects, and coastal–highland comparisons were based on these estimates. Best linear unbiased predictions (BLUPs) were obtained by treating genotypes as random, and broad-sense heritability was derived from the BLUPs.

#### Comparison between image analysis and MARViN data

To assess consistency between methods, seed surface area estimates from MARViN (generation 1) and the image analysis pipeline were correlated. Since MARViN lacked replication, two models were fitted. For image analysis:

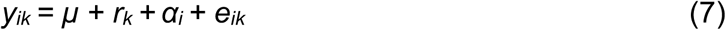

where *y_ik_* is the response of genotype *i* in replication *k*. μ is the overall mean, *r_k_* is the effect of replication *k*, *α_i_* is the fixed effect of genotype *i*, and *e_ik_* is the residual error. All effects were treated as fixed except for *e_ik_*. For the MARViN data:

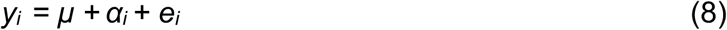

where μ is the overall mean, *α_i_* the genotypic effect, and *e_i_* the residual error.

## RESULTS

### Population structure analysis confirms two distinct genetic groups

Understanding how highland and coastal quinoa ecotypes differ in their responses is central to identifying genetic adaptation and the identification of genotypes for later breeding purposes. To address this, we selected 60 accessions representing the overall quinoa diversity and categorized them into ecotypes based on the geographical information in their passport data. To genetically validate this classification, we performed a principal component analysis (Figure 1) using 55 accessions with available whole-genome resequencing data (NCBI Sequence Read Archive, BioProject PRJNA673789). The first and second principal components together explained 14.2% of the total variance (PC1 = 8.2%; PC2 = 6.0%). The analysis revealed two main genetic groups that largely corresponded to the geographical origin of accessions and confirm the categorization based on passport data. Accessions from Peru and Bolivia clustered as highland quinoa, while those from Chile and the USA formed a distinct coastal group. Only two Peruvian accessions clustered with the Chilean and US accessions. Based on this pattern, we assigned the five accessions lacking genetic data to ecotypes according to their geographical information, resulting in 45 highland and 15 coastal accessions. The larger number of highland accessions reflects the composition of the available genebank accessions and a broader representation of accessions originating from the highlands of Peru and Bolivia than from coastal areas.

**Figure 1.**
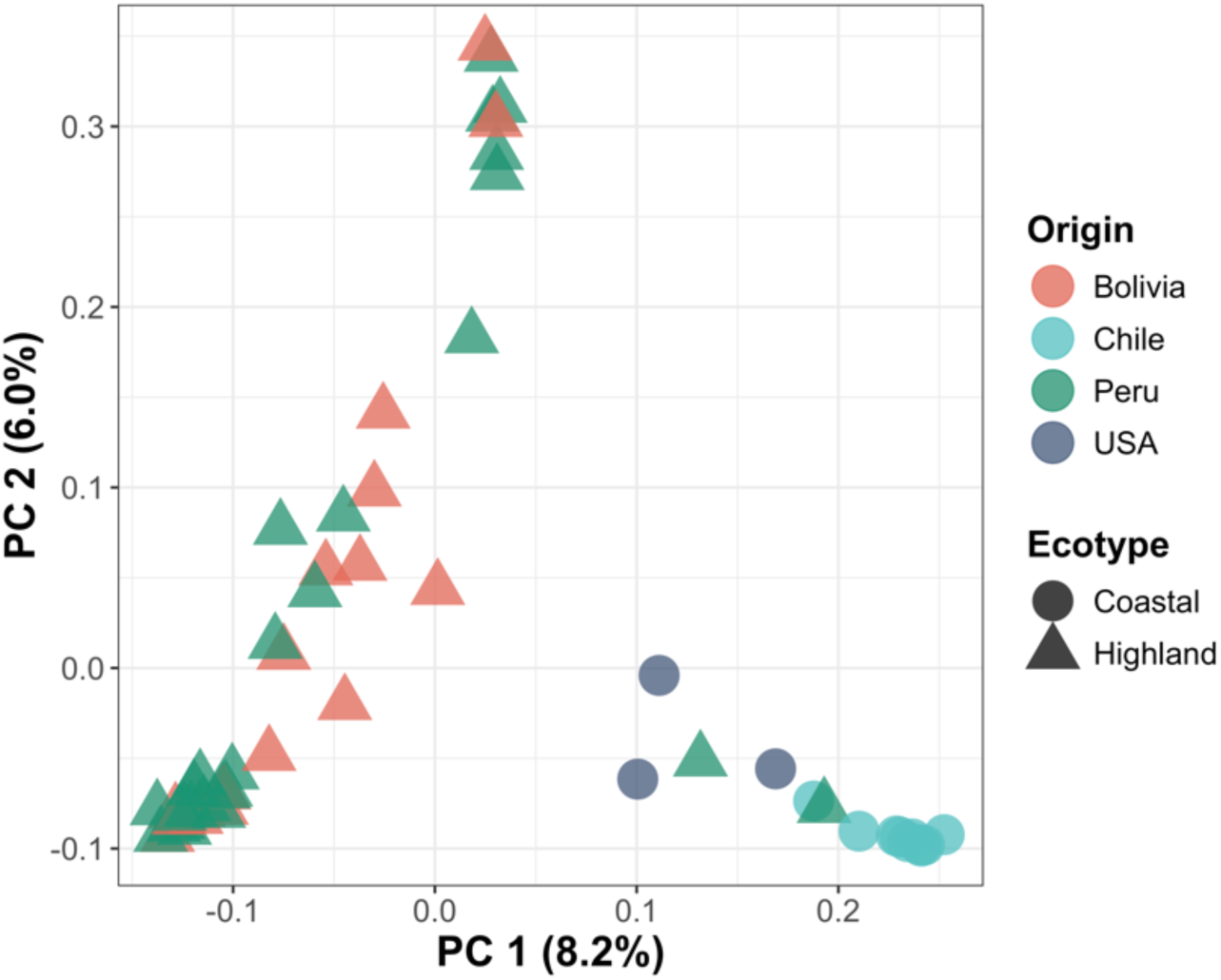
Population structure of quinoa accessions used in this study. A principal component analysis was conducted for 55 genebank accessions. The proportion of variance explained by Principal component 1 (PC1) is 8.2% and by PC2 is 6.0%. Colors represent the origin of the accessions based on passport data and symbols the ecotypes.

### Analysis of the field trial

#### Sowing date influences plant development and traits

To investigate the effects of low temperature on quinoa development, we conducted a single-year, single-location field experiment in which 60 accessions were sown at three different sowing dates: S1 (early March), S2 (early April), and S3 (late April). Mean soil temperatures during these sowing times were approximately 5 °C, 8 °C, and 11 °C, respectively (Table S1). Plants sown at S1 experienced pronounced cold stress, with an average minimum air temperature of 2.7 °C and 14 frost days during early growth. The experiment followed a randomized complete block design (RCBD) with three replicates. Within each replication, the 60 accessions were randomly assigned to 60 plots, each consisting of three adjacent rows corresponding to the three sowing dates. Nine phenotypic traits were recorded: mean emergence time, seedling emergence percentage, days to panicle emergence, flowering, maturity, plant height, panicle length, panicle width, and seed yield. A Wald F-test assessed the effects of genotype, sowing date, and replication on each trait. Both genotype and sowing date significantly affected all traits, and significant genotype × sowing date interactions were detected for days to panicle emergence, flowering, maturity, and panicle width (Table 2). Across all accessions, developmental stages were delayed under the earliest sowing condition. The mean growth cycle was longest for S1 (213 days), compared with 187 days for S2 and 163 days for S3. Similarly, seed emergence was markedly slower at S1 (26 days) than at S2 (15 days) and S3 (11 days), consistent with the lower soil temperatures at early sowing (Figure 2a).

**Figure 2.**
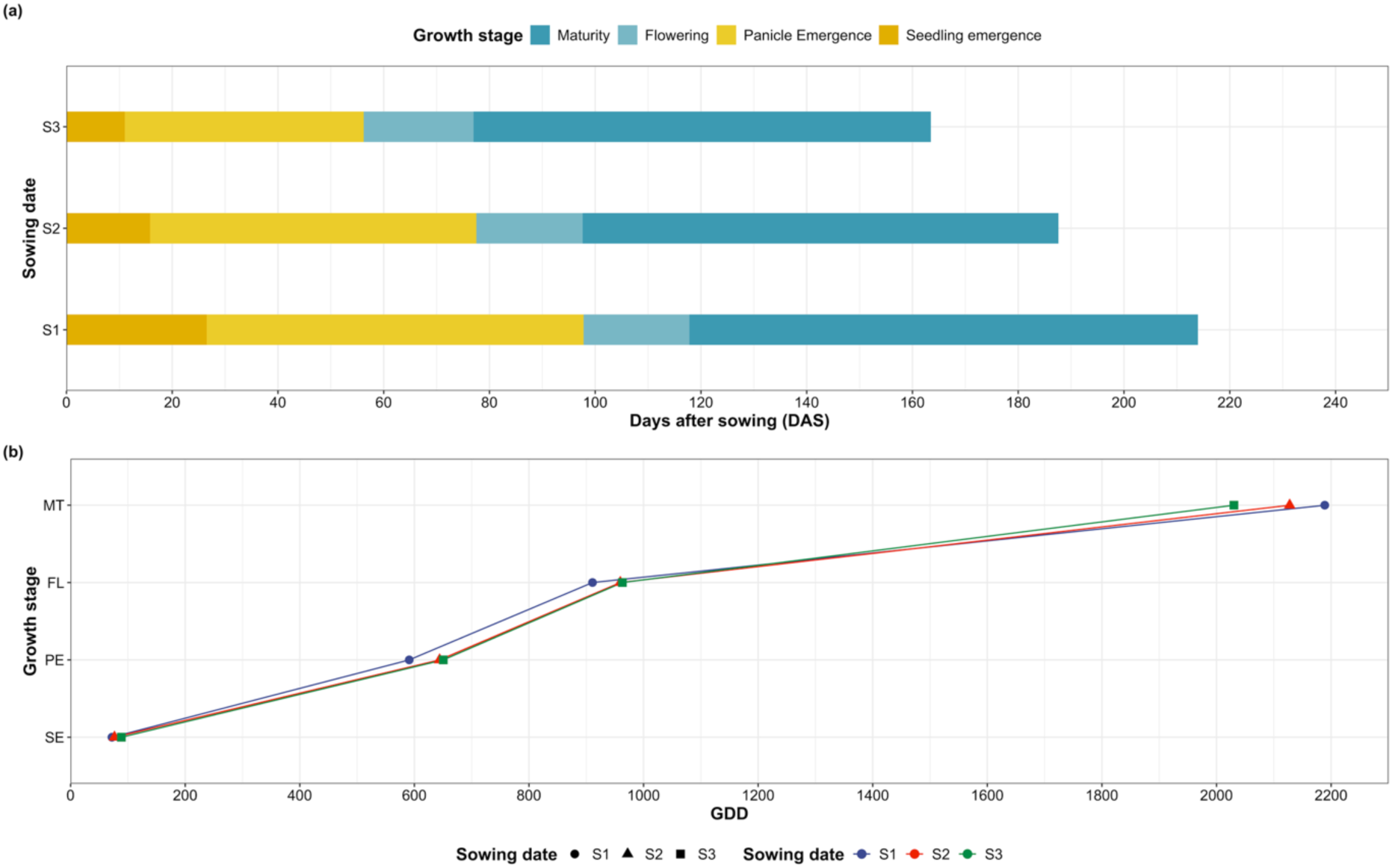
Comparison of mean growth stage duration and growing degree days for the three different sowing dates of the field experiment. (a) Mean growth stage duration in days after sowing (DAS) at sowing dates 1 (S1), 2 (S2), and 3 (S3). The x-axis represents days after sowing (DAS) and y-axis represents the three sowing dates. (b) Growing degree days (GDD) for the growth stages seedling emergence (SE), panicle emergence (PE), flowering (FL), and maturity (MT).

**Table 2.**
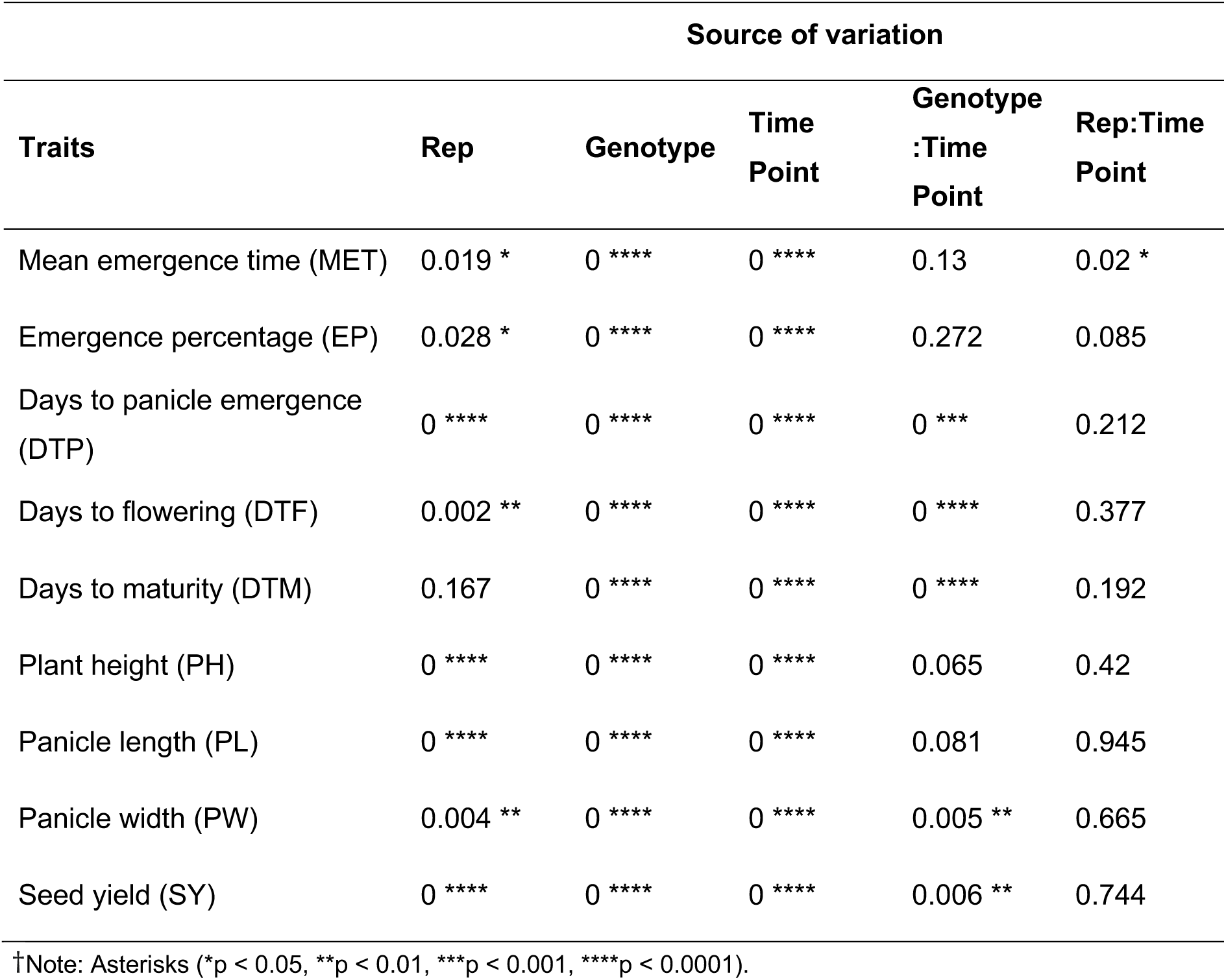
Wald F-test results for fixed effects in the field experiment evaluating the effects of low temperature conditions, reported as p-values.

Growing degree days (GDD) were calculated to quantify thermal time across developmental stages (Figure 2b) and to evaluate the influence of temperature accumulation on plant development. Despite differences in emergence timing, cumulative GDD at S1 was lower (72) than at S2 (76) and S3 (88). This trend persisted through panicle emergence and flowering, with S1 consistently accumulating fewer GDD than the later sowing dates. In contrast, plants sown at S1 required a higher cumulative GDD (2,188) to reach maturity compared with those at S3 (2,030), indicating that plants sown earlier experienced a prolonged growth period under cooler conditions. Although the duration of individual growth stages varied significantly, total GDD accumulation differed only slightly among sowing dates, suggesting that temperature accumulation exerted a stronger influence on developmental timing than sowing date per se (Figure 2b).

Mean trait comparisons across sowing dates revealed significant variation in phenotypic performance (Figure 3). Average emergence of seedlings remained below 50% across all treatments, with the highest emergence observed at S3 (nearly 40%) and the lowest at S1 (25%). Seed yield followed a similar trend: plants at S1 produced the highest mean yield per plot (64 g), whereas yields declined by approximately 30% at S2 (46 g) and 50% at S3 (35 g). Plant height also decreased progressively from S1 (165 cm) to S3 (149 cm), and a similar pattern could be observed with panicle length and width. Overall, both yield and key morphological traits declined consistently with later sowing dates, reflecting the cumulative effects of reduced thermal time and shortened growth duration.

**Figure 3.**
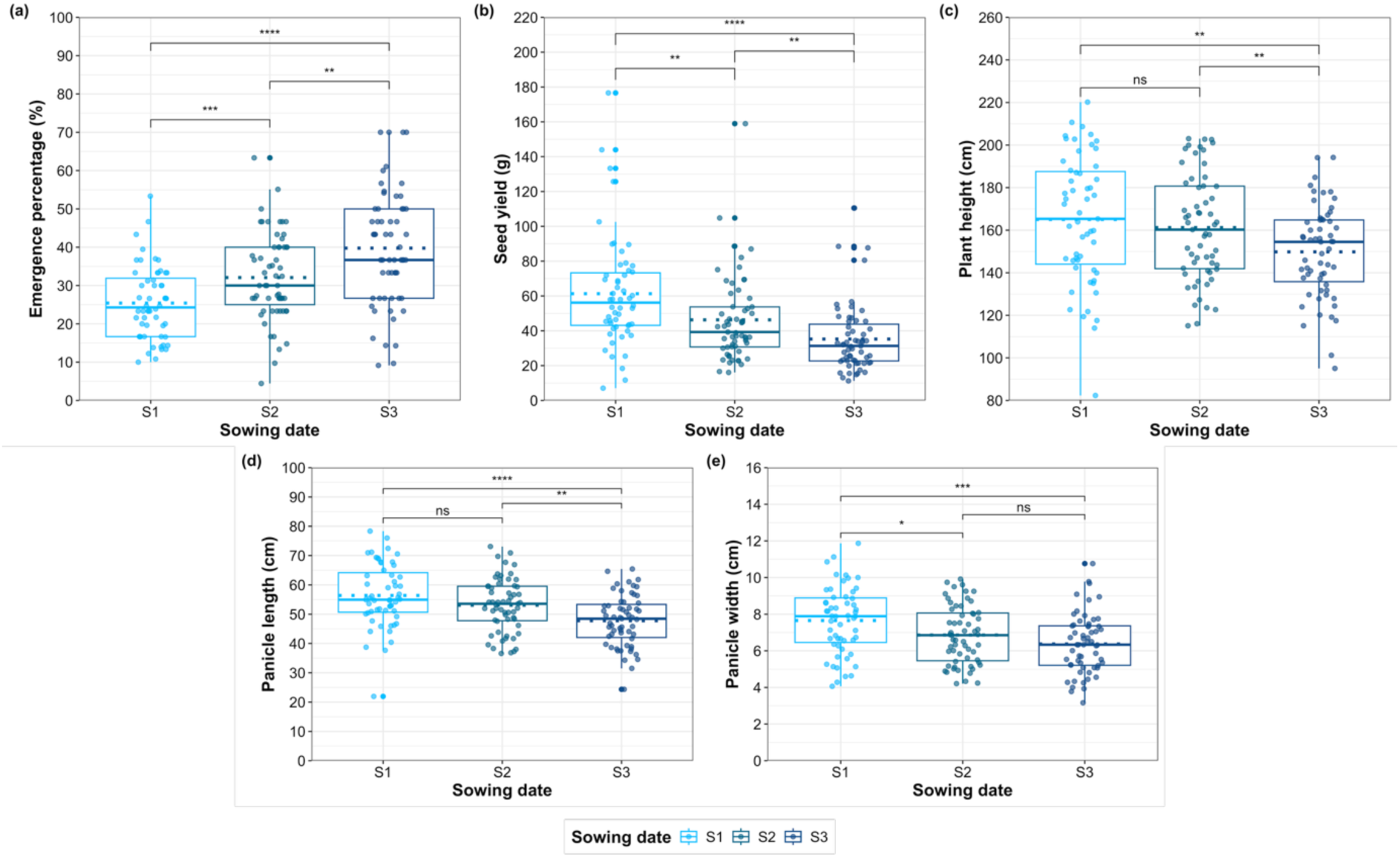
Best Linear Unbiased Estimated means (BLUEs) of 60 quinoa accessions for different traits across three sowing dates (S1, S2, and S3). The traits measured include (a) emergence percentage, (b) seed yield per plot (g), (c) plant height (cm), (d) panicle length (cm), and (e) panicle width (cm). The x-axis represents the sowing dates, while the y-axis indicates the respective measurement units for each trait. The dotted lines represents the mean. Statistical significance between sowing dates is indicated by asterisks (*p < 0.05, **p < 0.01, ***p < 0.001, ****p < 0.0001), and “ns” denotes non-significant differences.

#### Differential trait performance of coastal and highland ecotypes

Across all sowing dates, we observed phenotypic differences between coastal and highland groups of accessions (Figure 4). Significant variation occurred in S3 for emergence traits such as mean emergence time and emergence percentage (Figure 4a and 4b). At all sowing dates, highland accessions showed an earlier (i.e., quicker) emergence and had higher seedling emergence percentages compared to coastal accessions. Additionally, panicle emergence occurred later in coastal accessions than in highland ones across all sowing dates (Figure 4c), but days to flowering did not differ significantly between both groups across sowing dates (Figure 4d). In contrast, physiological maturity of coastal accessions occurred earlier than highland accessions, indicating a shorter overall growth period at all three sowing times (Figure 4e). Seed yield per plot also differed between ecotypes, with coastal accessions consistently producing higher yields across all sowing dates (Figure 4f). Plant height did not vary between the two groups at any sowing date (Figure 4g). In S2, coastal accessions developed shorter panicles compared to highland accessions (Figure 4h), but their panicles were wider (Figure 4i).

**Figure 4.**
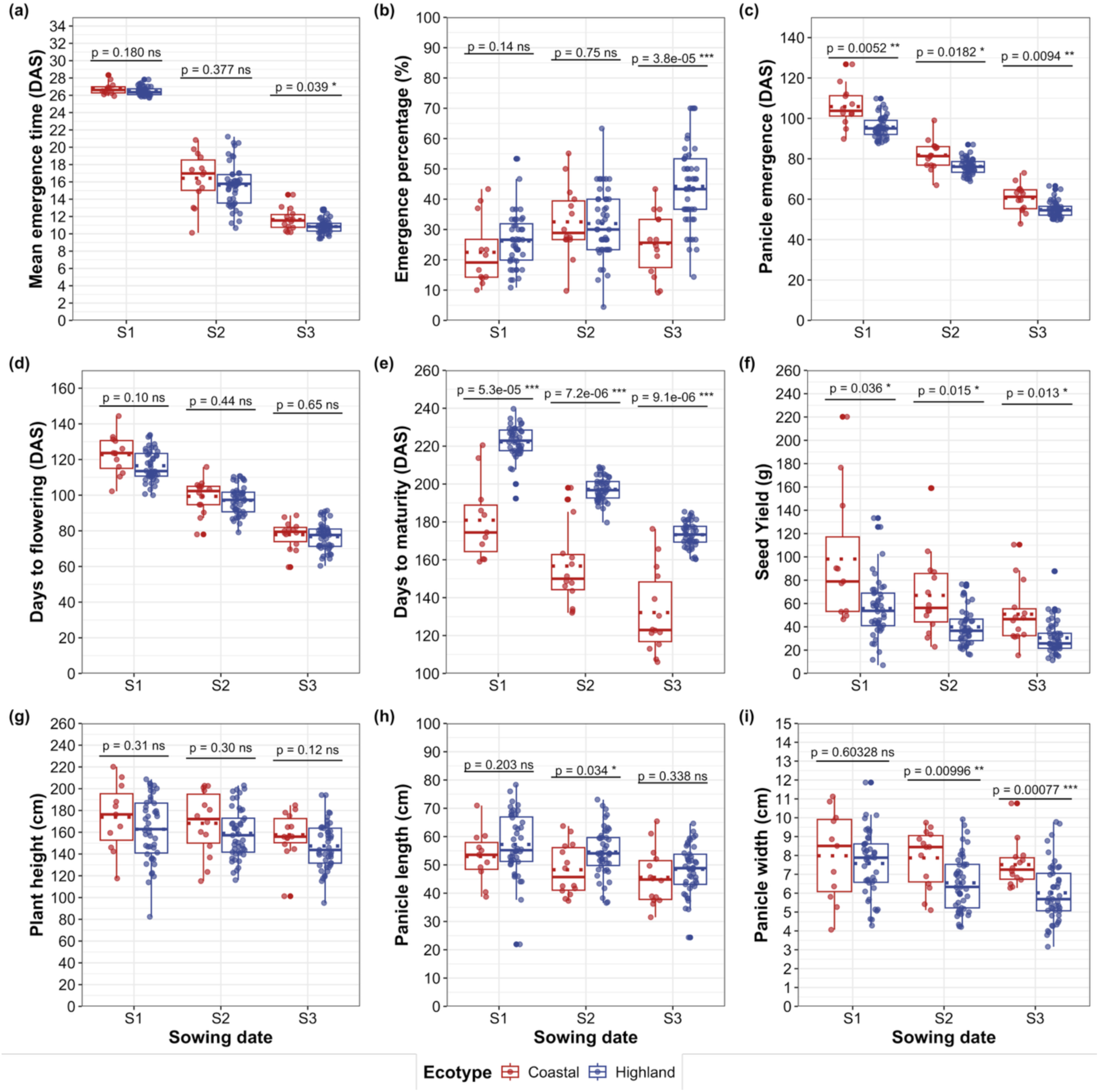
Phenotypic differences between coastal and highland quinoa ecotypes. Boxplots showing the estimated means (BLUEs) of 60 quinoa accessions from coastal (dark red) and highland ecotypes (blue) for different traits across three sowing dates (S1, S2, and S3). The traits measured include (a) mean emergence time (DAS) (b) emergence percentage, (c) panicle emergence (DAS), (d) days to flowering (DAS), (e) days to maturity (DAS), (f) seed yield per plot (g), plant height (cm), (h) panicle length, and (i) panicle width (cm). The x-axis represents the sowing dates, while the y-axis indicates the respective measurement units for each trait. The dotted lines represents the mean. Statistical significance between the ecotypes is indicated by p values (t-test) and asterisks (*p < 0.05, **p < 0.01, ***p < 0.001, ****p < 0.0001), and “ns” denotes non-significant differences.

We calculated the best linear unbiased estimates (BLUE) of 60 accessions from coastal and highland ecotypes to identify accessions that showed low seedling emergence and to compare their growth stages with respect to shorter growth cycles across different sowing dates (Figure S4). Coastal accessions consistently exhibited shorter growth periods than highland accessions across all sowing dates. Under cold stress conditions (S1), three coastal accessions (Ames-13751, PI-614888, and PI-634917) and one highland accession (CHEN-322) failed to germinate. Ames-13751 did not germinate under any sowing condition. In addition, Ames-13728 failed to mature in S1 and showed delayed maturity in the other sowing dates. Among the coastal accessions, Titicaca, PI-614886, and PI-634923 consistently exhibited shorter growth cycles. In contrast, no highland accession demonstrated consistently short growth durations, and all eventually reached maturity. Taken together, these observations highlight clear phenotypic differences between coastal and highland accessions, suggesting differential adaptation to environmental conditions.

#### Estimates of heritability and correlations between traits

To quantify the genetic contribution to phenotypic variation, broad-sense heritability (*H*²) was estimated following the method of Cullis et al., (2006) using Best Linear Unbiased Prediction (BLUP). Estimated *H*² values ranged from 0.36 to 0.95, with the lowest values observed for mean emergence time and seedling emergence percentage, and the highest for days to flowering and days to maturity (Table 3). The lower heritability of emergence traits indicates a stronger environmental influence compared with other phenological traits.

**Table 3.**
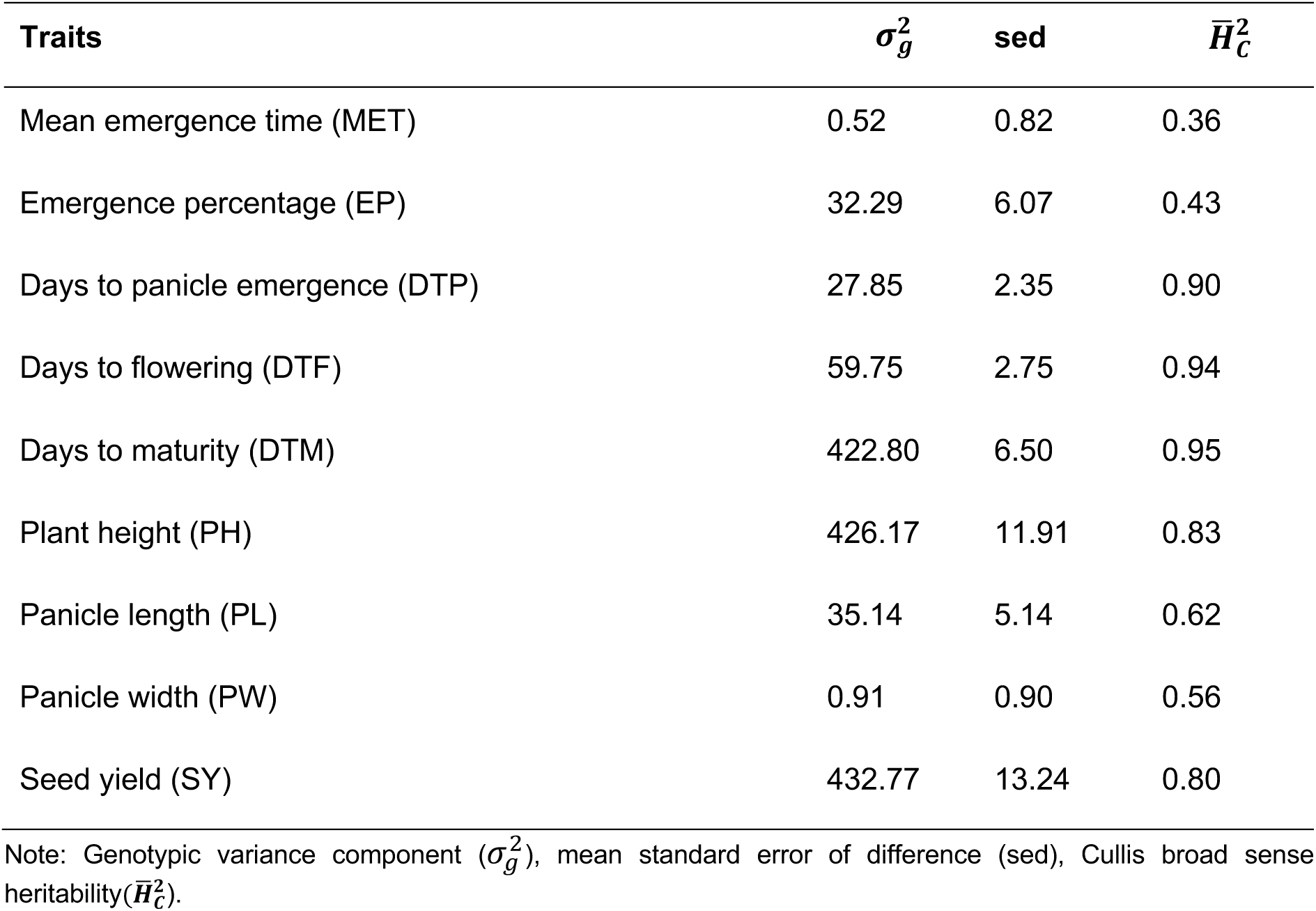
Genotypic variance, mean standard error difference, and heritability estimates for all traits.

To examine genetic relationships among traits, we calculated Pearson correlation coefficients for the nine measured variables (Figure S5). For both sowing dates S1 (Figure S5a) and S2 (Figure S5b), emergence traits showed no significant correlation with any other traits, consistent with their lower heritability and higher environmental sensitivity. Across all sowing dates, days to maturity were negatively correlated with seed yield, suggesting that earlier-maturing plants tend to produce higher yields. Plant height showed positive correlations with several traits, including days to panicle emergence, days to flowering, seed yield, panicle length, and panicle width. These associations indicate that plant height is influenced by both vegetative and reproductive development, and that taller plants generally exhibit superior physiological performance, resulting in greater seed yield.

#### Phenotypic principal component analysis (PCA) reveals variability among accessions

We performed a principal component analysis (PCA) using all phenotypic measurements to identify the variables that most strongly contribute to phenotypic variation (Rojas, 2003). The PCA showed that 81% of the total variation was represented by the first four principal components: PC1 = 34.3%, PC2 = 19.6%, PC3 = 17.7%, and PC4 = 9.2%. The first principal component (PC1) primarily distinguished accessions with later panicle emergence and flowering, but earlier maturity, greater plant height, compact panicles, and higher seed yield. This was reflected by the large positive coefficients for days to panicle emergence, plant height, days to flowering, seed yield, and panicle width, and a negative coefficient for days to maturity (Table S4). In the second principal component (PC2), days to maturity had the largest positive loading, representing accessions with late maturity. Panicle length contributed equally to both the second and third principal components. Emergence percentage contributed notably to both PC3 and PC4.

A biplot (Figure 5a) shows that most coastal accessions are characterized by high seed yield, dense panicles, and early maturity. In contrast, most highland accessions were lower yielding and later maturing. However, phenotypic data did not show strong clustering between coastal and highland accessions (Figures 5a and S6), in contrast to the clearer separation observed in the genetic diversity analysis (Figure 1).

**Figure 5.**
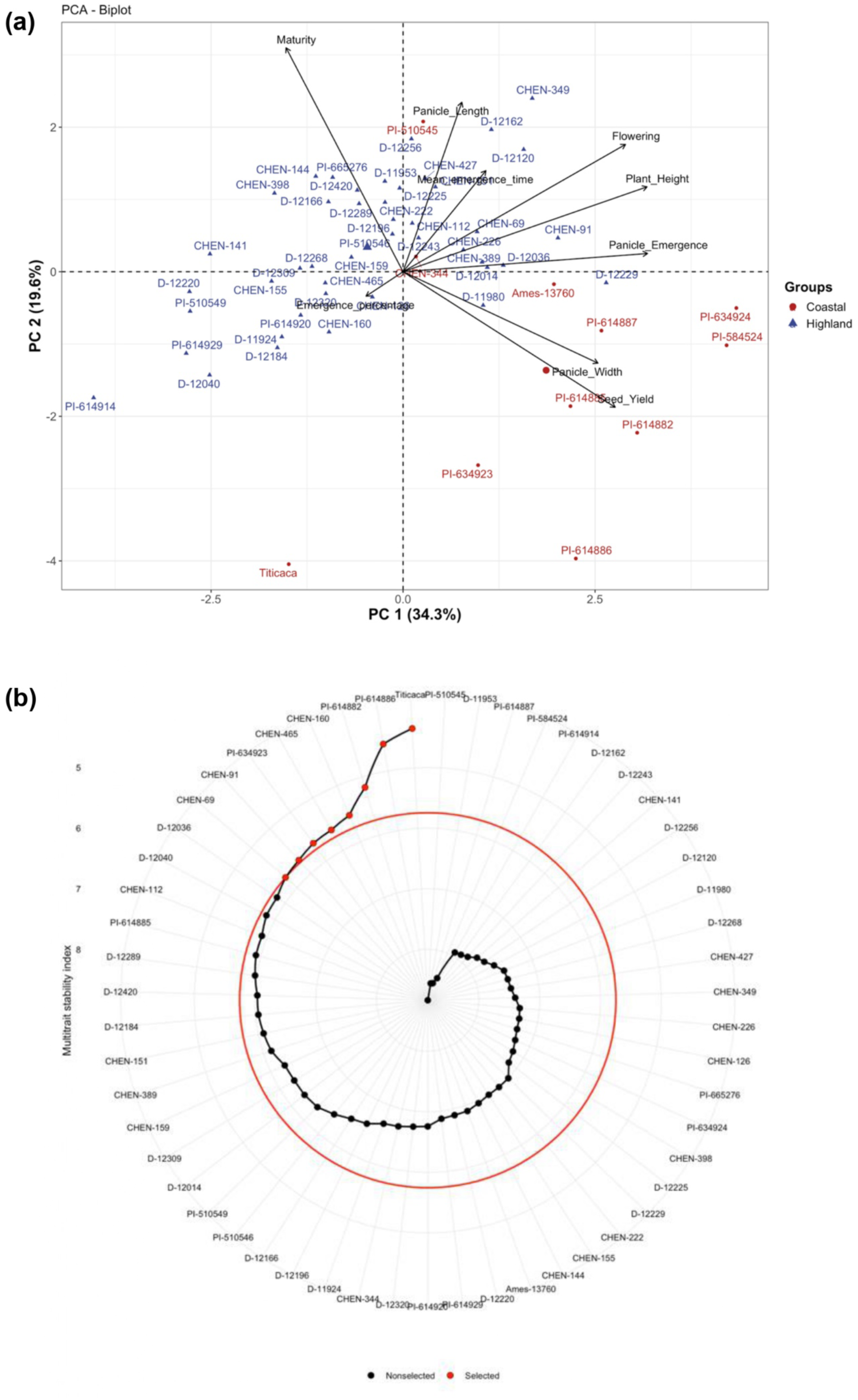
PCA of phenotypic data and multienvironmental stability analysis. a) PCA biplot of nine traits with first two principal components (PC1 = 34.3% and PC2 = 19.6%). The accessions were colored according to their ecotype coastal (dark red), highland (blue). b) Multi-trait stability index for 55 genotypes applied on mean emergence time (MET), emergence percentage (EP), days to flowering (DTF), days to maturity (DTM), plant height (PH), panicle width (PW), and seed yield (SY). The red cutoff circle indicates the cut point (5.7). The red points above the cutoff circle indicate genotypes which are stable across all the time points.

#### Multi-trait stability index identifies stable-performing accessions

To identify genotypes combining high mean performance with phenotypic stability, we computed the multi-trait stability index (MTSI; Olivoto et al., 2019 ) across all sowing dates for accessions that successfully germinated and reached maturity. In this study, the target quinoa ideotype was defined by high seedling emergence to ensure strong crop establishment, short growth cycles characterized by early emergence, flowering, and maturity, and compact, high-yielding plants of moderate height suitable for mechanical harvesting under European and Mediterranean conditions. Based on these criteria, we selected accessions that minimized mean emergence time, days to flowering, days to maturity, and plant height, while maximizing emergence percentage, panicle width, and seed yield.

Genotypes with the lowest MTSI values exhibited both high performance and stability across all sowing dates for the evaluated traits (Data S7). Applying a selection intensity of 15%, we considered genotypes with MTSI values below 5.7 as stable (Figure 5b). Among the 55 tested genotypes, eight met these criteria: four coastal accessions (Titicaca, PI-614886, PI-614882, and PI-634923) and four highland accessions (CHEN-160, CHEN-465, CHEN-69, and CHEN-91). For all analyzed traits, we report the mean of the selected genotypes (*Xₛ*), the population mean (*Xₒ*), and the selection differential (*Xₒ − Xₛ*) in Table S5. Negative selection differentials for mean emergence time, days to flowering, days to maturity, and plant height, together with positive differentials for emergence percentage and seed yield, demonstrate that the MTSI approach effectively identified genotypes matching the desired ideotype. The selected accessions exhibited earlier flowering and maturity, greater seed yield, and wider panicles, while maintaining strong emergence and optimal plant height (Table S6).

### Evaluation of quinoa seed germination traits under controlled conditions

Considering that differences in germination observed in our field experiment could have been influenced by factors other than low temperatures, such as humidity, soil characteristics, day length, or seed predation, we conducted a controlled laboratory experiment to isolate the effect of temperature. Germination was assessed under cold conditions (4.4 °C) and standard conditions (18.3 °C). In this experiment, we documented germination parameters using both manual measurements and a newly developed deep learning pipeline for automated analysis of images of seeds and seedlings at different time points throughout the experiment. The data obtained from manual and automated measurements were analyzed independently and then compared to evaluate the reliability and usefulness of the deep learning approach. Using both methods, we measured germination percentage and mean germination time. Additionally, we assessed viable germination percentage, which is defined as the percentage of seedlings that developed properly (i.e., with visible roots, shoots, and leaves) during manual scoring. Growth rates were also calculated based on seedling surface area, as extracted by the deep learning pipeline from the image data. A detailed description of the data generation workflow and the image analysis pipeline, which is based on a Mask R-CNN, is provided in the supplementary text.

#### Genotype and cold treatment affect seed germination under laboratory conditions

ANOVA performed on the laboratory germination data revealed effects of genotype, treatment, and their interaction on mean germination time, viable germination percentage, and growth rate (p<0.01; Table S7). Germination percentage, however, was affected only by genotype and the genotype x treatment interaction (p<0.01; Table S7). Estimated means (BLUEs) were calculated for all traits, assuming fixed genotypic effects, and used to test for mean differences between treatments and between coastal and highland ecotypes. Manual measurements of germination parameters showed that the average germination time under low-temperature conditions was longer (mean: 7.11 days after sowing [DAS]) compared to control conditions (mean: 1.46 DAS, p<0.0001; Figure 6a). Although final germination percentages did not differ between the two treatments (Figure 6b), the germination percentage of viable seedlings differed markedly (p<0.001; Figure 6c and 6e), with low temperature conditions resulting in a lower proportion of viable seedlings than room temperature conditions. Results from the image analysis also showed that low temperatures reduced both mean germination time and seedling growth rate (p<0.0001; Figure 6d and 6f).

**Figure 6.**
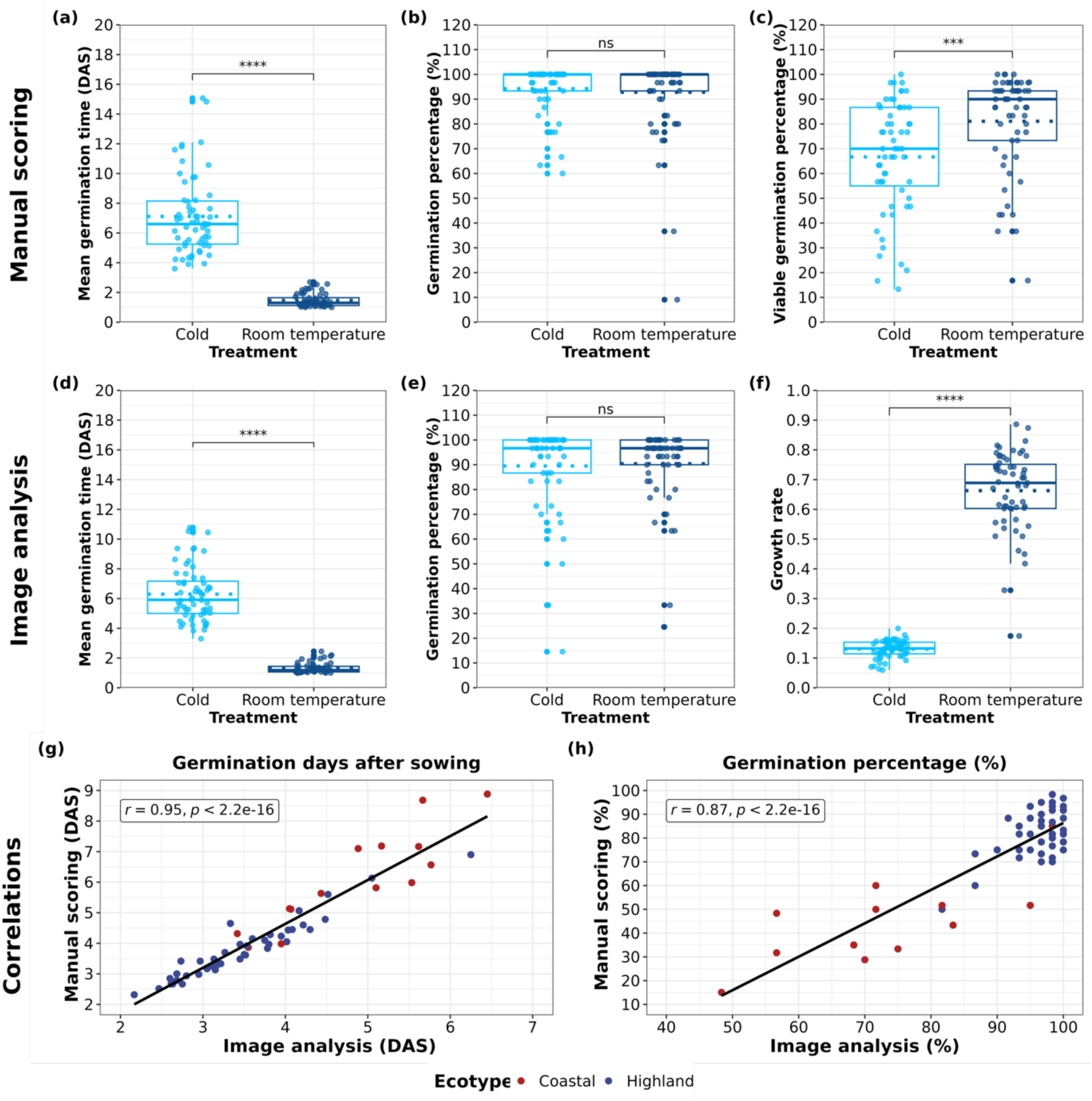
Differences of estimated means (BLUEs) in the laboratory assay of 60 quinoa accessions under cold and room temperature treatments. First row consists of results from manual scoring (a) mean germination time (DAS), (b) germination percentage (%), (c) viable germination percentage (%). Second row consists of results from image analysis (d) mean germination time (DAS), (e) germination percentage (%), (f) growth rate. The x-axis represents the treatments, while the y-axis indicates the respective measurement units for each trait. The dotted lines represents the mean. Statistical significance between the treatments based on a t-test is indicated by asterisks (*p < 0.05, **p < 0.01, ***p < 0.001, ****p < 0.0001), and ”ns” denotes non-significant differences. (g) and (h) Correlation plots between manual scoring (y-axis) and image analysis data (x-axis) for the trait (a) germination days after sowing (DAS) and (b) germination percentage (%). The *r* values are Pearson correlation coefficients and a p-value <0.05 correlation is considered significant. Accessions were colored by their assignment to the coastal (dark red) or highland (blue) ecotypes.

#### Correlations between the manual scoring and image analysis

Due to the tedious nature of manual scoring, automated image analysis can serve as a useful tool for these types of germination tests. To assess the extent of correlation between the outcomes of the two methods, we correlated the BLUE values for the traits mean germination time and germination percentage. For both traits we obtained strongly positive correlations (Pearson correlation, *r*=0.95 and *r*=0.87, respectively, p<0.0001; Figure 6g and 6h) supporting the accuracy and effectiveness of our novel image analysis approach.

#### Differences between quinoa ecotypes in germination parameters

Under low temperature conditions conditions, highland quinoa exhibited markedly superior germination performance compared with coastal accessions (all *P* < 0.01 or smaller; Table 4). Highland accessions achieved an average germination percentage of 98.6% and a viable germination percentage of 75.4%, whereas coastal accessions reached only 80.2% and 38.9%, respectively. Germination also occurred earlier in highland quinoa, with a mean germination time of 6.2 days after sowing (DAS), compared with 10.2 DAS in coastal accessions. Even under control conditions, highland accessions showed faster and more complete germination than coastal accessions. Image-based analyses confirmed manual assessments by finding consistent differences in germination performance between the two ecotypes (Table 4). Growth rate also differed significantly between groups: under low temperature conditions, highland accessions grew faster (0.14) than coastal accessions (0.10), and this pattern persisted under control conditions (0.69 vs. 0.58; Table 4).

**Table 4.**
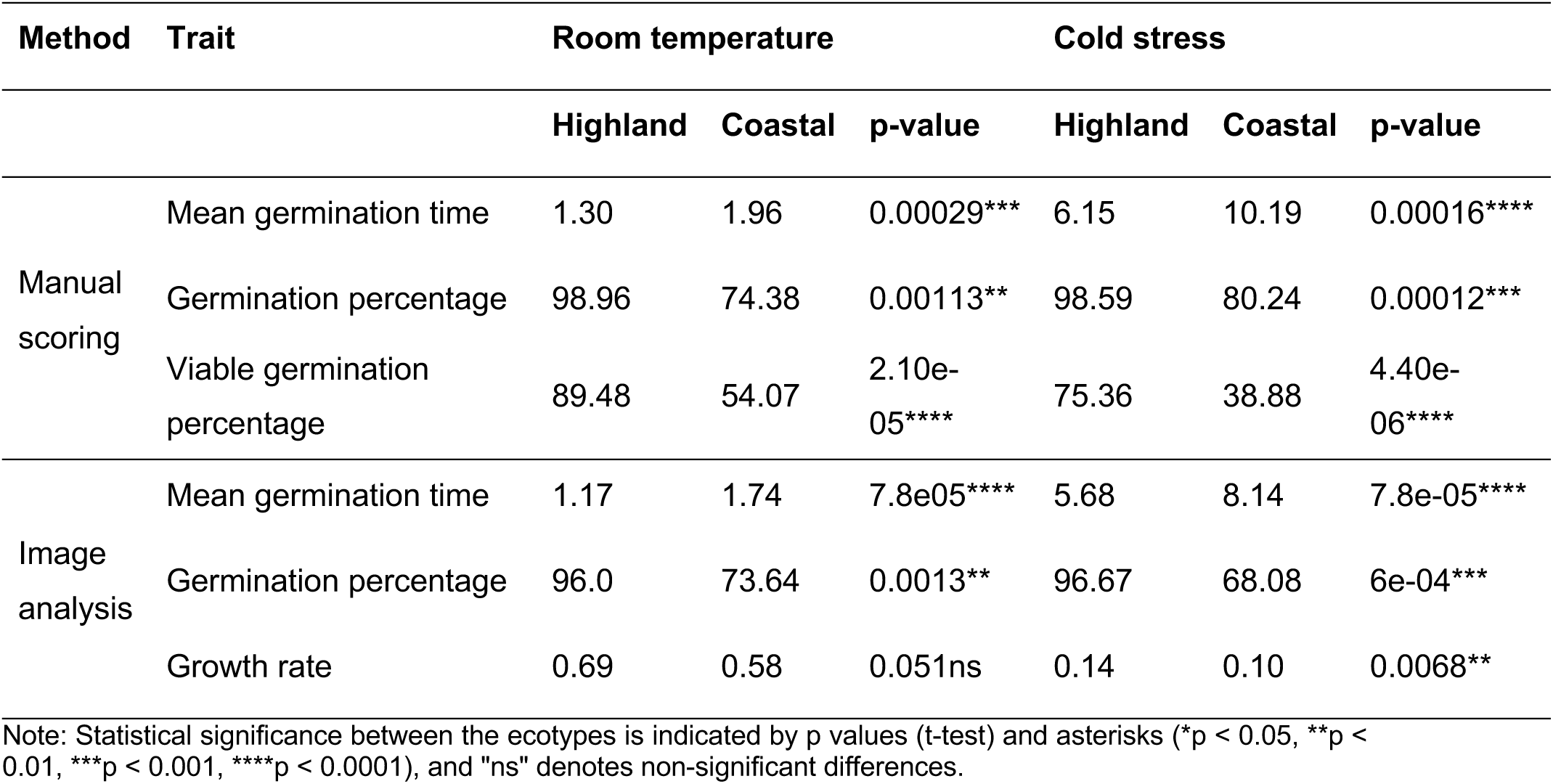
Mean and p-values (t-test) of germination traits in comparison between coastal and highland ecotypes within each treatment in the laboratory experiment.

| Method | Trait | Room temperature |  |  | Cold stress |  |  |
| --- | --- | --- | --- | --- | --- | --- | --- |
|  |  | Highland | Coastal | p-value | Highland | Coastal | p-value |
| Manual scoring | Mean germination time | 1.30 | 1.96 | 0.00029*** | 6.15 | 10.19 | 0.00016**** |
|  | Germination percentage | 98.96 | 74.38 | 0.00113** | 98.59 | 80.24 | 0.00012*** |
|  | Viable germination percentage | 89.48 | 54.07 | 2.10e-05**** | 75.36 | 38.88 | 4.40e-06**** |
| Image analysis | Mean germination time | 1.17 | 1.74 | 7.8e05**** | 5.68 | 8.14 | 7.8e-05**** |
|  | Germination percentage | 96.0 | 73.64 | 0.0013** | 96.67 | 68.08 | 6e-04*** |
|  | Growth rate | 0.69 | 0.58 | 0.051ns | 0.14 | 0.10 | 0.0068** |

Note: Statistical significance between the ecotypes is indicated by p values (t-test) and asterisks (\*p < 0.05, \*\*p < 0.01, \*\*\*p < 0.001, \*\*\*\*p < 0.0001), and "ns" denotes non-significant differences.

### Seed size differs between highland and coastal accessions

Seed size was assessed for 59 of the 60 accessions using two complementary approaches: the MARViN Seed Analyzer System and an automated image analysis pipeline based on a Mask R-CNN model. MARViN measurements were obtained for seeds from three successive generations with *G₀* (original), *G₁* (offspring of *G₀*), and *G₂* (offspring of *G₁*) to evaluate the heritability of seed size. Data were standardized by calculating relative values across generations. Based on MARViN measurements (Figure 7a), highland accessions produced smaller seeds than coastal accessions (*t*-test; p<0.05). Accessions were also classified as bitter (29) or sweet (30) according to saponin presence, revealing that bitter accessions had larger seeds than sweet ones (Figure 7b, p<0.0001). The heritability estimate for relative seed size was 0.89, indicating a strong genetic component for this trait.

**Figure 7.**
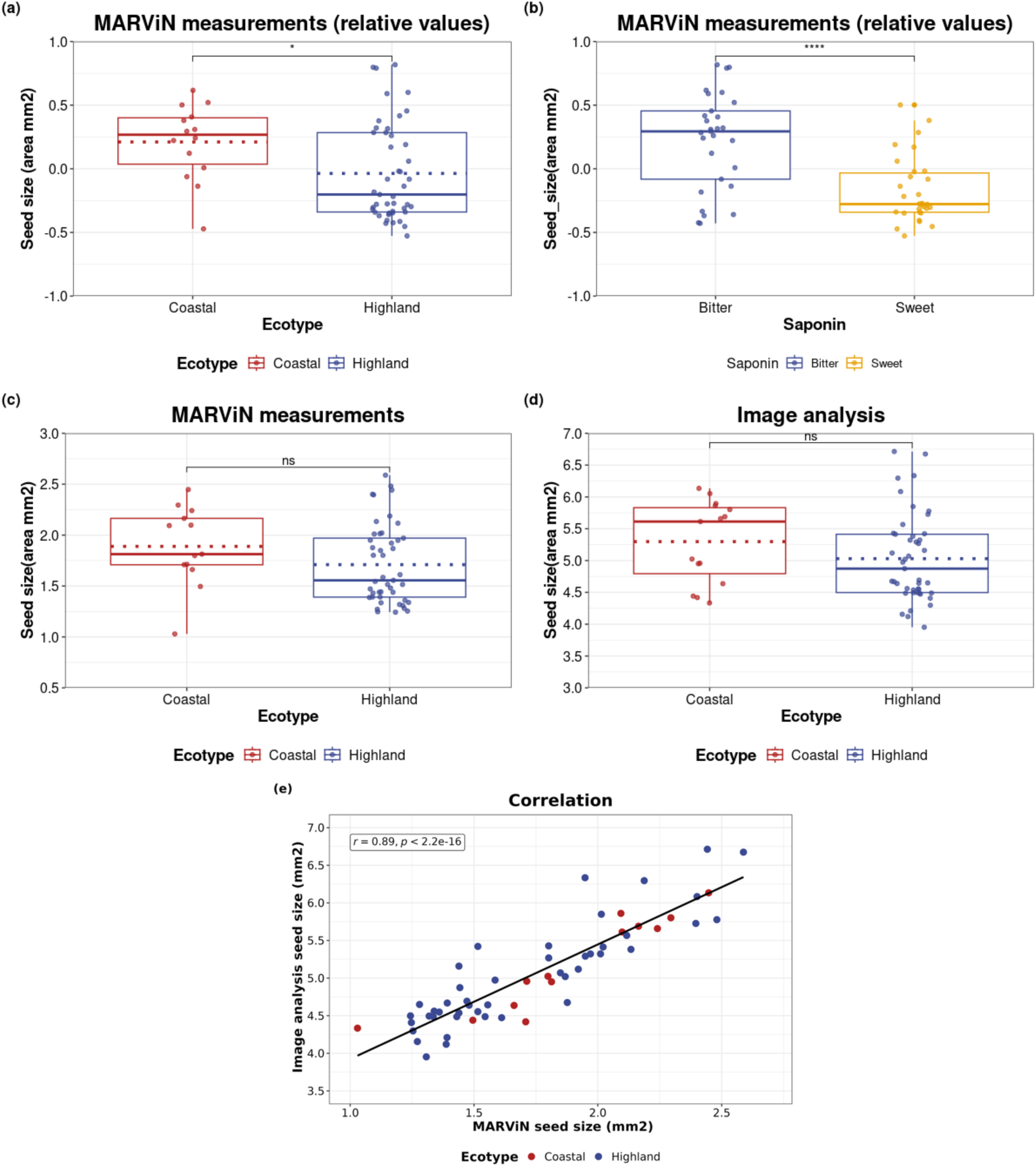
Estimated means (BLUEs) of quinoa accessions for the trait seed size. (a) seed size relative values from MARViN measurements of accessions from highland and coastal ecotypes (b) seed size relative values from MARViN measurements of bitter and sweet (saponin) accessions (c) normal values of MARViN measurements (generation 1) (d) seed size measurements from image analysis of accessions from highland and coastal ecotypes. X-axis represents different ecotypes, and the y-axis represents the seed size area in mm2. Statistical significance between the treatments based on a t-test is indicated by asterisks (*p < 0.05, **p < 0.01, ***p < 0.001, ****p < 0.0001), and ”ns” denotes non-significant differences. (e) Correlation plots between MARViN measurements and image analysis data for seed size. The *r* values are the pearson correlation coefficient value and a p-value <0.05 correlation is considered significant. The coastal accessions are represented in dark red, and the highland accessions are represented in blue colors.

Because MARViN measurements are time-intensive, we evaluated automated image analysis as a high-throughput alternative. Seed surface area was extracted from images captured on day 1 of the laboratory germination experiment. Comparisons between image-based and MARViN measurements (G₁ generation) showed no significant differences between the two methods. Both approaches consistently indicated that highland accessions tend to have smaller seeds, although the difference was not statistically significant (Figures 7c,d). The two measurement methods were strongly correlated (Pearson’s *r* = 0.89, p<0.0001; Figure 7e) and yielded similar heritability estimates for seed size (0.88 for MARViN and 0.94 for image analysis), confirming substantial genetic variation in this trait.

To examine the relationship between seed size and germination performance, we calculated Pearson correlation coefficients using image-derived data (Figure S7). Seed size was negatively correlated with growth rate (*r* = –0.69, p<0.001), indicating that smaller seeds are associated with faster seedling growth.

## DISCUSSION

Fast and resilient seed germination is a crucial step for a crop establishment, plant growth and eventually the final yield (Suo et al., 2023). Low temperatures is one of the major abiotic stresses which adversely influences germination, plant subsequent growth and phenological processes (Bhattacharya, 2022). In this study we investigated seedling emergence and phenotypic differences among 60 accessions of highland coastal quinoa under field conditions together with their germination patterns under laboratory conditions. Our analysis of quinoa seedling emergence under field conditions revealed that low temperatures during early crop stages delayed emergence and led to lower seedling emergence percentages compared to seeds sown under optimum temperature conditions. Highland accessions showed earlier seedling emergence and achieved higher emergence percentages, whereas coastal accessions matured earlier and produced higher seed yields. These field observations were supported by the laboratory experiment, which confirmed the negative impact of cold stress on quinoa seed germination.

### Interaction of sowing date and cold stress on trait expression under field conditions

In the field experiment, 60 quinoa accessions were sown at three different dates to evaluate the effects of low temperature on trait expression, particularly seedling emergence. The earliest sowing (S1, early March) occurred under cold conditions, with mean (5.6 °C) and minimum soil temperatures (2.7 °C) below the optimal range of 8–10 °C for quinoa germination (Jacobsen & Stølen, 1996; Table S1). Subsequent sowings in early April (S2) and late April (S3) represented more favorable temperature conditions, with mean soil temperatures of 8 °C and 10 °C, respectively. In S1, seedling emergence was delayed by 10–15 days, and emergence percentages were reduced by 7% and 15% relative to S2 and S3, respectively. Similar temperature-dependent delays and reductions in emergence have been documented in previous field and greenhouse studies (Bois et al., 2006; Hirich et al., 2014; Jacobsen & Stølen, 1996), which also noted substantial impairment of quinoa germination at suboptimal temperatures. These results indicate that the delayed emergence and lower establishment observed in our study were directly caused by the low soil temperatures experienced during early sowing.

Low temperatures following early sowing are also known to extend the vegetative phase of quinoa (Risi & Galwey, 1991), a trend reflected in our results: plants sown in S1 required more days to reach flowering compared with those in S2 and S3. Conversely, plants in S2 and S3 matured earlier, with the shortest growth duration observed in S3 (Figure 2a). These observations support previous findings that increasing temperature accelerates quinoa development (Hirich et al., 2014). Although the total growth period was longest for S1, flowering and maturity occurred at similar calendar dates across all sowing dates. This can be attributed to the accumulation of comparable growing degree days (GDD) at key developmental stages (Figure 2b). Rising temperatures beginning in June allowed all plants to accumulate sufficient thermal time, resulting in synchronized flowering and maturity.

The shortened vegetative period (from days to panicle emergence to days to flowering) in later sowings likely restricted vegetative growth and contributed to the reduced plant height observed in S3, consistent with the findings of Temel & Yolcu, (2020). Despite lower emergence percentages, plants sown in S1 achieved the highest seed yield. This pattern supports earlier reports that early-sown quinoa can effectively exploit the full growing season, even under reduced plant densities (Risi & Galwey, 1991). Consequently, although early sowing limited establishment, it enhanced overall productivity, yielding the highest total seed output among the three sowing dates. The differences in realized plant density among sowing dates may have contributed to variation in later growth and yield traits. Nevertheless, quinoa exhibits substantial phenotypic plasticity and compensatory responses to variation in plant density, which may buffer part of these effects (J. A. González et al., 2022).

Together, these findings demonstrate that temperature at sowing strongly influences early development and yield formation in quinoa. The clear differences in emergence dynamics, phenological timing, and productivity across sowing dates highlight the sensitivity of this crop to low temperature conditions during establishment. Since cold stress effects are confounded with the differences in season length and GDD, therefore the observed differences might also likely reflect combination of these factors rather than the effect of temperature alone.

### Ecotypic variation in phenotypic plasticity in response to cold stress

We observed that under low temperature conditions, highland accessions emerged earlier and achieved higher emergence percentages than coastal accessions (Figures 4a,b), reflecting their adaptation to high-altitude environments characterized by lower and more variable temperatures (Jacobsen et al., 2007; Le Tacon et al., 1992). Our findings are consistent with those of Jacobsen et al., (2005), who reported greater cold sensitivity in coastal compared with highland types. Three coastal accessions and one highland accession (Ames-13751, PI-614888, PI-634917, and CHEN-322) failed to germinate under cold stress, further supporting this pattern. The superior germination performance of highland accessions under cold conditions may reflect differences in seed vigor and reserve mobilization efficiency during early seedling development. Studies in oilseed rape *Brassica napus* (Nykiforuk & Johnson-Flanagan, 1999), and quinoa (Rosa et al., 2009; Yurin et al., 2026) suggested that low temperatures primarily delay metabolic processes associated with reserve mobilization and early growth. Low temperatures may not prevent germination itself (e.g., Bois et al., 2006) and genotypes with greater vigor may therefore establish more rapidly under suboptimal temperatures. The faster emergence and higher growth rates observed in highland accessions indicates adaptation to high-altitude environments, where rapid establishment under short growing seasons and fluctuating temperatures is likely a strongly selected trait.

Although coastal accessions, on average, reached panicle emergence and flowering later than highland accessions, they showed the shortest time span from flowering to maturity the three sowing dates (Figures 4c–e), which is a typical characteristic of Chilean coastal types (Risi & Galwey, 1989). Consequently, coastal accessions had a shorter total growth period than highland accessions across all sowing dates (Figure S4) and consistently produced higher seed yields (Figure 4f). Their superior performance under temperate conditions has been demonstrated in several European field trials (Jacobsen, 1997; Risi & Galwey, 1991), which explains why most European quinoa cultivars have been derived from coastal germplasm. The contrasting pattern of stronger early establishment in highland accessions but higher seed yield and shorter growth cycles in coastal accessions suggest a trade-off between early vigor and life-cycle duration. Selection in high-altitude environments may favor rapid establishment and stress resilience during early development, whereas coastal germplasm may allocate resources toward accelerated reproductive development and greater yield potential under temperate conditions. Similar life-history trade-offs between establishment and reproductive investment have been proposed for crops adapted to contrasting environments (Lundgren & Marais, 2020). Interestingly, because highland accessions tended to possess smaller seeds despite faster growth rates, differences in cold adaptation are unlikely to be explained solely by seed size and may instead reflect physiological differences in seed metabolism or reserve utilization efficiency. Future breeding programs could attempt to mitigate the trade-off and utilize the complementary strengths of both ecotypes to develop resilient, high-yielding quinoa varieties adapted to diverse and variable climatic conditions. The best-performing accessions identified in this study represent promising candidates for such breeding efforts.

### Heritability, trait correlations, and multi-trait stability analysis

We observed low heritability estimates for mean emergence time and emergence percentage, indicating that these traits are strongly influenced by environmental factors. In contrast, high heritability estimates were obtained for days to panicle emergence, flowering, maturity, plant height, and seed yield, suggesting a stronger genetic contribution to variation in these traits (Table 3). These results are consistent with those of Bhargava et al., (2007), who also reported substantial genetic control over similar phenological and yield-related traits.

Pearson correlation analyses (Figure S5) revealed no significant associations involving emergence traits, suggesting that these characteristics exert limited influence on later growth stages or morphological development. A consistent positive correlation was observed between days to panicle emergence and days to flowering across all sowing dates. Plant height showed weak positive correlations with both traits, indicating that earlier growth phases may moderately affect final plant stature. Moreover, plant height showed positive correlations with panicle length and width, suggesting that taller accessions tend to produce longer and more compact panicles, as previously noted (Risi & Galwey, 1989; Rojas, 2003). Finally, a significant negative correlation between seed yield and days to maturity supports the conclusion that early-maturing accessions tend to achieve higher yields.

### Principal component analysis of genetic and phenotypic variation

The initial classification of accessions into ecotypes based on passport data provided a useful first approximation of their geographical origins. The genotypic principal component analysis (PCA; Figure 1) confirmed this grouping and are consistent with previous studies identifying two major quinoa lineages: highland accessions from Bolivia and Peru, and coastal accessions from Chile and the United States (Patiranage et al., 2022; Zhang et al., 2017). Two highland accessions clustered with coastal types, likely reflecting seed exchange or inaccuracies in passport records, which sometimes list collection centers rather than true sampling locations. Phenotypic analyses (Figures 5a and S3) did not show clear ecotypic differentiation for traits such as emergence percentage, mean emergence time, or panicle length. However, the first two principal components distinguished accessions by key agronomic traits, particularly high yield and early maturity, indicating substantial genetic variation within the panel.

The multi-trait stability index (MTSI) further identified eight accessions, comprising four coastal and four highland accessions, with stable and superior performance across sowing dates (Figure 5b; Table S4). These accessions combined desirable traits such as high emergence, early maturity, short stature, compact panicles, and high seed yield. The European cultivar *Titicaca* of coastal ancestry confirmed its strong adaptation to Central European conditions, while coastal accession PI-614886, the reference genotype for the quinoa genome (Jarvis et al., 2017), represents a valuable resource for future studies. Among highland accessions, CHEN-160 exhibited a short growth cycle and CHEN-91 produced the highest yields. Because the stability analysis treated the three sowing dates as separate environments and was conducted at a single site, results should be interpreted cautiously. Broader multi-environment trials may uncover additional genotype-by-environment (G×E) interactions and refine current assessments of trait stability (Teressa et al., 2021).

### Consistency between field and laboratory experiments

The consistency of germination traits across both field and laboratory experiments provides strong evidence for differential adaptation between the two main quinoa ecotypes. Importantly, while the field experiment reflects the response to multifactorial early-season environmental conditions, the controlled laboratory experiment specifically isolated temperature effects and independently confirmed the superior germination performance of highland accessions under low-temperature conditions. The agreement between both experiments therefore supports our that temperature contributes substantially to ecotypic differences in germination but also acknowledges that field responses emerge from multiple interacting environmental factors. Although the field trial was conducted in a single year and location only, the inclusion of three sowing dates within one growing season offers valuable internal replication and allowed to identify growing degree days as a key component determining quinoa phenology. The complementary experiment under controlled laboratory conditions further strengthens the robustness and generalizability of our results on seed and germination traits. The strong agreement between field and laboratory results suggests that high-throughput germination experiments can be efficiently implemented under controlled conditions using standardized imaging protocols. This approach enables efficient screening of extensive genebank collections and segregating breeding populations to identify genetic variation for favorable germination traits from highland accessions that may be introgressed into early-maturing, high-yielding coastal genotypes. Overall, the consistency between laboratory and field observations suggests that germination traits measured under controlled conditions may partly predict field establishment under early-season conditions. Interestingly, the superior germination and emergence performance of highland accessions did not translate into higher yield, indicating a potential trade-off between early vigor and later reproductive performance. Similar genotype-dependent relationships between early vigor and field performance have been reported previously in quinoa (Ahmadi et al., 2026).

### Implications of image analysis using deep learning for quinoa research and breeding

A key innovation of this study was the application of a deep learning–based image analysis pipeline to measure seed size and germination traits, thereby enabling efficient processing of large sample sets and the generation of robust phenotypic data. Deep learning methods are increasingly used in crop science for precision agriculture, yield prediction, and automated phenotyping (Jabed & Murad, 2024; Upadhyay et al., 2025; Vithlani & Dabhi, 2023). Using our automated setup, we achieved 95.1% detection accuracy for seeds and seedlings and 100% accuracy for QR-code and ruler elements—values comparable to or exceeding those reported for other crops (Fan et al., 2023; Genze et al., 2020; Nehoshtan et al., 2021; Zhao et al., 2023). Despite the challenges of detecting small objects such as seeds, our approach reliably analyzed approximately 7,000 images from 60 accessions within a short time frame. This high-throughput capability accelerates phenotyping and facilitates the rapid identification of desirable genotypes, such as those with enhanced cold tolerance. Automated scores showed strong agreement with manual records (Figures 6g,h), confirming the accuracy of detection. The pipeline reduced manual workload by about 90% while improving reproducibility and expanding trait coverage, which enables large-scale genetic analyses such as QTL mapping of germination traits.

Model optimization further improved performance: training the full ResNet-101 backbone enhanced detection precision, and emphasizing object classification over segmentation yielded better accuracy, likely due to the difficulty of distinguishing germinated from non-germinated seeds. Unlike previous approaches, we systematically optimized and reported Mask R-CNN hyper-parameters, providing a framework for future high-throughput phenotyping applications (see Supplementary Text). These results demonstrate the potential of deep learning–based phenotyping to substantially enhance the precision, scale, and efficiency of quinoa improvement and modern crop breeding programs. However, the reduced performance observed in external validation experiments and the need for image filtering indicate that the current implementation remains dependent on the specific imaging setup and may require retraining or adaptation for broader applications using transfer learning as in our previous work (Kienbaum et al., 2021; Lozano-Isla et al., 2025). Similar limitations in model generalization across imaging domains are frequently reported in image-based plant phenotyping studies (R.-F. Wang et al., 2025).

## Conclusions

In summary, our findings demonstrate that sowing conditions strongly influenced early development and yield formation in quinoa. While low temperatures likely contributed substantially to differences in emergence and early growth, the observed responses may also reflect variation in season length, accumulated thermal time, and other environmental factors associated with sowing date. Similar interactions between planting date, temperature, and accumulated radiation have been reported previously in quinoa and may jointly shape phenological development and productivity (Hirich et al., 2014). The differences in germination traits, development and yield between coastal and highland quinoa suggest that breeding efforts can focus on combining the advantageous traits of both ecotypes that exhibit a high heritability to develop improved quinoa varieties for production environments in temperate regions in which periods of cold temperature or even frost at early growth stages can be expected (Figure 8). The use of enabling technologies, such as AI-supported large-scale plant phenotyping, can accelerate this process by enhancing trait selection and breeding efficiency. Beyond breeding applications, future integration of transcriptomic and multi-omics analyses with phenotypic data may help uncover the molecular and physiological mechanisms underlying ecotypic differences in germination performance, phenology, and yield formation under cold stress and changing environments (Lai & Zhou, 2025; Maldonado-Taipe et al., 2024).

**Figure 8.**
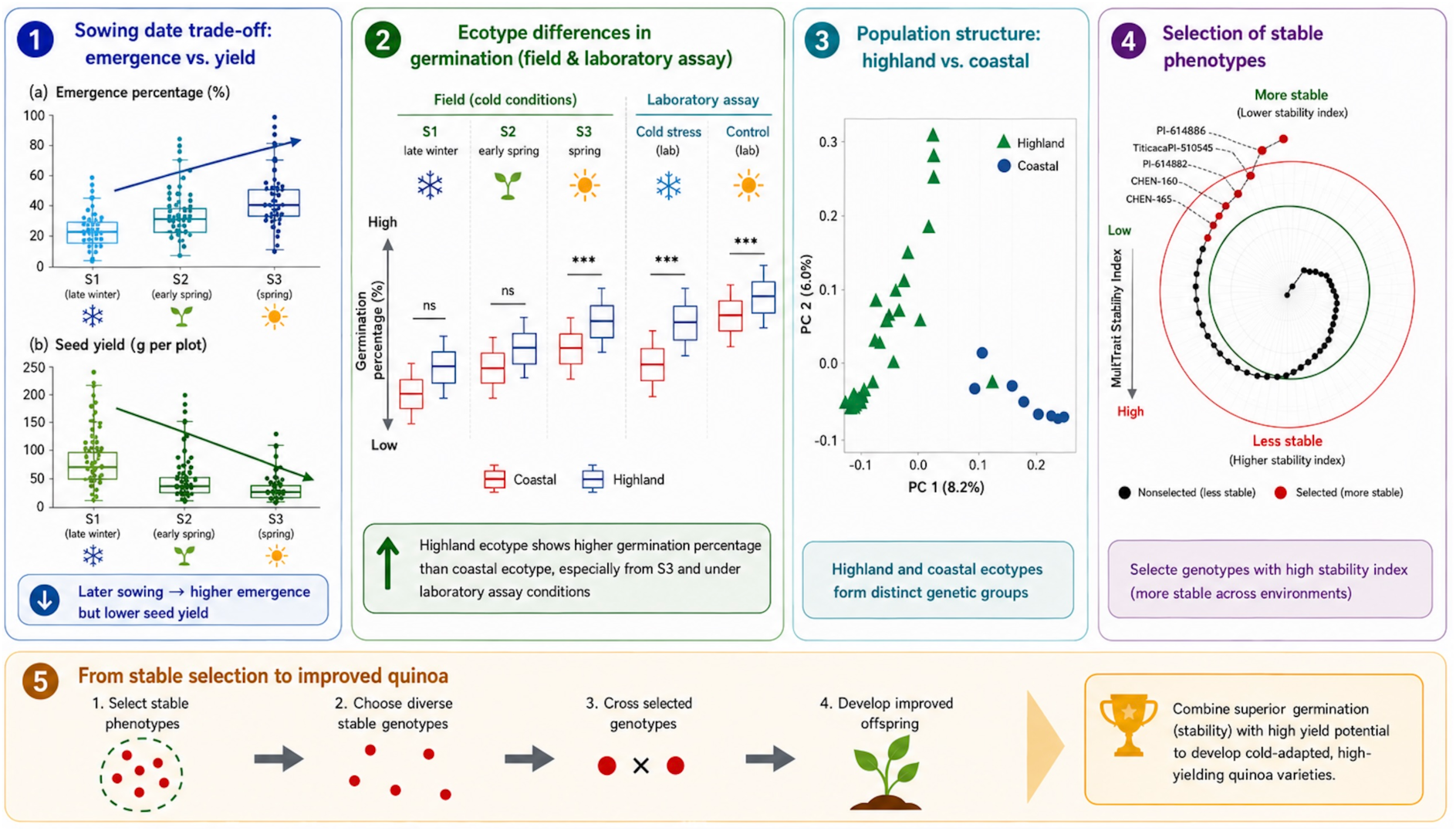
Schematic illustration of how the investigation of germination traits, phenology, and yield can support the identification of parental genotypes for breeding. The integrated evaluation of multiple traits, including germination performance, phenology, and yield stability, enables the identification of stable genotypes within genetic resources. These selected genotypes can subsequently be used as crossing parents to develop improved and broadly adapted cultivars that combine complementary traits from differently adapted genetic resources.

## Supporting information

R Scripts

## Acknowledgements

We thank Viola Abraham, Vanessa Haseneder, Sophie Otterbach, Yaraslova Tkachenko, Emma Sperling, Quentin Burandt, Ali Baturygil, Flavio Lozano-Isla, and all group members for their assistance in the field. We are grateful to Hans-Peter Piepho for advice on statistical analyses and to Felix Bartusch for support with BinAC. Computational resources were provided by the High Performance and Cloud Computing Group at the University of Tübingen, the state of Baden-Württemberg through bwHPC, and the German Research Foundation (DFG, grant no. INST 37/935-1 FUGG).

## Author Contributions

**Niharika Rakasi**

Institute of Plant Breeding, Seed Science and Population Genetics, University of Hohenheim

**Contributions :** Study design, Performing experiments and Data collection, Data and codes curation Statistical analysis, Writing- original draft and, Editing

**Lydia Kienbaum**

Institute of Plant breeding, Seed Science and Population Genetics, University of Hohenheim

**Contributions :** Deep learning image analysis, Data and codes curation, Writing, Review, and Editing

**Katharina B. Böndel**

Institute of Plant breeding, Seed science and Population Genetics, University of Hohenheim

**Contributions :** Study design, Analysis, Writing, Review, and Editing, and Supervision

**Jan-David Wiederstein**

Institute of Plant breeding, Seed science and Population Genetics, University of Hohenheim

**Contribution :** Performing experiments and data collection

**Naveen Kumar Gangaraju**

Institute of Crop Science, University of Hohenheim

**Contribution :** Performing experiments and data collection

**Sandra M. Schmöckel**

Institute of Crop Science, University of Hohenheim

**Contributions :** Study design, Performing experiments and data collection, Review, and Editing

**Karl J. Schmid**

Institute of Plant Breeding, Seed Science and Population Genetics, University of Hohenheim

**Contributions :** Study design, Review, Editing, and Supervision

## Data availability

All phenotypic data and R scripts used for the analysis are available as supplementary material. The raw seed germination images are available from the corresponding author upon request.

