## Supplementary Figures for "Phenotypic differentiation between highland and coastal quinoa under cold stress conditions"

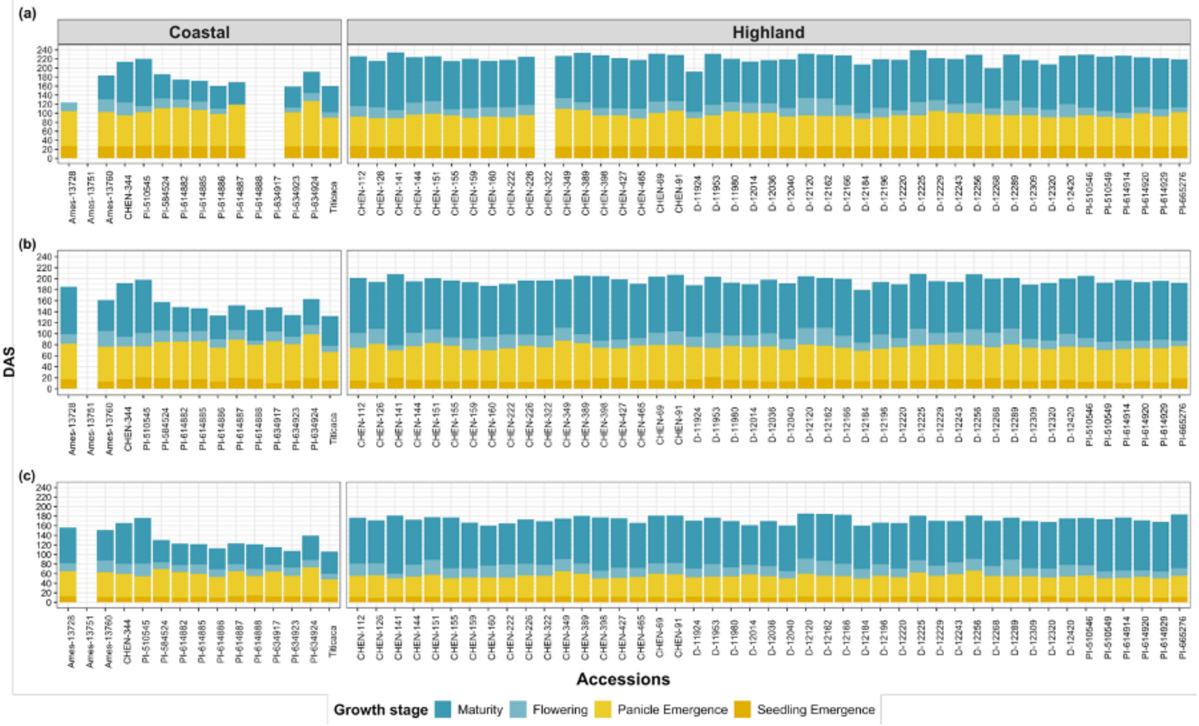


**Figure S1. Estimated means (BLUEs) of different growth stages of the 60 quinoa accessions at three sowing dates** (a) S1, (b) S2, and (c) S3. Within each plot the x-axis represents the accessions, and the y-axis represents days after sowing (DAS). The coastal accessions are represented in a small left-side grid and the highland accessions are represented in a long right-side grid. Each stacked bar represents seedling emergence, panicle emergence, flowering, and maturity in the order of base to top respectively.

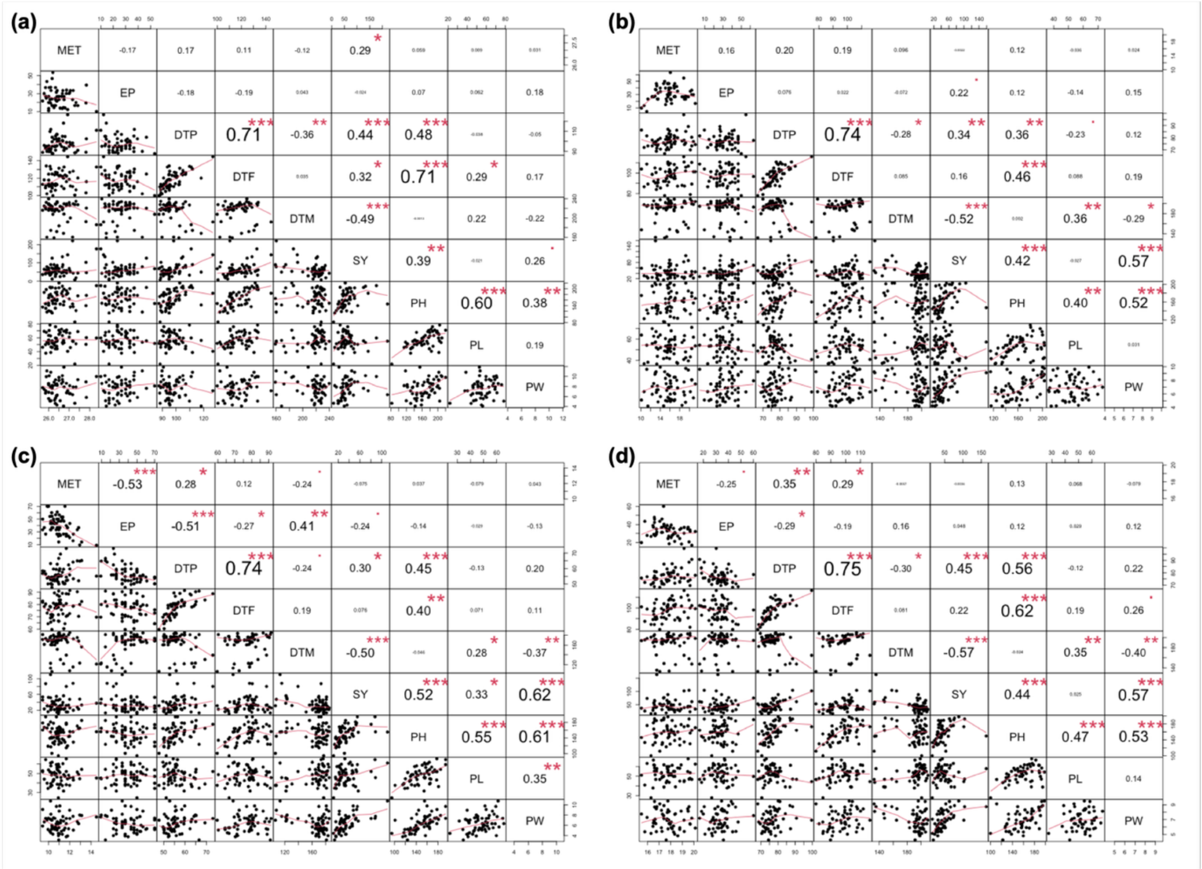


**Figure S2**. **Pearson's correlation coefficients among nine traits** in (a) S1, (b) S2, (c) S3, and (d) all sowing dates combined. The traits are given in the diagonal of each plot: mean emergence time (MET), emergence percentage (EP), days to panicle emergence (DTP), days to flowering (DTF), days to maturity (DTM), seed yield (SY), plant height (PH), panicle length (PL), and panicle width (PW). Above the diagonal are Pearson's correlation coefficients along with their significance levels (***$p<0.001$, **$p<0.01$). The bottom left boxes are scattered plots with fitted lines (in red), representing pairwise relationships between the traits.

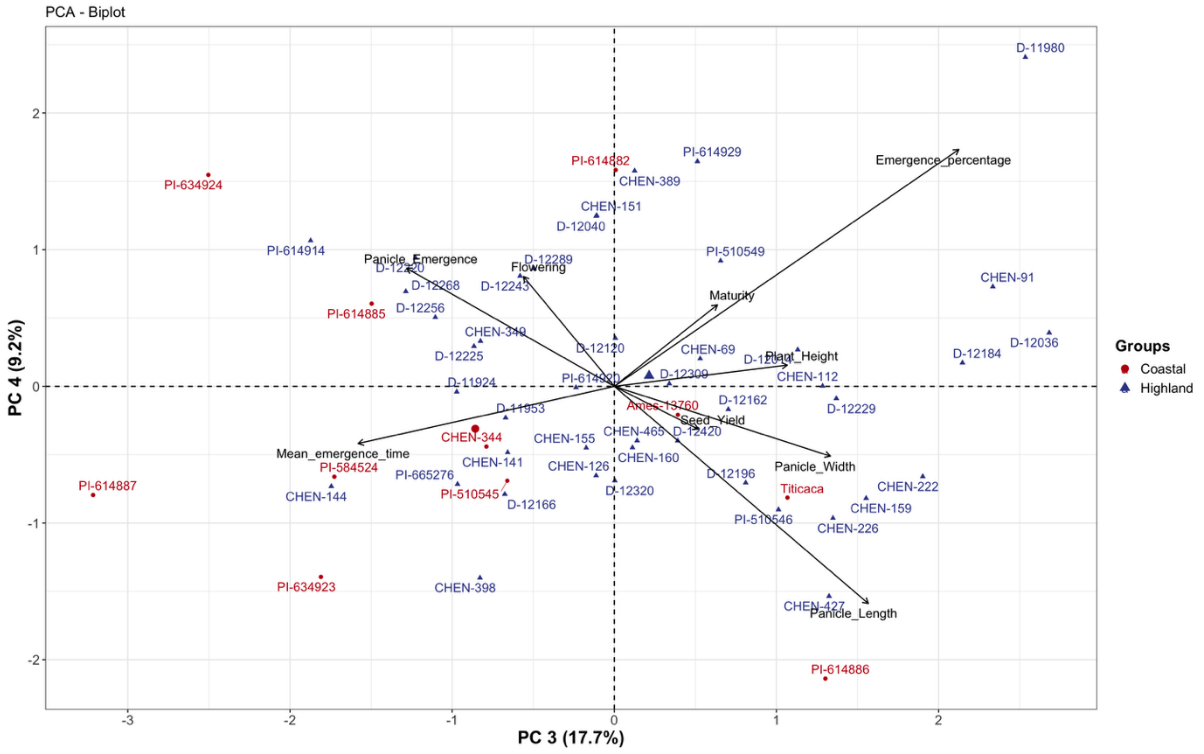


**Figure S3. PCA biplot of nine traits with third and fourth principal components** (PC 3 = 17.7% and PC 4 = 9.2%). The accessions were colored according to their ecotype (coastal in dark red and highland in blue).

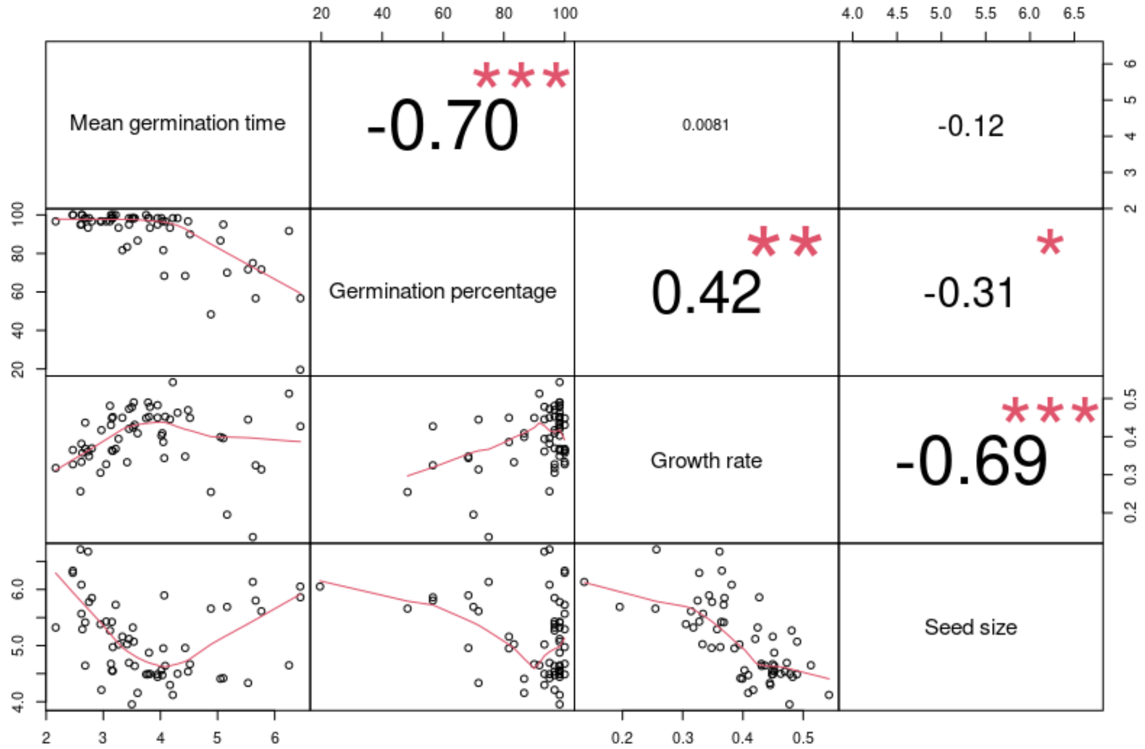


**Figure S4. Pearson's correlation coefficients from laboratory germination experiment**: Traits are given in the diagonal of each plot: mean germination time, germination percentage, growth rate and seed size. Pearson's correlation coefficients along with their significance levels (***p<0.001, **p<0.01). The bottom left boxes are scattered plots with fitted lines (in red), representing pairwise relationships between the traits.

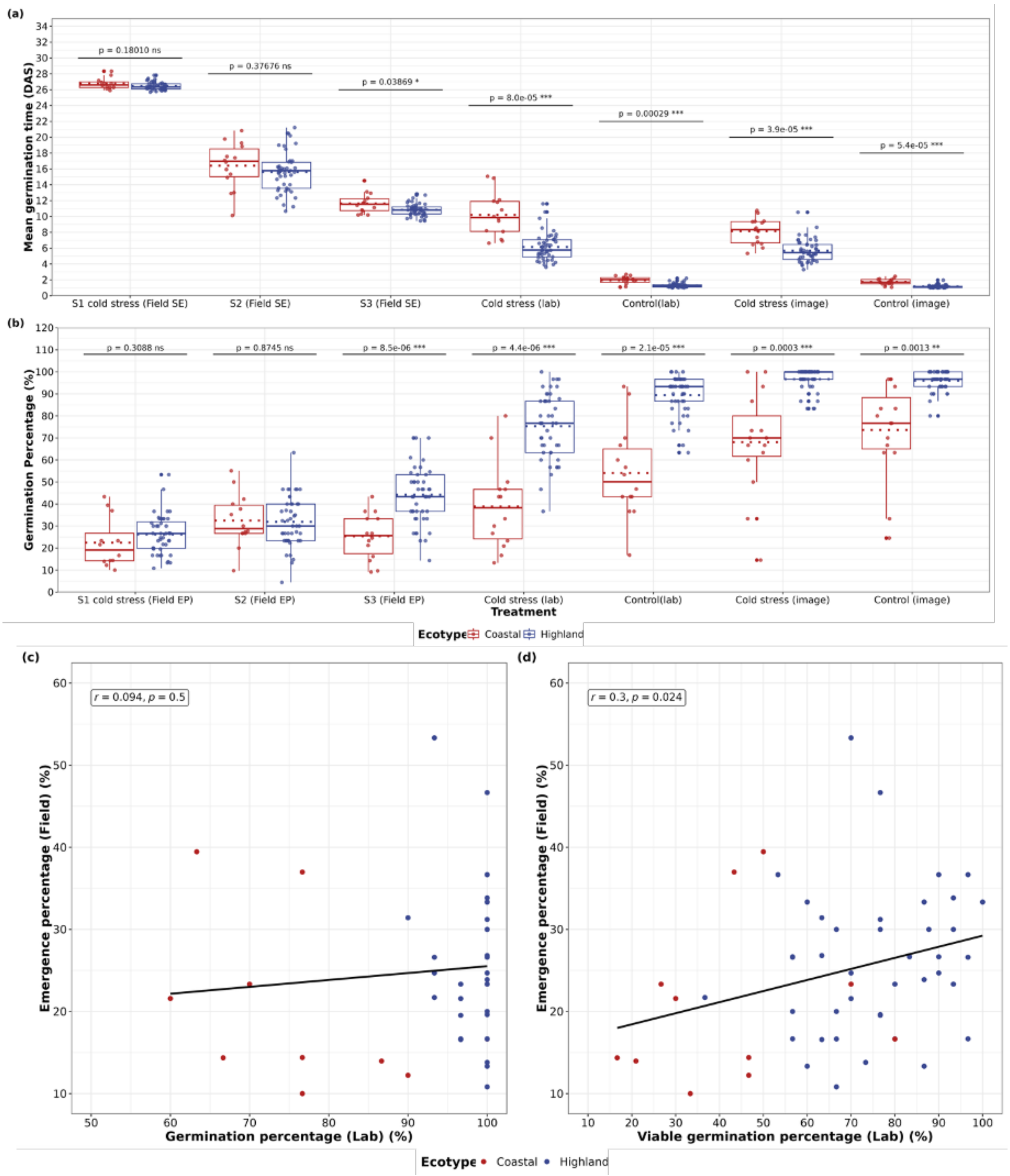


**Figure S5. Boxplots showing the estimated means (BLUEs) of 60 quinoa accessions from the coastal (dark red) and highland ecotypes (blue) for different traits**. The traits measured include (a) mean germination time (DAS) (b) germination percentage. The x-axis represents the treatments: S1, S2, and S3 represent the different sowing dates in the field experiment, cold (lab) and control (lab), cold (image) and control (image), are observations from cold and control temperature treatments under lab conditions, respectively, and, while the y-axis indicates the respective measurement units for each trait. The dotted lines represents the mean. Statistical significance between the ecotypes is indicated by p-values (t-test) and asterisks (*p< 0.05, **p < 0.01, ***p < 0.001, ****p < 0.0001), and "ns" denotes nonsignificant differences. (c) and (d) correlation plots of field seedling emergence %(y-axis) with laboratory germination % (x-axis) and viable germination percentage % (x-axis) respectively and  for the trait BLUEs. The r values represent the correlation coefficient value and the $p < 0.05$ means the correlation is significant. The accessions were colored according to their ecotype (coastal in dark red and highland in blue).

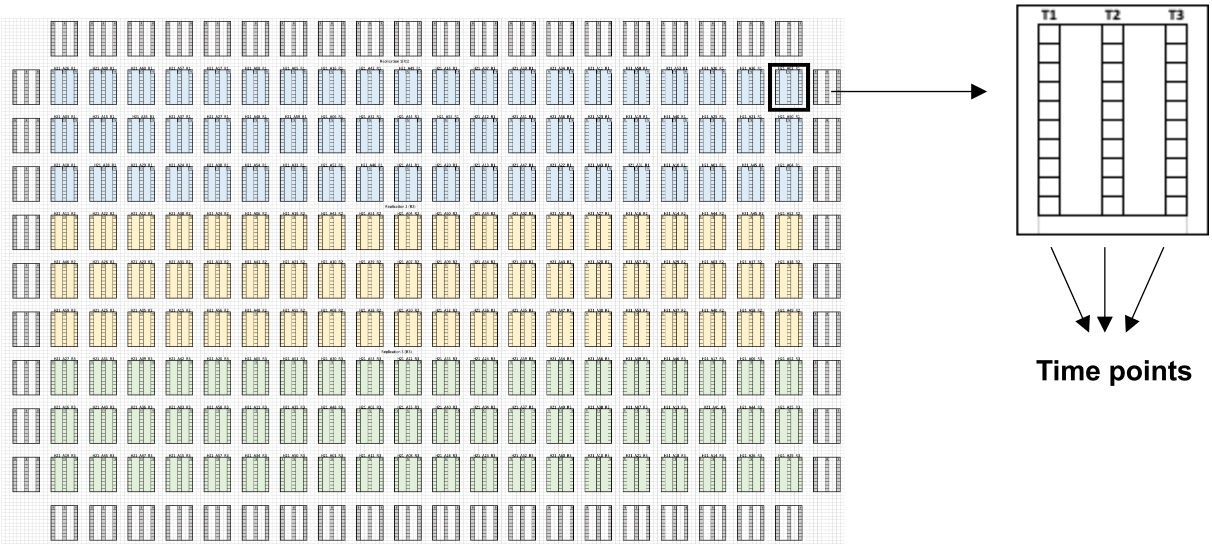


**Figure S6. Experimental field layout with three replications** R1 (blue), R2 (yellow), and R3 (green). The 60 accessions were randomly allocated to main plots within each replication.  The main plot was split into three sub plots S1, S2, and S3 which represent the three different sowing dates. In S1 and S2 one seed was sown per hill, while in S3 three seeds were sown. The entire experimental unit was surrounded by border plants.

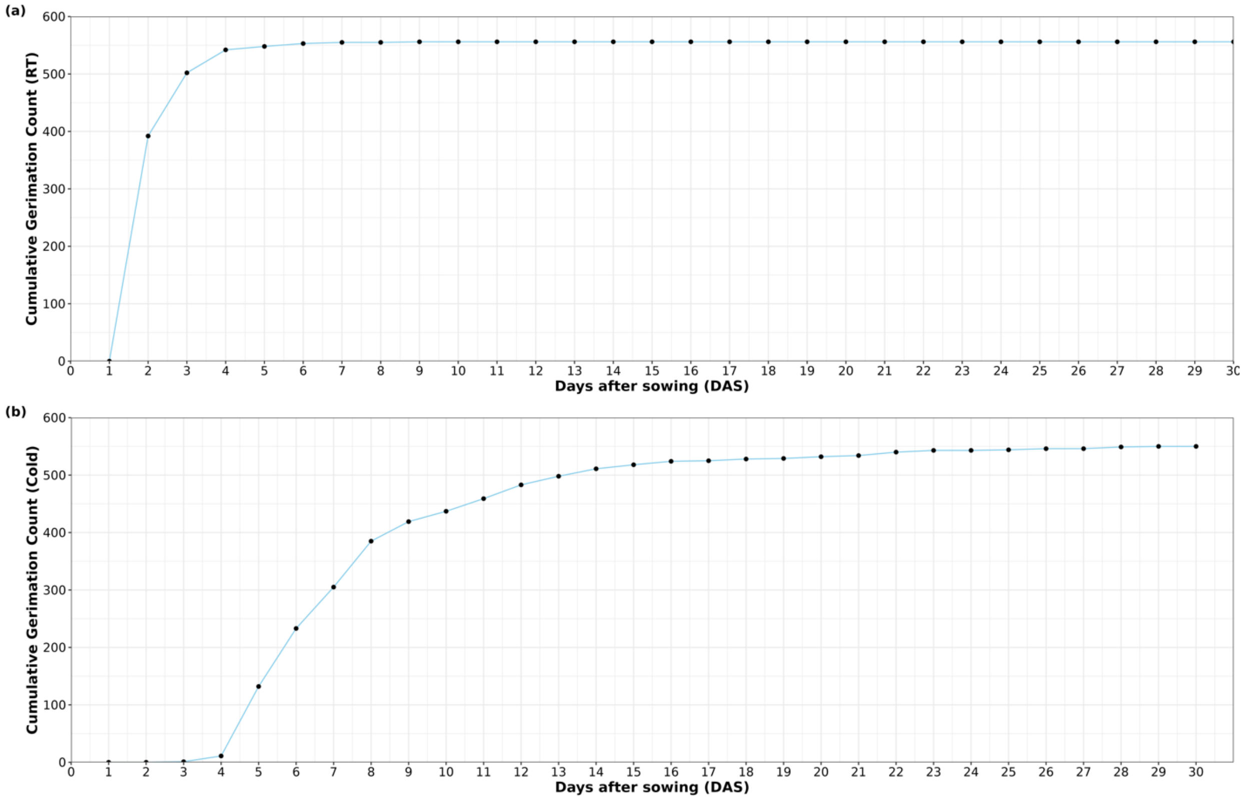


**Figure S7. Cumulative germination counts in laboratory experiment under two treatments**: room temperature (RT) (a) and cold stress (b). The days after sowing (DAS) are shown on the x-axis and the cumulative germination counts are shown on the y-axis.

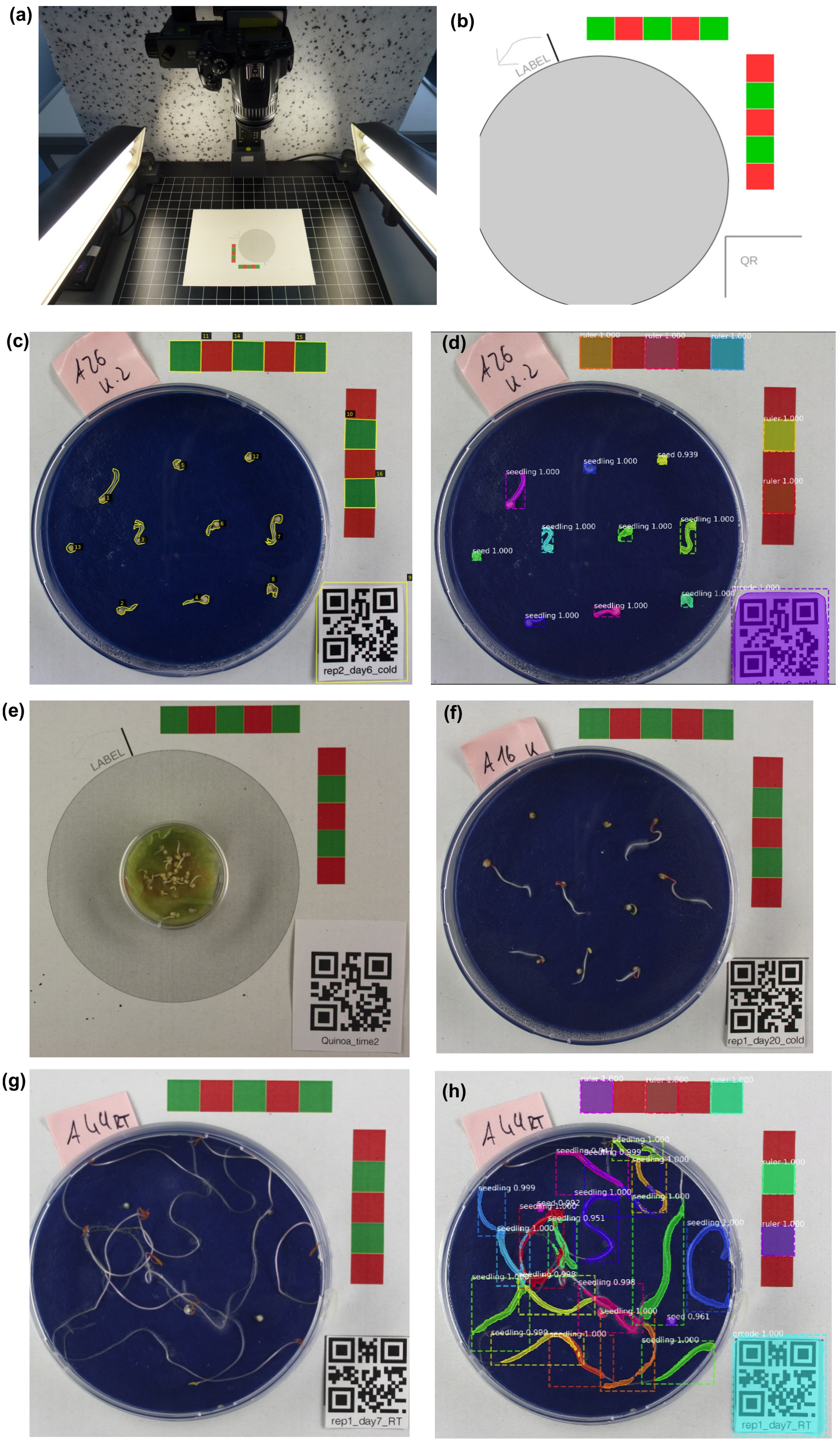


**Figure S8. Photography, annotation and image detection**. (a) Photographic set-up in which the Petri dishes with the seeds and seedlings were photographed. (b) Layout with ruler elements on which the Petri dish and the respective QR-code was positioned for photography. (c) Example image that was annotated with VIA: ruler elements (10, 11, 14-16), QR-code (9), seeds (12, 13), and seedlings (1-8). (d) The same example image as in (c) with detection: the QR-code, the five ruler elements, two seeds and eight seedlings were detected correctly. (e) Example image of the other images which were done with smaller Petri dishes and an uneven and differently colored underground. (f) Example image of an accession where one seed (middle right) that germinated but then did not develop into a viable seedling. (g) Example image with older and very long seedlings that were not detectable anymore. (h) The same example image as in (g) with annotation. A total of 18 seedlings and two seeds was detected while the correct number was ten seedlings and zero seeds.

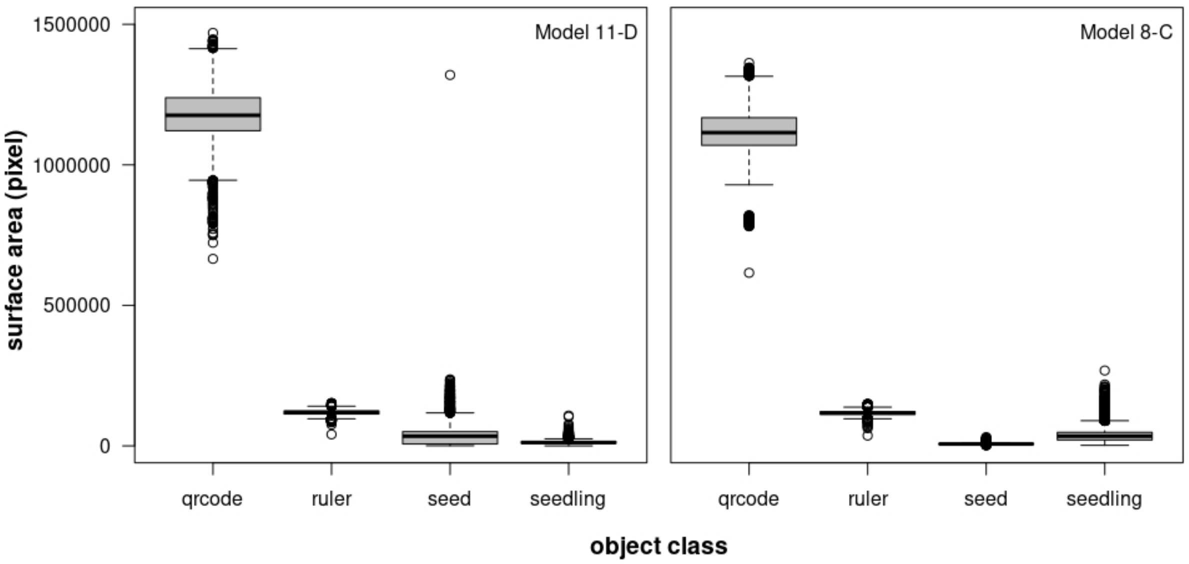


**Figure S9. Detected surface area by the deep learning image analysis pipeline.** Surface area in pixel for the four object classes (QR code, ruler, seed, and seedling) for the two best trained models 11-D (left) and 8-C (right).
