## Supplementary Tables for "Phenotypic differentiation between highland and coastal quinoa under cold stress conditions"

**Table S1.** Weather conditions during the crop growth period

| Month | Relative humidity (%) | | Monthly soil temperature (°C) | | Monthly air temperature (°C) | | Frost days* | |
|---|---|---|---|---|---|---|---|---|
| | Mean | Min. | Max. | Mean | Min. | Max. | Mean | |
| March | 71.2 | 2.7 | 11.9 | 5.6 | -5.4 | 24.5 | 5.0 | 14 |
| April | 65.0 | 4.3 | 12.4 | 8.3 | -4.3 | 23.6 | 6.9 | 11 |
| May | 72.2 | 8.1 | 14.5 | 11.6 | -1.4 | 30.7 | 10.9 | 1 |
| June | 74.0 | 12.3 | 20.8 | 17.6 | 4.1 | 33.8 | 18.9 | 0 |
| July | 79.5 | 16.6 | 22.4 | 19.2 | 8.7 | 30.3 | 18.4 | 0 |
| August | 82.3 | 15.7 | 21.1 | 18.4 | 6.8 | 33.6 | 16.8 | 0 |
| September | 81.0 | 13.1 | 18.6 | 16.3 | 2.2 | 29.6 | 15.3 | 0 |
| October | 86.2 | 7.4 | 15.2 | 11.1 | -2.8 | 22.6 | 8.4 | 4 |

†Frost days represents the number of days when the minimum air temperature is below 0°C.

**Table S2.** Coefficient of vectors on the principal components

| Traits | PC 1 | PC 2 | PC 3 | PC 4 |
|---|---|---|---|---|
| MET | 0.277 | 0.359 | -0.515 | -0.136 |
| EP | -0.123 | -0.087 | 0.693 | 0.565 |
| DTP | 0.816 | 0.065 | -0.418 | 0.282 |
| DTF | 0.742 | 0.451 | -0.183 | 0.262 |
| DTM | -0.39 | 0.794 | 0.207 | 0.194 |
| PH | 0.814 | 0.301 | 0.348 | 0.051 |
| PL | 0.196 | 0.601 | 0.51 | -0.518 |
| PW | 0.651 | -0.324 | 0.434 | -0.166 |
| SY | 0.708 | -0.48 | 0.169 | -0.102 |

†Note: Mean emergence time (MET), emergence percentage (EP), days to panicle emergence (DTP), days to flowering (DTF), days to maturity (DTM), plant height (PH), panicle length (PL), panicle width (PW), seed yield (SY).

**Table S3.** Estimates of the original mean (Xo), the mean of the selected individuals (Xs), and the selection differential (SD).

| Trait | $X_o$ | $X_s$ | SD |
|---|---|---|---|
| MET | 17.8 | 17.2 | -0.521 |
| EP | 33.3 | 34.2 | 0.843 |
| DTF | 97.4 | 94.9 | -2.47 |
| DTM | 189 | 167 | -21.8 |
| PH | 158 | 157 | -1.31 |
| PW | 6.95 | 8.29 | 1.34 |
| SY | 48.5 | 70.7 | 22.2 |

†Note: Mean emergence time (MET), emergence percentage (EP), days to flowering (DTF), days to maturity (DTM), plant height (PH), panicle width (PW), seed yield (SY).

**Table S4.** Estimated means (BLUEs) of the top eight genotypes selected by multi-trait stability index for their stability in three sowing dates (S1, S2, and S3) for the traits Mean emergence time (MET), emergence percentage (EP), days to flowering (DTF), days to maturity (DTM), plant height (PH), panicle width (PW) and seed yield (SY)

| Time point | Accession | MET | EP | DTF | DTM | PH | PW | SY |
|---|---|---|---|---|---|---|---|---|
| S1 | Titicaca | 25.9 | 43.3 | 102.1 | 160.2 | 117.56 | 11.12 | 78.92 |
| S1 | PI-614886 | 27.1 | 23.3 | 110.4 | 160.4 | 146.0 | 6.35 | 220.2 |
| S1 | PI-614882 | 26.10 | 36.9 | 130.3 | 174.43 | 202.7 | 10.8 | 90.31 |
| S1 | PI-634923 | 26.2 | 14.3 | 112.3 | 159.0 | 165.3 | 8.8 | 89.8 |
| S1 | CHEN-160 | 26 | 26.6 | 113.3 | 216 | 121.7 | 8.3 | 58.6 |
| S1 | CHEN-465 | 26.1 | 23.8 | 109.9 | 217.5 | 172.2 | 6.6 | 71.9 |
| S1 | CHEN-69 | 26.4 | 33.3 | 126 | 232 | 161.8 | 8.25 | 73.7 |
| S1 | CHEN-91 | 26.3 | 36.6 | 126.6 | 229 | 201.8 | 9.3 | 74.5 |
| S2 | Titicaca | 14.9 | 50 | 78 | 132 | 115.1 | 9.25 | 49.6 |
| S2 | PI-614886 | 13.01 | 37.76 | 90.2 | 132.8 | 147.3 | 9.11 | 158.9 |
| S2 | PI-614882 | 15.9 | 40 | 103.3 | 148.6 | 201 | 8.63 | 88.5 |
| S2 | PI-634923 | 15.3 | 20 | 94.6 | 134 | 136.8 | 6.5 | 42.3 |
| S2 | CHEN-160 | 15.2 | 23.3 | 94.6 | 187 | 144.8 | 7.5 | 45.45 |
| S2 | CHEN-465 | 15.09 | 14.7 | 92.0 | 190.8 | 157.1 | 5.01 | 44.6 |
| S2 | CHEN-69 | 15.7 | 32.5 | 101.4 | 203.8 | 166.8 | 6.8 | 46.7 |
| S2 | CHEN-91 | 14 | 40 | 104 | 207 | 199.9 | 9.9 | 61.4 |
| S3 | Titicaca | 10.3 | 36.6 | 59.6 | 106 | 101.2 | 7 | 31.3 |
| S3 | PI-614886 | 10.19 | 33.3 | 69 | 113 | 155.3 | 10.76 | 110.49 |
| S3 | PI-614882 | 10.1 | 43.3 | 78 | 122.6 | 181.05 | 7.16 | 51.6 |
| S3 | PI-634923 | 11.9 | 21.2 | 72.6 | 107.4 | 144.5 | 8.94 | 47.6 |

| | | | | | | | | |
|---|---|---|---|---|---|---|---|---|
| S3 | CHEN-160 | 10.6 | 43.3 | 77 | 160.3 | 131.7 | 7.3 | 34.4 |
| S3 | CHEN-465 | 10.3 | 54.5 | 71.8 | 166.0 | 139.7 | 6 | 34.2 |
| S3 | CHEN-69 | 11.05 | 36.6 | 81.6 | 181 | 160 | 9.7 | 47.4 |
| S3 | CHEN-91 | 9.45 | 50 | 82.6 | 181.3 | 178.3 | 9.08 | 53.8 |

**Table S5.** Wald F-test results for fixed effects in quinoa cold stress laboratory experiment, reported as p-values

| Method | Source of variation | | | | |
|---|---|---|---|---|---|
| | Traits | Rep | Treatment | ID | ID:Treatment |
| Manual documentation | MGT | 0.202 | 0 **** | 0 **** | 0 **** |
| | GP | 0.737 | 0.567 | 0 **** | 0 **** |
| | VGP | 0.306 | 0.002 ** | 0 **** | 0.001 *** |
| Image analysis | MGT | 0.007** | 0 **** | 0 **** | 0 **** |
| | GP | 0.434 | 0.423 | 0 **** | 0.003** |
| | Growth Rate | 0 **** | 0 **** | 0 **** | 0 **** |

†Note: Mean germination time (MGT), germination percentage (GP), viable germination percentage (VGP). Asterisks (*p < 0.05, **p < 0.01, ***p < 0.001, ****p < 0.0001).

**Table S6.** Details of sowing and harvesting of the three different time points.

| Time point | Date of sowing | Seeds per hill | Plants per plot | Date of final harvesting |
|---|---|---|---|---|
| S1 | 04 March | 1 | 10 | 31$^{st}$ October |
| S2 | 01 April | 1 | 10 | 31$^{st}$ October |
| S3 | 30 April | 3 | 30 | 31$^{st}$ October |

**Table S7.** mAP@[.5:.95] and penalty result of models with different hyperparamers. Each unique hyperparameter combination (models 1-12) was replicated five times (A-E).

| Model | Loss | Heads.m | Mask_resolution | mAP@[.5:.95] | Penalty |
|---|---|---|---|---|---|
| 1-A | mask1class1 | heads | 28x28 | 58.47 | - |
| 1-B | mask1class1 | heads | 28x28 | 54.60 | - |
| 1-C | mask1class1 | heads | 28x28 | 54.99 | - |
| 1-D | mask1class1 | heads | 28x28 | 51.62 | - |
| 1-E | mask1class1 | heads | 28x28 | 52.33 | - |
| 2-A | mask1class10 | heads | 28x28 | 60.26 | - |
| 2-B | mask1class10 | heads | 28x28 | 61.69 | - |
| 2-C | mask1class10 | heads | 28x28 | 60.26 | - |
| 2-D | mask1class10 | heads | 28x28 | 55.07 | - |
| 2-E | mask1class10 | heads | 28x28 | 55.63 | - |
| 3-A | mask10class1 | heads | 28x28 | 57.38 | - |
| 3-B | mask10class1 | heads | 28x28 | 55.53 | - |
| 3-C | mask10class1 | heads | 28x28 | 51.55 | - |
| 3-D | mask10class1 | heads | 28x28 | 57.22 | - |
| 3-E | mask10class1 | heads | 28x28 | 57.05 | - |
| 4-A | mask1class1 | heads | 56x56 | 53.89 | - |
| 4-B | mask1class1 | heads | 56x56 | 59.08 | - |
| 4-C | mask1class1 | heads | 56x56 | 59.48 | - |
| 4-D | mask1class1 | heads | 56x56 | 58.10 | - |

| | | | | | |
|---|---|---|---|---|---|
| 4-E | mask1class1 | heads | 56x56 | 57.12 | 57 |
| 5-A | mask1class10 | heads | 56x56 | 57.63 | - |
| 5-B | mask1class10 | heads | 56x56 | 62.01 | - |
| 5-C | mask1class10 | heads | 56x56 | 60.80 | - |
| 5-D | mask1class10 | heads | 56x56 | 55.19 | - |
| 5-E | mask1class10 | heads | 56x56 | 56.76 | - |
| 6-A | mask10class1 | heads | 56x56 | 58.37 | - |
| 6-B | mask10class1 | heads | 56x56 | 59.36 | - |
| 6-C | mask10class1 | heads | 56x56 | 54.03 | - |
| 6-D | mask10class1 | heads | 56x56 | 58.73 | - |
| 6-E | mask10class1 | heads | 56x56 | 54.64 | - |
| 7-A | mask1class1 | all | 28x28 | 55.55 | - |
| 7-B | mask1class1 | all | 28x28 | 53.02 | - |
| 7-C | mask1class1 | all | 28x28 | 56.75 | - |
| 7-D | mask1class1 | all | 28x28 | 58.34 | - |
| 7-E | mask1class1 | all | 28x28 | 55.21 | - |
| 8-A | mask1class10 | all | 28x28 | 54.49 | - |
| 8-B | mask1class10 | all | 28x28 | 63.35 | 23 |
| 8-C | mask1class10 | all | 28x28 | 63.16 | 22 |
| 8-D | mask1class10 | all | 28x28 | 59.58 | - |
| 8-E | mask1class10 | all | 28x28 | 57.20 | - |
| 9-A | mask10class1 | all | 28x28 | 59.66 | - |
| 9-B | mask10class1 | all | 28x28 | 52.96 | - |
| 9-C | mask10class1 | all | 28x28 | 60.31 | - |

| | | | | | |
|---|---|---|---|---|---|
| 9-D | mask10class1 | all | 28x28 | 62.95 | - |
| 9-E | mask10class1 | all | 28x28 | 54.81 | - |
| 10-A | mask1class1 | all | 56x56 | 58.73 | - |
| 10-B | mask1class1 | all | 56x56 | 59.39 | - |
| 10-C | mask1class1 | all | 56x56 | 61.25 | - |
| 10-D | mask1class1 | all | 56x56 | 62.88 | - |
| 10-E | mask1class1 | all | 56x56 | 58.62 | - |
| 11-A | mask1class10 | all | 56x56 | 61.86 | - |
| 11-B | mask1class10 | all | 56x56 | 58.38 | - |
| 11-C | mask1class10 | all | 56x56 | 62.84 | - |
| 11-D | mask1class10 | all | 56x56 | 64.39 | 24 |
| 11-E | mask1class10 | all | 56x56 | 64.38 | 24 |
| 12-A | mask10class1 | all | 56x56 | 53.94 | - |
| 12-B | mask10class1 | all | 56x56 | 58.37 | - |
| 12-C | mask10class1 | all | 56x56 | 55.47 | - |
| 12-D | mask10class1 | all | 56x56 | 49.78 | - |
| 12-E | mask10class1 | all | 56x56 | 59.47 | - |

**Table S8.** Least squares means of mAP values of significant model parameters. Means with the same letter are not significantly different.

| Parameter | p-value | Least squares means | Standard error |
|---|---|---|---|
| **heads.m** | 0.048222 * | | |
| all | | 58.57[a] | 0.56 |
| heads | | 56.96[b] | 0.56 |
| **loss** | 0.003501 ** | | |
| mask1class10 | | 59.75[a] | 0.69 |
| mask1class1 | | 56.97[b] | 0.69 |
| mask10class1 | | 56.58[b] | 0.69 |

**Table S9.** Result of penalty scoring of the validation images from another germination experiment used for evaluation of the pipe-line.

| Image Set | Model Update | Seed/ seedlings error | Missed Seed/ seedlings | Multiple detections of same seed/ seedling | Multiple seed/ seedlings detected as one | Other update | Total Penalty count |
|---|---|---|---|---|---|---|---|
| All images (207 seeds) | without update | 0 | 203 | 0 | 2 | 13 | 218 |
| | 10 epochs, replicate 1 | 12 | 117 | 5 | 7 | 3 | 144 |
| | 10 epochs, replicate 2 | 20 | 106 | 1 | 5 | 0 | 132 |
| | 10 epochs, replicate 3 | 8 | 138 | 1 | 10 | 0 | 157 |
| | 30 epochs, replicate 1 | 14 | 96 | 9 | 21 | 5 | 145 |
| | 30 epochs, replicate 2 | 11 | 79 | 4 | 13 | 3 | 110 |
| | 30 epochs, replicate 3 | 12 | 83 | 3 | 11 | 3 | 112 |
| | 50 epochs, replicate 1 | 9 | 86 | 3 | 9 | 3 | 110 |
| | 50 epochs, replicate 2 | 51 | 109 | 0 | 1 | 2 | 163 |
| | 50 epochs, replicate 3 | 20 | 127 | 10 | 21 | 9 | 187 |

| | | | | | | | |
|---|---|---|---|---|---|---|---|
| Images with 10-30 seeds (157 seeds) | without update | 0 | 156 | 0 | 1 | 6 | 163 |
| | 10 epochs, replicate 1 | 9 | 108 | 0 | 6 | 3 | 126 |
| | 10 epochs, replicate 2 | 14 | 95 | 0 | 4 | 0 | 113 |
| | 10 epochs, replicate 3 | 5 | 127 | 0 | 8 | 0 | 140 |
| | 30 epochs, replicate 1 | 10 | 91 | 2 | 20 | 1 | 124 |
| | 30 epochs, replicate 2 | 9 | 77 | 0 | 12 | 2 | 100 |
| | 30 epochs, replicate 3 | 9 | 82 | 1 | 10 | 1 | 103 |
| | 50 epochs, replicate 1 | 8 | 84 | 0 | 8 | 1 | 101 |
| | 50 epochs, replicate 2 | 26 | 101 | 0 | 1 | 0 | 128 |
| | 50 epochs, replicate 3 | 12 | 116 | 4 | 18 | 7 | 157 |
| Images with 5 seeds (50 seeds) | without update | 0 | 47 | 0 | 1 | 7 | 55 |
| | 10 epochs, replicate 1 | 3 | 9 | 5 | 1 | 0 | 18 |
| | 10 epochs, replicate 2 | 6 | 11 | 1 | 1 | 0 | 19 |
| | 10 epochs, replicate 3 | 3 | 11 | 1 | 2 | 0 | 17 |
| | 30 epochs, replicate 1 | 4 | 5 | 7 | 1 | 4 | 21 |

| | | | | | | | |
|---|---|---|---|---|---|---|---|
| | 30 epochs, replicate 2 | 2 | 2 | 4 | 1 | 1 | 10 |
| | 30 epochs, replicate 3 | 3 | 1 | 2 | 1 | 2 | 9 |
| | 50 epochs, replicate 1 | 1 | 2 | 3 | 1 | 2 | 9 |
| | 50 epochs, replicate 2 | 25 | 8 | 0 | 0 | 2 | 35 |

†1: Other errors included for example when part of the labelling or the Petri dish was detected as seedling or when a seed was detected as QR-code.

**Table S10.** Result of penalty scoring of the random images from another germination experiment analyzed with the best updated model (30 epochs, replicate 2).

| Image set | All 20 images (191 seeds) | Images with 10+ seeds (141 seeds) | Images with 5 seeds (50 seeds) |
|---|---|---|---|
| **Seed/seedling error** | 16 | 11 | 5 |
| **Missed seeds/seedlings** | 77 | 71 | 6 |
| **Multiple detections of the same seed/seedling** | 3 | 1 | 2 |
| **Multiple seeds/seedlings detected as one** | 11 | 11 | 0 |
| **Other errors**[1] | 3 | 1 | 2 |
| **Total penalty count** | 110 | 95 | 15 |

†1: Other errors included for example when part of the labelling or the Petri dish was detected as seedling or when a seed was detected as QR-code.
