## Supplementary Text for "Phenotypic differentiation between highland and coastal quinoa under cold stress conditions"

This supplementary text contains additional details of extended methods, results and discussion of the deep learning image analysis pipeline that were not included in the main text. We specially focus on the experimental setup to capture the images, on the methodology of rating the detection results, and on the choice, analysis and optimization of Mask R-CNN hyper-parameters. This information is intented to support and clarify the findings discussed.

#### Results

##### Performance of deep learning approach for image analysis of germination in petri dishes

**Statistics of image analysis** We developed a deep learning image analysis pipeline to extract germination progress from images taken during the seed germination experiment, in an automated and objective way. To investigate the influence of the model training parameters, we run the ANOVA model on the m$AP$@[.5:.95] score, a standard metric to evaluate the model's ability to accurately detect and classify objects, in which we dropped the non-significant 3-way interaction first and subsequently also all 2-way interactions because they were both not significant. The remaining factors with a significant influence on the m$AP$@[.5:.95] score were the `loss` and `heads.m`.

The `loss` parameter influences the trade-off between optimization on segmentation or classification, the `heads.m` parameter refers to the size of the network architecture used for training. Least squares means for `heads.m` show that training the full layers of the network architecture (`all`) was significantly better than optimizing only on a part of the network (`heads`), indicating that the more intensive training with all layers is required to obtain highly accurate training results (Table S8). Further, the `loss` parameter `mask1class10` was significantly better than all other losses, showing that the classes were rather difficult to distinguish and a higher weight on classes rather than on masks was necessary to improve the results.

**Overall accuracy and results of image analysis pipeline** The goal of our pipeline was to correctly detect seeds, seedlings, ruler elements and QR-codes on each image as well as to obtain surface areas of seeds and seedlings to investigate the rate at which the seedlings grew over the course of the experiment. Since after training, different models were best for the m$AP$@[.5:.95] score and our penalty approach, where we scored misclassifications and non-detected elements, we decided to execute the pipeline with both models. Overall, model 11-D (best according to its m$AP$@[.5:.95] score) was able to analyze 5867 of the 6937 images (84.58%) and model 8-C (best according to its penalty) 6934 (99.96%). To evaluate which of the two models performed better in detecting the different elements on the images we first estimated the number of possible misclassifications based on the extracted surface area. Since the seedling surface area showed the greatest distribution and overlap with seeds (which was to be expected since seedlings are initially only slightly larger than seeds) and there was no apparent misclassification of ruler elements or QR-codes (except one QR-code which was classified as a seed in model 11-D, [Figure S9]), we focused on the seeds, i.e. we estimated how many non-seed objects might be classified as seeds. To do so we used the seed calls from the images taken at day 1 of replicate 1 from the Petri dishes kept under cold conditions as a reference set and calculated the maximum seed area. We selected this subset as reference because it showed no misclassifications. Model 11-D had a total of 36,465 seed calls of which 22,990 had a surface area that was more than 10% larger than the largest seed in our reference set. In comparison, model 8-C had a total of 16,124 seed calls of which only 78 had a surface area that was more than 10% larger. Even though the 10% are somewhat arbitrarily set, these numbers suggest that we have to expect many more misclassifications under model 11-D than under model 8-C. As a second check we counted the number of objects of each class per image. This showed that with model 11-D the QR-code was detected only in 5632 of the 5867 images (95.99%) that could be analysed while with model 8-C the QR-code was detected on all images. Furthermore, on a higher number of images less than the expected ten seeds/seedlings were detected with model 11-D than with model 8-C (4237 vs 818). Taken together these findings suggested that model 8-C was performing better (more images analyzed, less misclassifications and less missed objects) and we decided to use the output of the pipeline when run with model 8-C for all analyses. The original output from the pipeline is provided as Data S8 for model 8-C and Data S9 model 11-D.

**Evaluation with other images** In order to investigate whether our newly established pipeline to score germination of quinoa seeds would also work in other set-ups (e.g., different back ground, variable seed numbers) we tested it on a set of images taken during another germination experiment. We observe the highest penalties for the validation images when no update is made where only 1.93% of the seeds or seedlings were detected (Table S9). We tested three different numbers of epochs in the update and penalties for the different epochs varied with the 30 epoch models having on average the lowest penalty. This suggests not only that an update is required but also that 30 epochs are sufficient for the update in our test with 30 training and 20 validation images. Seed and seedling detection also worked better for images with only five seeds. For instance the 30 epoch models had on average 77.3% of the seeds and seedlings detected on images with only five seeds while on the images with ten to 30 seeds and seedlings on average only 31.4% were detected. We then also applied the best updated model to a random subset of 20 images where we could detect 33.3% of seeds and seedlings on images with ten to 30 seeds and seedlings and 74.0% on images with only five seeds or seedlings (Table S10). There have been no problems to detect ruler elements and QR-codes without the update which indicates that the update is primarily required when small objects like seeds are to be detected.

### Discussion

Deep learning approaches have found widespread application in various aspects of crop sciences, including precision agriculture, crop yield prediction, automatic harvesting, weed and plant disease detection, crop research and plant breeding (Jabed & Murad, 2024; Murphy et al., 2024; Upadhyay et al., 2025; Vithlani & Dabhi, 2023). Here, we established a deep learning approach to score germination on images taken daily during a germination experiment conducted under controlled laboratory conditions on Petri dishes. Based on our penalty scoring, we observed a 95.1% detection accuracy for seeds and seedlings and 100% detection accuracy for QR-codes and ruler elements which is slightly higher than 90% accuracy for seeds/seedlings reached in a recent study on vegetables (Nehoshtan et al., 2021). Considering the fact that small objects such as seeds are difficult to detect and to segment from background noise in images, this is a very good result. Traditionally the best deep learning model is chosen based on the m$AP$@[.5:.95] score and other studies have reported 97.9%, 94.2%, 94.3% and 95.39% for *Zea mays*, *Secale cereale*, *Pennisetum glaucum and Oryza sativa*, respectively (Genze et al., 2020; Zhao et al., 2023).However, in our case the model with the best m$AP$@[.5:.95] (64.39, model 11-D) did not have the lowest penalty in a random subset of images and performed worse in comparison to the model with the lowest penalty (model 8-C) which had a slightly lower m$AP$@[.5:.95] score (63.16). This may be because our objects of interest were rather small and therefore difficult to segment properly. Resolution of small objects is limited which makes it difficult for the algorithm to segment them properly from the background and from each other (Liu et al., 2021), which may also explain why the updated pipeline worked better for images of plates with only five seeds than of plates with a higher number where seeds were often very close to each other. It also shows that in some situations the m$AP$@[.5:.95] score may not be the best option to choose the best model and that a visual inspection and evaluation of a random subset of images (as we did here with the penalty approach) is useful to ensure that the best model is chosen. Furthermore, the high correlation between manually documented germination and the germination scoring by the pipeline confirms that our pipeline can detect seeds and seedlings successfully (Figures 6g and 6h).

The evaluation of our pipeline with other images revealed that an update to adjust for a different set-up is required. But if this is done, the pipeline is applicable to other images from seed germination experiments. Using a deep learning approach for scoring germination and other traits has several advantages over manual documentation. It takes less time, for example the manual documentation of the 120 Petri dishes took us each day between 60 and 90 minutes while the same number of Petri dishes could be photographed in less than 10 minutes. It is also less biased because it is objective and therefore highly reproducible. The main advantage however may be that it is much easier to measure the size of the seedlings because it is very difficult to do that manually due to the curly nature of the seedlings. But there are also disadvantages which could be addressed in future improvements of our pipeline. It is currently not possible to differentiate between viable and non-viable seedlings. We observed that several seeds germinated but then stopped in their development and would not have survived if we had planted them (Figure S8f). Since we had documented this manually, we could perform correlations between our field and laboratory experiment with both germinated seeds and viable seedlings and observed a better correlation with viable seedlings than with germinated seeds suggesting that germination alone is not a good indicator of survival or performance under a specific condition (Figures S5c and S5d). An improvement of the pipeline could be achieved, for example, by adding a second step where seedlings are further segmented into leaves, shoot and root and presence of all three would indicate a viable seedling. Another disadvantage was that older seedlings (app. > 2 cm) were often overlapping on the Petri dishes and could therefore not be detected properly anymore (Figure S8g and S8h). More distance between the seeds (and larger Petri dishes and/or fewer seeds per Petri dish) could help to circumvent this problem in future studies. Furthermore, for future studies on germination experiments, we recommend training all layers of the Mask R-CNN architecture and putting a higher weight on the optimization of the classification error in the loss function. Depending on the availability of computing resources, even more than five replications of each parameter combination should be carried out. Also, we noticed that an even underground, an underground color that is of good contrast to the seed color(s) and avoiding handwritten labels (which can occasionally be detected as seedlings) greatly improved the detection. To our knowledge, we are first reporting the automatic measurement of seedling surface in cm² in germination experiments, which is very useful in analyzing germination dynamics in crop research.

#### MATERIALS AND METHODS

##### Development of a deep learning image analysis pipeline to score germination

To extract the germination progress of the seeds and calculate the growth rate of the seedlings per Petri dish in an automated and objective way, we took images in a standardized manner and developed a pipeline which outputs the number and surface area of germinated (seedling) and non-germinated (seed) seeds for each image. In the pipeline, we used a deep learning approach which implements the Mask R-CNN model (Kaiming et al., 2017). In detail, we used the previous adaptation of the matter port implementation (https://github.com/matterport/Mask_RCNN, last accessed on 24.02.2023) for plant phenotyping of maize cobs (Kienbaum et al., 2021) and its further improvement on quinoa panicles (Lozano-Isla et al., 2025) and adjusted it to match the requirements for the detection of seeds and seedling stages and therefore germination dynamics. We first trained a set of possible detection models based on annotated training images and then applied the two most suitable models to the total set of 6937 images to count the number of seeds and seedlings per image and calculate the surface area of each seed/seedling based on ruler elements of known size.

**Experimental set up for taking images of the Petri dishes with seeds and seedlings** First, we developed a standardized setting with external light sources and a ruler layout to photograph the Petri dishes every 24 hours with a Canon EOS 1000D camera to ensure high picture quality and conformity (Figure S8a and S8b). The Petri dishes were positioned in the center, surrounded by a quick response (QR) code and ruler elements (0.95 cm x 0.95 cm), which allowed us to measure the surface area of seeds and seedlings precisely.

**Annotation of training and validation images** We selected 120 images to train and another 30 to validate the models, which results in a split ratio of 4:1 or 80:20. This is a standard split ratio in image analysis with deep learning, because it avoids both under-fitting (i.e., a too simple model with poor performance on training and validation) and over-fitting (i.e., a too complex model with good performance on training data, but poor performance on validation data). Both training and validation images contained a good representation of seed colors, seedling sizes and positions of the seeds/seedlings on the plates and to each other. These criteria were chosen based on preliminary trials. The images were manually annotated with the VIA image annotator software version 2.0.8 (https://www.robots.ox.ac.uk/~vgg/software/via/, last accessed on 18.04.2025, (Dutta & Zisserman, 2019). We annotated four different object classes on which the models were then trained to correctly differentiate between them: seed (not germinated), seedling (germinated, i.e., root clearly visible), ruler and qrcode (Figure S8c).

**Choice of training parameters for MASK R-CNN** Based on our experience from previous studies (Kienbaum et al., 2021; Lozano-Isla et al., 2025) we varied only a few hyperparameters to find a good model. The `loss` (referring to the difference between predicted and ground truth masks) was tested with an equal loss for each the mask and the class optimization (`mask1class1`) and with 10-fold more weight on either the mask (`mask10class1`) or the class (`mask1class10`) because since our data set contained very small objects (i.e., the seeds) it was not known whether it would be more difficult to optimize the segmentation or the classification. Further, the parameter `heads.m` was tested with training only the `heads` of the network architecture and keeping all other layers from the pre-trained coco weights frozen, or training `all` layers. We again chose both options since it was not known whether the more intensive training was needed for the germination images. The `mask_resolution` was varied from the standard `28x28` to a higher resolution of `56x56` (with the additional cost of computational time) at the final mask branch of Mask R-CNN. Combining the different parameter settings resulted in a total of twelve different models, of which each was replicated five times to allow for a solid statistical analysis of the influencing model (Table S7).

**Model selection** To determine the best model for scoring germination, we compared all models with the m$AP$@[.5:.95] score, which is a standard metric to evaluate a model's ability to accurately detect and classify objects. Since the models with the highest m$AP$@[.5:.95] scores varied in their classification errors, we developed a penalty approach to take these errors into account. Each image consisted of five ruler elements, one QR-code and ten seeds or seedlings. We gave a penalty of 1 for each element that was not found or misclassified. Since the penalty scoring could only be done manually and was therefore time-consuming, we performed it only on the four best models according to the m$AP$@[.5:.95] score (and on one intermediate model for comparison) and only on a randomly selected subset of 35 images. We selected both the model with the highest m$AP$@[.5:.95] score (66.39, model 11-D) and the model with the lowest penalty score (0.63 per image, model 8-C) for the final pipeline with which we analyzed all images from our seed germination experiment.

**Statistical analysis on model parameters** To investigate which parameters influenced the m$AP$@[.5:.95] score, a statistical analysis was performed using an ANOVA model with type III error from the R package `car` (John & Sanford, 2019), in which all parameters (`loss`, `heads.m`, `mask_resolution`) and their 2-way and 3-way interactions were included. Model selection was carried out base d on significance of $p<0.05$ dropping non-significant factors. Least square means were calculated on significant factors using the R package `emmeans` (Russell V., 2020).

**Pipeline set up and running** We developed a Python pipeline (python version 3.7) to automatically detect seeds, seedlings, ruler elements and QR-code on each image, to extract the surface area of each object in pixel and to calculate the surface area in $cm^2$ of each seed and seedling using the ruler elements as standard. On average, the analysis took 12.5 seconds per image on a 16-kernel CPU workstation.

**Detection evaluation of models 8-C and 11-D** We employed several approaches to evaluate the detection of the different objects per image in order to confirm the quality of the outcome from the two models 8-C and 11-D. First, we used the extracted surface area of each object to identify misclassifications. Our rationale behind this was that any misclassified object would not match with the other objects of its class. Second, we counted the objects per image because any image should contain exactly 16 objects: one QR-code, five ruler elements, and a total of ten seeds and seedlings. Taken together our findings suggested that model 8-C was more reliable (see Results) and therefore we discarded model 11-D at this step and only used the output of 8-C for the analyses. Even though model 8-C was clearly superior to model 11-D, it still did not score all images 100% correctly (see Results). We therefore included only images where at most ten seeds and/or seedlings were detected into our analysis.

**Evaluation with other images** We investigated whether our trained model 8-C is also applicable to other germination images than the ones taken during our seed germination experiment (e.g., different sized Petri dishes, different number of seeds, and different underground). We tested it on a set of images taken from germinations of quinoa seeds on smaller Petri dishes (diameter 3.2 cm), with either a low fixed seed number (five) or high and variable seed numbers (ten to 30), and a different underground (light green, uneven (Figure S8e). We first applied the trained model as is to 20 validation images (ten with low and ten with high seed numbers) without any update. Then we updated the model by training it with 30 training images of this other set (20 with low and ten with high seed numbers). We chose to update the model 8-C on either ten, 30 or 50 epochs with each three replicates using the best hyper-parameter set resulting from the statistical analysis which were trained of `all` layers in `heads.m` and the `mask1class10` loss. Then we used our previously developed penalty approach (see above) to evaluate the validation images and thereby identify the best updated model. Finally, we analyzed a random subset of 20 images with the best updated model (which was replicate 2 with 30 epochs) and applied again our previously developed penalty approach for a final assessment of the pipeline.
