## Supplementary material for "Phenotypic differentiation between highland and coastal quinoa under cold stress conditions": README.md

- **Data S1** – Multi-trait stability index (MTSI) values for the evaluated genotypes.
- **Data S2** – Detailed information of the 60 accessions included in this study.
- **Data S3** – Field data for individual plant trait measurements.
- **Data S4** – Field data for trait values averaged at the plot level.
- **Data S5** – Laboratory germination data for all 60 accessions.
- **Data S6** – Laboratory germination data for all 60 accessions derived from image analysis.
- **Data S7** – MARViN dataset for seeds from three successive quinoa generations (G₀, G₁, and G₂).
- **Data S8** – Image analysis output corresponding to model 8-C.
- **Data S9** – Image analysis output corresponding to model 11-D.
