## Supplementary material for "Phenotypic differentiation between highland and coastal quinoa under cold stress conditions": R Scripts: Paper_analysis.R

```
library(agricolae)
library(xlsx)
library(tidyverse)
library(ggplot2)
library(GGally)
library(grid)
library(gridExtra)
library(ggpubr)
library(ggsignif)
library(ggthemes)
library(ggsci)
library(magrittr)
library(dplyr)
library(FactoMineR)
library(Rmisc)
library(lsmeans)
library(metan)
library(Hmisc)
library(factoextra)
library(RColorBrewer)
library(corrplot)
library(PerformanceAnalytics)
library(quantmod)
library(wesanderson)
library(xlsx)
library(readODS)
library(tidyverse)
library(dplyr)
library(purrr)
library(asreml)
library(patchwork)
library(cowplot)
#FIELD EXPERIMENT------------------------------------------------------------------------------
#1 PCA analysis--------------------------------------------------------------
###1.1 Genotypic PCA-------------------------------------------------------

install.packages("BiocManager")
BiocManager::install("SNPRelate")
sBiocManager::install("LEA")
force=TRUE
library("SNPRelate") 
library("LEA")
library("vcfR")
rm("vcf")
vcf <- c("quinoa_field2021_miss0.80_pruned_plink2.vcf")

system.time(snpgdsVCF2GDS(vcf, "quinoa_60field.gds", method="biallelic.only"))
snpgdsSummary("quinoa_60field.gds")
genofile <- snpgdsOpen("quinoa_60field.gds")
#snpset <- snpgdsLDpruning(genofile, ld.threshold=0.2)


pca <- snpgdsPCA(genofile, autosome.only=FALSE)
pc.percent <- pca$varprop*100
percentages <- round(pc.percent, 2)
percentages

table <- data.frame(sample.id = pca$sample.id, # make data frame
                    PC1 = pca$eigenvect[,1],   # 1st principal component
                    PC2 = pca$eigenvect[,2],   # 2nd principal component   # 3rd principal component
                    stringsAsFactors = FALSE)

origin_data<-read.csv("acc.csv",header = TRUE)#merging with the accession data
origin_data$ID <- gsub("H21_", "", origin_data$ID)
m1 <- origin_data[,c(1,3,4,5,6)] # Merging with the passport data
table <- merge(table, m1, by.x = "sample.id", by.y = "Taxa", all.x = TRUE)
view(table)

theme<-theme(panel.background = element_blank(),panel.border=element_rect(fill=NA),
             panel.grid.major = element_blank(),panel.grid.minor = element_blank(),
             strip.background=element_blank(),axis.text.x=element_text(colour="black"),
             axis.text.y=element_text(colour="black"),axis.ticks=element_line(colour="black"),
             plot.margin=unit(c(1,1,1,1),"line"))
nudge <- position_nudge(y = 1)
p_pca<-ggplot(table,aes(x=PC1,y=PC2,color=Origin,shape=Type,
                        frame=TRUE,frame.color="Type",label=TRUE,
                        addEllipses=TRUE, ellipse.level=0.95))+
  geom_point(alpha=0.8, size=7)+ 
  scale_color_manual(values = c("Bolivia" = "#E46C5A",  
                                "Chile" = "#57C3C2",  
                                "Peru" = "#199674",  
                                "USA" = "#485D7F"))+
  labs(x="PC 1 (8.2%)",y="PC 2 (6.0%)",shape="Ecotype")+
  theme_bw()+
  theme(axis.text.x = element_text( size=12),
        axis.text.y = element_text( size=12),
        axis.title.x = element_text( size = 16, face = "bold"),
        axis.title.y = element_text( size = 16, face = "bold"),
        legend.title = element_text( face= "bold",size = 14),
        legend.text = element_text(size=12)
  )

p_pca ## Figure 1

ggsave(plot = p_pca,  dpi = 600, filename = "Fig_1.png")

## data_file1 consists of data except flowering, maturity and seed yield.
## data_file2 consits of data of flowering, maturity and seed yield
##acc file consists of accessions data
##Tag_103 file consists of temperature data

##Data loading -----------------------------------------------------------------

data <- read.csv("fielddata_file1.csv", header = TRUE)
str(data)
summary(data)
data$X <- NULL
data$ID <- gsub("H21_", "", data$ID)
data$ID <- as.factor(as.character(data$ID))
data$Rep <- as.factor(as.character(data$Rep))
data$Row <- as.factor(as.character(data$Row))
data$Col <- as.factor(as.character(data$Col))
data$Plants <- as.factor(as.character(data$Plants))
data$Time_Point <- as.factor(as.character(data$Time_Point))
data$Plot_ID<-as.factor(as.character(data$Plot_ID))


#origin_data<-read.csv("acc.csv",header = TRUE)
#origin_data$ID <- gsub("H21_", "", origin_data$ID)

#2. GDD calculation  -----------------------------------------------------------------------

data_t1 <- filter(data, Time_Point == "T1")
data_t2 <- filter(data, Time_Point == "T2")
data_t3 <- filter(data, Time_Point == "T3")

data_temp <- read.csv("Temperature_field.csv", header = TRUE)

data_temp$GDD_Ger <-  ifelse(data_temp$GDD_Ger <= 0, 0, data_temp$GDD_Ger)
data_temp$GDD <-  ifelse(data_temp$GDD <= 0, 0, data_temp$GDD)

data_temp <- data_temp %>%
  group_by(Time_Point)%>%
  mutate(Ac_GDD_ger = cumsum(GDD_Ger),
         Ac_GDD_Pemr = cumsum(GDD),
         Ac_GDD_fl = cumsum(GDD),
         Ac_GDD_mt = cumsum(GDD))
####i. merging GDD values in T1 ---------------------------

data_ger_t1 <- data_temp %>%
  filter(Time_Point == "T1")%>%
  select(c(11,12,14,16))

data_pem_t1 <- data_temp %>%
  filter(Time_Point == "T1")%>%
  select(c(11,12,15,17))

data_t1 <- merge(data_t1, data_ger_t1, 
                 by.x = "Germination_DAS",by.y = "DAS", all.x = TRUE)
data_t1 <- merge(data_t1, data_pem_t1, 
                 by.x = "Panicle_Emergence",by.y = "DAS", all.x = TRUE)

data_t1 <- data_t1 %>%
  select(-c(13,16))%>%
  relocate(Panicle_Emergence, .after = Plants)%>%
  relocate(Germination_DAS, .after = Plants)

####ii. merging GDD values in T2 -------------------------------

data_ger_t2 <- data_temp %>%
  filter(Time_Point == "T2")%>%
  select(c(11,12,14,16))

data_pem_t2 <- data_temp %>%
  filter(Time_Point == "T2")%>%
  select(c(11,12,15,17))

data_t2 <- merge(data_t2, data_ger_t2, 
                 by.x = "Germination_DAS",by.y = "DAS", all.x = TRUE)
data_t2 <- merge(data_t2, data_pem_t2, 
                 by.x = "Panicle_Emergence",by.y = "DAS", all.x = TRUE)

data_t2 <- data_t2 %>%
  select(-c(13,16))%>%
  relocate(Panicle_Emergence, .after = Plants)%>%
  relocate(Germination_DAS, .after = Plants)

####iii. merging GDD values in T3 ---------------------------------

data_ger_t3 <- data_temp %>%
  filter(Time_Point == "T3")%>%
  select(c(11,12,14,16))

data_pem_t3 <- data_temp %>%
  filter(Time_Point == "T3")%>%
  select(c(11,12,15,17))

data_t3 <- merge(data_t3, data_ger_t3, 
                 by.x = "Germination_DAS",by.y = "DAS", all.x = TRUE)
data_t3 <- merge(data_t3, data_pem_t3, 
                 by.x = "Panicle_Emergence",by.y = "DAS", all.x = TRUE)

data_t3 <- data_t3 %>%
  select(-c(13,16))%>%
  relocate(Panicle_Emergence, .after = Plants)%>%
  relocate(Germination_DAS, .after = Plants)

#3 Data per plant and plot
#3.1 Complete data with 10 plants per plot --------------------------------------------------

data <- rbind(data_t1, data_t2, data_t3)
data<- data%>%
  rename(Time_Point= Time_Point.x)
data <- merge(data, origin_data, by= "ID", all.x = TRUE)
write.csv(data,"data_PerPlant.csv", na = "NA")

#3.2 Data per plot -------------------------------------------------------
# averaged for each plot 
avg_data <- data %>%
  group_by(ID, Rep, Time_Point, Plot_ID) %>%
  summarise_at(vars(Germination_DAS:Panicle_Width, Ac_GDD_ger, Ac_GDD_Pemr),
               .funs = mean,
               na.rm = TRUE)
##i. calculating germination percentage --------------------------------
perc <- data %>%
  dplyr::group_by(ID,Rep,Time_Point) %>%
  dplyr::summarise(Germination_Perc = sum(!is.na(Germination_DAS))/n()*100)
avg_data <- merge(avg_data, perc, by = c("ID", "Rep", "Time_Point"))
avg_data$Germination_Perc <- 
  ifelse(avg_data$Germination_Perc == 0 , NA, avg_data$Germination_Perc)

str(avg_data)

##ii. merging Flowering, maturity, and seed yield per plot -------------------------------
data2<-read.csv("fielddata_file2.csv", header = TRUE)
data2$X <- NULL
data2$ID <- as.factor(as.character(data2$ID))
data2$Rep <- as.factor(as.character(data2$Rep))
data2$Time_Point <- as.factor(as.character(data2$Time_Point))
data2$Plot_ID<-as.factor(as.character(data2$Plot_ID))

avg_data <- merge(avg_data, data2, by = c("ID","Rep","Time_Point","Plot_ID"))  

avg_data[which(avg_data == 0)] <- NA 
avg_data[avg_data == "NaN"] <- NA

#3.3 GDD for flowering and maturity  ------------------------------------------------------------------------

avg_data_t1 <- filter(avg_data, Time_Point == "T1")
avg_data_t2 <- filter(avg_data, Time_Point == "T2")
avg_data_t3 <- filter(avg_data, Time_Point == "T3")

##i. merging GDD in T1 ---------------------------------------
data_fl_t1 <- data_temp %>%
  filter(Time_Point == "T1") %>%
  dplyr::select(11,12,15,18)

data_mt_t1 <- data_temp %>%
  filter(Time_Point == "T1") %>%
  dplyr::select(11,12,19)


avg_data_t1 <- merge(avg_data_t1, data_fl_t1, 
                     by.x = "Flowering",by.y = "DAS", all.x = TRUE)
avg_data_t1 <- merge(avg_data_t1, data_mt_t1, 
                     by.x = "Maturity",by.y = "DAS", all.x = TRUE)

avg_data_t1 <- avg_data_t1 %>%
  dplyr::select(-c(16,19))%>%
  relocate(Flowering, .after = Panicle_Emergence)%>%
  relocate(Maturity, .after = Flowering)


##ii.merging GDD in T2 ------------------------------------------------

data_fl_t2 <- data_temp %>%
  filter(Time_Point == "T2") %>%
  dplyr::select(11,12,15,18)

data_mt_t2 <- data_temp %>%
  filter(Time_Point == "T2") %>%
  dplyr::select(11,12,19)


avg_data_t2 <- merge(avg_data_t2, data_fl_t2, 
                     by.x = "Flowering",by.y = "DAS", all.x = TRUE)
avg_data_t2 <- merge(avg_data_t2, data_mt_t2, 
                     by.x = "Maturity",by.y = "DAS", all.x = TRUE)

avg_data_t2 <- avg_data_t2 %>%
  dplyr::select(-c(16,19))%>%
  relocate(Flowering, .after = Panicle_Emergence)%>%
  relocate(Maturity, .after = Flowering)

##iii. merging GDD T3 --------------------------------------------------
data_fl_t3 <- data_temp %>%
  filter(Time_Point == "T3") %>%
  dplyr::select(11,12,15,18)

data_mt_t3 <- data_temp %>%
  filter(Time_Point == "T3") %>%
  dplyr::select(11,12,19)


avg_data_t3 <- merge(avg_data_t3, data_fl_t3, 
                     by.x = "Flowering",by.y = "DAS", all.x = TRUE)
avg_data_t3 <- merge(avg_data_t3, data_mt_t3, 
                     by.x = "Maturity",by.y = "DAS", all.x = TRUE)

avg_data_t3 <- avg_data_t3 %>%
  dplyr::select(-c(16,19))%>%
  relocate(Flowering, .after = Panicle_Emergence)%>%
  relocate(Maturity, .after = Flowering)


avg_data <- rbind(avg_data_t1, avg_data_t2, avg_data_t3)
avg_data <- merge(avg_data,origin_data, by = "ID", all.x = TRUE)
avg_data <- relocate(avg_data, Germination_Perc, .after = Germination_DAS)
avg_data<- avg_data%>%
  relocate(Seed_Yield, .after = Maturity)
### creating row and column data --------------------------------------
data_r <- data[c(3,4,5)]
data_r <- data_r %>%
  distinct(Plot_ID, .keep_all = TRUE)
avg_data<-merge(avg_data,data_r,by="Plot_ID",all.x = TRUE)
avg_data <- relocate(avg_data, Row,Col,Plot_ID, .after = ID)

avg_data<- avg_data%>% rename(Time_Point=Time_Point.x)

#3.4 Final data ---------------------------------------
write.csv(avg_data,"data_PerPlot.csv", na = "NA") # data used for all statistical analysis

#4.Statistical Analysis---------------------------------------------------------


avg_data$ID <- as.factor(as.character(avg_data$ID))
avg_data$Rep <- as.factor(as.character(avg_data$Rep))
avg_data$Time_Point <- as.factor(as.character(avg_data$Time_Point))
avg_data$Plot_ID<-as.factor(as.character(avg_data$Plot_ID))
avg_data$Accession<-as.factor(as.character(avg_data$Accession))
avg_data$Type<-as.factor(as.character(avg_data$Type))
avg_data$Origin<-as.factor(as.character(avg_data$Origin))
avg_data$Germination_Perc<-as.numeric(as.character(avg_data$Germination_Perc))
avg_data$Flowering<-as.numeric(as.character(avg_data$Flowering))
avg_data$Maturity<-as.numeric(as.character(avg_data$Maturity))
##i. data ordering for residual factor in model-------------------------------------------------------------------

avg_data <- avg_data[with(avg_data,order(Plot_ID,Time_Point)),]

#4.1 BLUPs calculation-----------------------------------------------------------------

traits <- names(avg_data[c(7:17,19,20)])

traits

var <- ID <- ID_T <- ID_R_T <-list(traits)

str(avg_data$Ac_GDD_Pemr)

avg_data <- droplevels(avg_data)

for (i in traits)
{
  mod_blup <- asreml(as.formula(paste(i, "~ Rep + Time_Point+ Rep:Time_Point")),
                     random = ~ID + ID:Time_Point,
                     residual= ~ Plot_ID:ar1(Time_Point),
                     data = avg_data)
  
  pred_blups_ID <- predict(mod_blup, classify = "ID")
  pred_blups_IDxT <- predict(mod_blup, classify = "ID:Time_Point")
  pred_blups_IDxRxT <- predict(mod_blup, classify = "ID:Rep:Time_Point")
  
  bp_ID <- pred_blups_ID$pvals
  bp_ID <- bp_ID %>%
    dplyr::select(c(1:2))
  
  bp_IDxT <- pred_blups_IDxT$pvals
  bp_IDxT <- bp_IDxT %>%
    dplyr::select(c(1:3))
  
  bp_IDxRxT <- pred_blups_IDxRxT$pvals
  bp_IDxRxT <- bp_IDxRxT %>%
    dplyr::select(c(1:4))
  
  Variance <- as.data.frame(summary(mod_blup)$varcomp)
  Variance <- Variance %>%
    mutate(avsed = pred_blups_ID$avsed,
           h2 = c(1-(((pred_blups_ID$avsed)^2)/(2*(Variance[1,1])))))
  
  var[[paste(i)]] <- Variance
  ID[[paste(i)]] <- bp_ID
  ID_T[[paste(i)]] <- bp_IDxT
  ID_R_T[[paste(i)]] <- bp_IDxRxT
  
}

Ran_var <- do.call(rbind.data.frame, var)
Ran_var <- Ran_var[-1,]

Ran_var$Factor <- rownames(Ran_var)
Ran_var <- Ran_var %>%
  remove_rownames()%>%
  relocate(Factor, .before = component) %>%
  separate(Factor, c("Trait", "Factor"), sep = "([.])", extra = "merge")%>%
  mutate_if(is.numeric, round, digits = 2)

Var_h2 <- Ran_var %>%
  distinct(Trait, .keep_all = TRUE)

ID <- ID[-1]

Blups_ID <- cbind(ID$Germination_DAS, ID$Germination_Perc$predicted.value, 
                  ID$Panicle_Emergence$predicted.value,ID$Flowering$predicted.value,
                  ID$Maturity$predicted.value,
                  ID$Seed_Yield$predicted.value, ID$Plant_Height$predicted.value,
                  ID$Panicle_Length$predicted.value, ID$Panicle_Width$predicted.value,
                  ID$Ac_GDD_ger$predicted.value, ID$Ac_GDD_Pemr$predicted.value,
                  ID$Ac_GDD_fl$predicted.value, ID$Ac_GDD_mt$predicted.value)

names(Blups_ID)[2:14] <- traits

traits
Blups_ID <- merge(Blups_ID, origin_data, by = "ID")

write.csv(Blups_ID,"Blups_ID.csv", na="NA")

ID_T <- ID_T[-1]

Blups_ID_T <- cbind(ID_T$Germination_DAS, ID_T$Germination_Perc$predicted.value, 
                    ID_T$Panicle_Emergence$predicted.value, ID_T$Flowering$predicted.value, 
                    ID_T$Maturity$predicted.value,
                    ID_T$Seed_Yield$predicted.value, ID_T$Plant_Height$predicted.value,
                    ID_T$Panicle_Length$predicted.value, ID_T$Panicle_Width$predicted.value,
                    ID_T$Ac_GDD_ger$predicted.value, ID_T$Ac_GDD_Pemr$predicted.value,
                    ID_T$Ac_GDD_fl$predicted.value, ID_T$Ac_GDD_mt$predicted.value)

names(Blups_ID_T)[3:15] <- traits

Blups_ID_T <- merge(Blups_ID_T, origin_data, by = "ID")

ID_R_T <- ID_R_T[-1]

Blups_ID_R_T <- cbind(ID_R_T$Germination_DAS, ID_R_T$Germination_Perc$predicted.value, 
                      ID_R_T$Panicle_Emergence$predicted.value, ID_R_T$Flowering$predicted.value,
                      ID_R_T$Maturity$predicted.value,
                      ID_R_T$Seed_Yield$predicted.value,ID_R_T$Plant_Height$predicted.value,
                      ID_R_T$Panicle_Length$predicted.value, ID_R_T$Panicle_Width$predicted.value,
                      ID_R_T$Ac_GDD_ger$predicted.value, ID_R_T$Ac_GDD_Pemr$predicted.value,
                      ID_R_T$Ac_GDD_fl$predicted.value, ID_R_T$Ac_GDD_mt$predicted.value)

names(Blups_ID_R_T)[4:16] <- traits

Blups_ID_R_T <- merge(Blups_ID_R_T, origin_data, by = "ID")

##i. Heritability output-calculation of heritability is including in the above loop ----------------------------------------------------------
Var_h2$component<-as.numeric(as.character(Var_h2$component))
Var_h2$avsed<-as.numeric(as.character(Var_h2$avsed))
Var_h2$h2<-as.numeric(as.character(Var_h2$h2))

Var_h2<- Var_h2%>%
  mutate_if(is.numeric,round,digits=2)

#4.2 BLUEs Calculation------------------------------------------------------------
traits <- names(avg_data[c(7:17,19,20)])

an <- ID <- ID_T <- ID_R_T <-list(traits)

avg_data <- droplevels(avg_data)

for (i in traits)
{
  mod_blue <- asreml(as.formula(paste(i, "~ Rep + ID + Time_Point+ ID:Time_Point+ Rep:Time_Point")),
                     residual= ~ Plot_ID:ar1(Time_Point),
                     data = avg_data)
  
  pred_blues_ID <- predict(mod_blue, classify = "ID")
  pred_blues_IDxT <- predict(mod_blue, classify = "ID:Time_Point")
  pred_blues_IDxRxT <- predict(mod_blue, classify = "ID:Rep:Time_Point")
  
  blues_ID <- pred_blues_ID$pvals
  blues_ID <- blues_ID %>%
    dplyr::select(c(1:2))
  
  blues_IDxT <- pred_blues_IDxT$pvals
  blues_IDxT <- blues_IDxT %>%
    dplyr::select(c(1:3))
  
  blues_IDxRxT <- pred_blues_IDxRxT$pvals
  blues_IDxRxT <- blues_IDxRxT %>%
    dplyr::select(c(1:4))
  
  #Variance <- as.data.frame(summary(mod_blue)$varcomp)
  anova <- as.data.frame(wald(mod_blue))
  
  an[[paste(i)]] <- anova
  ID[[paste(i)]] <- blues_ID
  ID_T[[paste(i)]] <- blues_IDxT
  ID_R_T[[paste(i)]] <- blues_IDxRxT
  
}

anv <- do.call(rbind.data.frame, an)
anv <- anv[-1,]

anv$Factor <- rownames(anv)
anv <- anv %>%
  remove_rownames()%>%
  relocate(Factor, .before = Df) %>%
  separate(Factor, c("Trait", "Factor"), sep = "([.])", extra = "merge")

ID <- ID[-1]

Blues_ID <- cbind(ID$Germination_DAS, ID$Germination_Perc$predicted.value, 
                  ID$Panicle_Emergence$predicted.value,ID$Flowering$predicted.value, 
                  ID$Maturity$predicted.value,
                  ID$Seed_Yield$predicted.value, ID$Plant_Height$predicted.value,
                  ID$Panicle_Length$predicted.value, ID$Panicle_Width$predicted.value,
                  ID$Ac_GDD_ger$predicted.value, ID$Ac_GDD_Pemr$predicted.value,
                  ID$Ac_GDD_fl$predicted.value, ID$Ac_GDD_mt$predicted.value)

names(Blues_ID)[2:14] <- traits

Blues_ID <- merge(Blues_ID, origin_data, by = "ID")

ID_T <- ID_T[-1]

Blues_ID_T <- cbind(ID_T$Germination_DAS, ID_T$Germination_Perc$predicted.value, 
                    ID_T$Panicle_Emergence$predicted.value,ID_T$Flowering$predicted.value,
                    ID_T$Maturity$predicted.value,
                    ID_T$Seed_Yield$predicted.value, ID_T$Plant_Height$predicted.value,
                    ID_T$Panicle_Length$predicted.value, ID_T$Panicle_Width$predicted.value,
                    ID_T$Ac_GDD_ger$predicted.value, ID_T$Ac_GDD_Pemr$predicted.value,
                    ID_T$Ac_GDD_fl$predicted.value, ID_T$Ac_GDD_mt$predicted.value)

names(Blues_ID_T)[3:15] <- traits

Blues_ID_T <- merge(Blues_ID_T, origin_data, by = "ID")

ID_R_T <- ID_R_T[-1]

Blues_ID_R_T <- cbind(ID_R_T$Germination_DAS, ID_R_T$Germination_Perc$predicted.value, 
                      ID_R_T$Panicle_Emergence$predicted.value,ID_R_T$Flowering$predicted.value,
                      ID_R_T$Maturity$predicted.value,
                      ID_R_T$Seed_Yield$predicted.value, ID_R_T$Plant_Height$predicted.value,
                      ID_R_T$Panicle_Length$predicted.value, ID_R_T$Panicle_Width$predicted.value,
                      ID_R_T$Ac_GDD_ger$predicted.value, ID_R_T$Ac_GDD_Pemr$predicted.value,
                      ID_R_T$Ac_GDD_fl$predicted.value, ID_R_T$Ac_GDD_mt$predicted.value)

names(Blues_ID_R_T)[4:16] <- traits

Blues_ID_R_T <- merge(Blues_ID_R_T, origin_data, by = "ID")

Blues_ID_R_T$Germination_Perc <- ifelse(Blues_ID_R_T$Germination_Perc <= 0, NA, Blues_ID_R_T$Germination_Perc)

###4.2.1 Wald test results extraction (Table 1)-----------------------------------------------
str(anv)
anv$`Pr(Chisq)`<-as.numeric(anv$`Pr(Chisq)`)
anv$`Pr(Chisq)` <- round(anv$`Pr(Chisq)`, digits = 3)

stars <- ifelse(anv$`Pr(Chisq)` < .0001, "****",
                ifelse(anv$`Pr(Chisq)` < .001, "***",
                       ifelse(anv$`Pr(Chisq)` < .01, "**",
                              ifelse(anv$`Pr(Chisq)` < .05, "*", ""))))


test_anv <- anv
test_anv$`Pr(Chisq)` <- paste(test_anv$`Pr(Chisq)`, stars, "")
write.csv(test_anv, "Wald_results_field.csv", na="NA" ) # Table 4

## The codes provided here are in the sequence same as results section in the main paper.
# Wald-test results were provided in section 4.2.1
#5. Results---------------------------------------------------------------------
##The following datasets consists of the output of BLUEs
Blues_ID_R_T<- read.csv("Blues_ID_R_T.csv", header=TRUE)
Blues_ID_R_T <- Blues_ID_R_T[,-1]
Blues_ID_T<- read.csv("Blues_ID_T.csv", header=TRUE)
Blues_ID_T <- Blues_ID_T[,-1]
Blues_ID<- read.csv("Blues_ID.csv", header=TRUE)
Blues_ID <- Blues_ID[,-1]

## Figure_2a Time points (Growth stages)-------------------------------------------------------------------
HSD_ger<-  HSD.test(aov(Germination_DAS~Time_Point,data=Blues_ID_R_T),"Time_Point",group=T)$groups
HSD_ger<-as.data.frame(HSD_ger)
HSD_ger$Time_Point<-rownames(HSD_ger)
HSD_ger <- HSD_ger %>%
  remove_rownames()%>%
  relocate(Time_Point, .before = Germination_DAS)
HSD_pe<-  HSD.test(aov(Panicle_Emergence~Time_Point,data=Blues_ID_R_T),"Time_Point",group=T)$groups
HSD_pe<-as.data.frame(HSD_pe)
HSD_pe$Time_Point<-rownames(HSD_pe)
HSD_pe<-HSD_pe%>%
  remove_rownames()%>%
  relocate(Time_Point, .before= Panicle_Emergence)
HSD_fl<-HSD.test(aov(Flowering~Time_Point,data=Blues_ID_R_T),"Time_Point",group=T)$groups
HSD_fl<-as.data.frame(HSD_fl)
HSD_fl$Time_Point<-rownames(HSD_fl)
HSD_fl<-HSD_fl%>%
  remove_rownames()%>%
  relocate(Time_Point, .before= Flowering)
HSD_mt<-HSD.test(aov(Maturity~Time_Point,data=Blues_ID_R_T),"Time_Point",group=T)$groups
HSD_mt<-as.data.frame(HSD_mt)
HSD_mt$Time_Point<-rownames(HSD_mt)
HSD_mt<-HSD_mt%>%
  remove_rownames()%>%
  relocate(Time_Point, .before= Maturity)

HSD<-merge(HSD_ger,HSD_pe,by=c("Time_Point"),all.x =TRUE)
HSD<-merge(HSD,HSD_fl,by=c("Time_Point"),all.x =TRUE)
HSD<-merge(HSD,HSD_mt,by=c("Time_Point"),all.x=TRUE)

HSD<-HSD[-c(3,5,7,9)]

HSD_diff <- HSD %>%
  mutate(diff_pe = Panicle_Emergence - Germination_DAS,
         diff_fl = Flowering - Panicle_Emergence,
         diff_mt = Maturity - Flowering)
HSD_diff <- HSD_diff[-c(3:5)]

HSD_diff<-HSD_diff%>%
  relocate(diff_pe, .before= Germination_DAS) %>%
  relocate(diff_fl, .before= diff_pe) %>%
  relocate(diff_mt, .before= diff_fl)

HSD_diff <- gather(HSD_diff, Growth_Stage, DAS, diff_mt:Germination_DAS, factor_key=TRUE)

Fig_DAS_T<-ggplot(HSD_diff, aes(fill=Growth_Stage, x=DAS, y=Time_Point))+
  geom_bar(width = 0.3, position="stack",stat="identity")+
  labs(y = "Sowing date", x = " Days after sowing (DAS)", tag="(a)")+
  scale_y_discrete(labels= c("T1"="S1","T2"="S2","T3"="S3"))+
  scale_x_continuous(limits = c(0,250),
                     breaks = seq(0,240,20), expand = c(0,0))+
  scale_fill_manual(values=wes_palette("Zissou1"),name = "Growth stage", 
                    labels = c("Maturity", "Flowering",
                               "Panicle Emergence","Seedling emergence"))+
  theme_bw()+theme(
    axis.title.x = element_text( size = 14, face = "bold"),
    plot.tag=element_text(face="bold"),
    axis.text.x = element_text(size=12, colour = "black"),
    axis.text.y = element_text(size=12, colour = "black"),
    axis.title.y = element_text( size = 14, face = "bold"),
    legend.position="top",legend.title=element_text(face = "bold",size=14), 
    legend.text=element_text(size=12))

Fig_DAS_T
ggsave(plot =Fig_DAS_T,width= 10.77, height=5.95, dpi = 600, filename = "Fig_DAS_T.png")
## Figure_2b GDD graphs (growth stages)------------------------------------------------------------------
GDD_data<- Blues_ID_T[c(1,2,12:15,16,18)]


HSD_Ac_ger<-  HSD.test(aov(Ac_GDD_ger~Time_Point:Type,data=GDD_data),"Time_Point",group=T)$groups
HSD_Ac_ger<-as.data.frame(HSD_Ac_ger)
HSD_Ac_ger$Time_Point<-rownames(HSD_Ac_ger)
HSD_Ac_ger <- HSD_Ac_ger %>%
  remove_rownames()%>%
  relocate(Time_Point, .before = Ac_GDD_ger)

HSD_Ac_pe<-  HSD.test(aov(Ac_GDD_Pemr~Time_Point,data=GDD_data),"Time_Point",group=T)$groups
HSD_Ac_pe<-as.data.frame(HSD_Ac_pe)
HSD_Ac_pe$Time_Point<-rownames(HSD_Ac_pe)
HSD_Ac_pe <- HSD_Ac_pe %>%
  remove_rownames()%>%
  relocate(Time_Point, .before = Ac_GDD_Pemr)

HSD_Ac_fl<-  HSD.test(aov(Ac_GDD_fl~Time_Point,data=GDD_data),"Time_Point",group=T)$groups
HSD_Ac_fl<-as.data.frame(HSD_Ac_fl)
HSD_Ac_fl$Time_Point<-rownames(HSD_Ac_fl)
HSD_Ac_fl <- HSD_Ac_fl %>%
  remove_rownames()%>%
  relocate(Time_Point, .before = Ac_GDD_fl)

HSD_Ac_mt<-  HSD.test(aov(Ac_GDD_mt~Time_Point,data=GDD_data),"Time_Point",group=T)$groups
HSD_Ac_mt<-as.data.frame(HSD_Ac_mt)
HSD_Ac_mt$Time_Point<-rownames(HSD_Ac_mt)
HSD_Ac_mt <- HSD_Ac_mt%>%
  remove_rownames()%>%
  relocate(Time_Point, .before = Ac_GDD_mt)

HSD_GDD<-merge(HSD_Ac_ger,HSD_Ac_pe,by=c("Time_Point"),all.x =TRUE)
HSD_GDD<-merge(HSD_GDD,HSD_Ac_fl,by=c("Time_Point"),all=TRUE)
HSD_GDD<-merge(HSD_GDD,HSD_Ac_mt,by=c("Time_Point"),all=TRUE)

HSD_GDD<-HSD_GDD[-c(3,5,7,9)]

HSD_GDD<- gather(HSD_GDD, Growth_stage,Acc_GDD, Ac_GDD_ger:Ac_GDD_mt, factor_key =TRUE )

Fig_GDD_T<-ggplot(HSD_GDD,aes(group=Time_Point, y=Growth_stage, x=Acc_GDD, shape = Time_Point))+
  geom_line(aes(color=Time_Point))+
  geom_point(aes(color=Time_Point),size=3)+
  scale_shape_manual(values = c(16, 17, 15),name = "Sowing date", 
                     labels = c("S1", "S2",
                                "S3"))+
  labs(y = "Growth stage", x = "GDD", tag="(b)")+
  scale_y_discrete(label=c("SE","PE","FL","MT"))+
  scale_color_aaas( name="Sowing date", labels= c("T1"="S1","T2"="S2","T3"="S3"))+ 
  scale_x_continuous(limits = c(0,2300),
                     breaks = seq(0,2300,200), expand = c(0,0))+
  theme_bw()+
  theme(axis.title.x = element_text(size = 14, face = "bold"),
        axis.text.x = element_text(size=12, colour="black"),
        axis.text.y = element_text(size=12, colour="black"),
        plot.tag=element_text(face="bold"),
        axis.title.y = element_text(size = 14, face = "bold"),
        legend.position = "bottom",
        legend.title=element_text(face = "bold",size=14), 
        legend.text=element_text(size=12))
Fig_GDD_T

Fig_2<- ggarrange(Fig_DAS_T,Fig_GDD_T, nrow=2)
Fig_2 ## Figure 2
ggsave(plot = Fig_2,width= 15.77, height=9.95, dpi = 600, filename = "Fig_2.png")

##Figure_3 comparisons across three sowing dates (S1, S2, and S3)-------------------------------------------------------------
my_comparisons = list( c("T1", "T2"), c("T2", "T3"), c("T1", "T3") )
Fig_FGP <- ggplot(Blues_ID_T,aes(x=Time_Point, y = Germination_Perc,color=Time_Point))+
  geom_boxplot()+
  stat_summary(fun = mean, geom = "crossbar",linetype="dotted",width=.75) +
  geom_jitter(position = position_jitterdodge(jitter.width = 0.3), alpha = 0.7)+
  labs(y = "Emergence percentage (%)", x = "Sowing date", tag="(a)",color="Sowing date")+
  stat_compare_means(comparisons = my_comparisons, label="p.signif",method="t.test", label.y = c(70, 80, 90))+
  scale_color_manual(values = c("deepskyblue", "deepskyblue4", "dodgerblue4"), 
                     labels = c("T1" = "S1", "T2" = "S2", "T3"="S3") )+
  scale_x_discrete(labels=c("S1", "S2","S3"))+
  scale_y_continuous(limits = c(0,100),
                     breaks = seq(0,100,10), expand = c(0,0))+
  #scale_color_discrete(labels = c("T1" = "S1", "T2" = "S2", "T3"="S3")) +
  theme_bw()+
  theme(axis.title = element_text(size = 14, face = "bold"),
        axis.text = element_text(size=12, colour="black"),
        legend.position = "none",
        plot.tag=element_text(face = "bold"),
        legend.title = element_text(face = "bold",size=14), 
        legend.text=element_text( size=12))

Fig_FGP
ggsave(plot = Fig_FGP,  dpi = 600, filename = "Fig_FGP.png")

Fig_SY <- ggplot(Blues_ID_T,aes(x=Time_Point, y = Seed_Yield,color=Time_Point))+
  geom_boxplot()+
  stat_summary(fun = mean, geom = "crossbar",linetype="dotted",width=.75) +
  geom_jitter(position = position_jitterdodge(jitter.width = 0.3), alpha = 0.7)+
  labs(y = "Seed yield (g)", x = "Sowing date", tag="(b)",color="Sowing date")+
  stat_compare_means(comparisons = my_comparisons, label="p.signif",method="t.test", label.y = c(180, 190, 200))+
  scale_color_manual(values = c("deepskyblue", "deepskyblue4", "dodgerblue4"), 
                     labels = c("T1" = "S1", "T2" = "S2", "T3"="S3") )+
  scale_x_discrete(labels=c("S1", "S2","S3"))+
  scale_y_continuous(limits = c(0,220),
                     breaks = seq(0,220,20), expand = c(0,0))+
  theme_bw()+
  theme(axis.title = element_text(size = 14, face = "bold"),
        axis.text = element_text(size=12, colour="black"),
        legend.position = "none",
        plot.tag=element_text(face = "bold"),
        legend.title = element_text(face = "bold",size=14), 
        legend.text=element_text(size=12))


Fig_SY

mean(Blues_ID_T$Seed_Yield[Blues_ID_T$Time_Point == "T3"], na.rm = TRUE)
ggsave(plot = Fig_SY,  dpi = 600, filename = "Fig_SY.png")

Fig_PH <- ggplot(Blues_ID_T,aes(x=Time_Point, y = Plant_Height,color=Time_Point))+
  geom_boxplot()+
  stat_summary(fun = mean, geom = "crossbar",linetype="dotted",width=.75) +
  geom_jitter(position = position_jitterdodge(jitter.width = 0.3), alpha = 0.7)+
  labs(y = "Plant height (cm)", x = "Sowing date", tag="(c)",color="Sowing date")+
  stat_compare_means(comparisons = my_comparisons, label="p.signif",method="t.test", label.y = c(220, 230, 240))+
  scale_color_manual(values = c("deepskyblue", "deepskyblue4", "dodgerblue4"), 
                     labels = c("T1" = "S1", "T2" = "S2", "T3"="S3") )+
  scale_x_discrete(labels=c("S1", "S2","S3"))+
  scale_y_continuous(limits = c(80,260),
                     breaks = seq(80,260,20), expand = c(0,0))+
  theme_bw()+
  theme(axis.title = element_text(size = 14, face = "bold"),
        axis.text = element_text(size=12, colour="black"),
        legend.position = "none",
        plot.tag=element_text(face = "bold"),
        legend.title = element_text(face = "bold",size=14), 
        legend.text=element_text( size=12))

Fig_PH
ggsave(plot = Fig_PH,  dpi = 600, filename = "Fig_PH.png")

Fig_PL <- ggplot(Blues_ID_T,aes(x=Time_Point, y = Panicle_Length,color=Time_Point))+
  geom_boxplot()+
  stat_summary(fun = mean, geom = "crossbar",linetype="dotted",width=.75) +
  geom_jitter(position = position_jitterdodge(jitter.width = 0.3), alpha = 0.7)+
  labs(y = "Panicle length (cm)", x = "Sowing date", tag="(d)",color="Sowing date")+
  stat_compare_means(comparisons = my_comparisons, label="p.signif",method="t.test", label.y = c(80, 85, 90))+
  scale_color_manual(values = c("deepskyblue", "deepskyblue4", "dodgerblue4"), 
                     labels = c("T1" = "S1", "T2" = "S2", "T3"="S3") )+
  scale_x_discrete(labels=c("S1", "S2","S3"))+
  scale_y_continuous(limits = c(0,100),
                     breaks = seq(0,100,10), expand = c(0,0))+
  theme_bw()+
  theme(axis.title = element_text(size = 14, face = "bold"),
        axis.text = element_text(size=12, colour="black"),
        legend.position = "none",
        plot.tag=element_text(face = "bold"),
        legend.title = element_text(face = "bold",size=14), 
        legend.text=element_text( size=12))

Fig_PL
ggsave(plot = Fig_PL,  dpi = 600, filename = "Fig_PL.png")

Fig_PW <- ggplot(Blues_ID_T,aes(x=Time_Point, y = Panicle_Width,color=Time_Point))+
  geom_boxplot()+
  stat_summary(fun = mean, geom = "crossbar",linetype="dotted",width=.75) +
  geom_jitter(position = position_jitterdodge(jitter.width = 0.3), alpha = 0.7)+
  labs(y = "Panicle width (cm)", x = "Sowing date", tag="(e)",color="Sowing date")+
  stat_compare_means(comparisons = my_comparisons, label="p.signif",method="t.test", label.y = c(12, 13, 14))+
  scale_color_manual(values = c("deepskyblue", "deepskyblue4", "dodgerblue4"), 
                     labels = c("T1" = "S1", "T2" = "S2", "T3"="S3") )+
  scale_x_discrete(labels=c("S1", "S2","S3"))+
  scale_y_continuous(limits = c(0,16),
                     breaks = seq(0,16,2), expand = c(0,0))+
  theme_bw()+
  theme(axis.title = element_text(size = 14, face = "bold"),
        axis.text = element_text(size=12, colour="black"),
        legend.position = "none",
        plot.tag=element_text(face = "bold"),
        legend.title = element_text(face = "bold",size=14), 
        legend.text=element_text( size=12))

Fig_PW
ggsave(plot = Fig_PW,  dpi = 600, filename = "Fig_PW.png")

legend <- get_legend(
  Fig_FGP + theme(legend.position = "bottom")
)

top_row <- plot_grid(Fig_FGP, Fig_SY, Fig_PH, ncol = 3, align = "v")

# Bottom row with 2 centered plots using rel_widths to shift them
bottom_row <- plot_grid(NULL, Fig_PL, Fig_PW, NULL, ncol = 4, rel_widths = c(0.5, 1, 1, 0.5))

# Combine top and bottom rows
main_plot <- plot_grid(top_row, bottom_row, ncol = 1, rel_heights = c(1, 0.8))

# Add the common legend below
Fig_Box_tp <- plot_grid(main_plot, legend, ncol = 1, rel_heights = c(4, 0.3))


Fig_Box_tp
ggsave(plot = Fig_Box_tp, width=15.93, height=9.96, dpi = 600, filename = "Fig_3.png")

##Figure_4 comparisons between Highland and Coastal within each Time_Point -------------------------------------
# Convert to factors
Blues_ID_T$Type <- factor(Blues_ID_T$Type, levels = c("Coastal","Highland"))
Blues_ID_T$Time_Point <- factor(Blues_ID_T$Time_Point)


###Germination DAS
# Perform pairwise comparisons
stat_test <- compare_means(
  Germination_DAS ~ Type,
  group.by = "Time_Point",
  data = Blues_ID_T,
  method = "t.test"
  #method="wilcox.test"
)

# Print results 
print(stat_test)

#  Manually calculate positions for lines and p-values

# 1. Define x-axis positions for dodged boxplots
dodge_width <- 0.8
time_points <- levels(Blues_ID_T$Time_Point)
n_timepoints <- length(time_points)

# Calculate x-coordinates for Highland (left) and Coastal (right) within each Time_Point
x_positions <- data.frame(
  Time_Point = rep(time_points, each = 2),
  Type = rep(c( "Coastal", "Highland"), times = n_timepoints),
  x = rep(1:n_timepoints, each = 2) + c(-dodge_width/2, dodge_width/2)
)

# 2. Define y_positions for horizontal lines (above the data)
y_max <- 32  # Based on your y-axis limit
y_start <- y_max - 2
y_step <- 2  # Spacing between lines
y_positions <- seq(y_start, y_start - (n_timepoints - 1) * y_step, by = -y_step)

# 3. Prepare p-value annotations with significance stars
stat_test$p_annot <- ifelse(
  stat_test$p < 0.001, "***",
  ifelse(stat_test$p < 0.01, "**",
         ifelse(stat_test$p < 0.05, "*", "ns"))
)

# Ensure stat_test is ordered to match Time_Point levels
stat_test <- stat_test[order(stat_test$Time_Point), ]


# 4. Create the plot
Fig_gr <- ggplot(Blues_ID_T, aes(x = Time_Point, y = Germination_DAS, color = Type)) +
  geom_boxplot(position = position_dodge(dodge_width)) +
  stat_summary(fun = mean, geom = "crossbar",linetype="dotted",width=.75,position = position_dodge(dodge_width)) +
  geom_jitter(position = position_jitterdodge(jitter.width = 0.3, dodge.width = dodge_width), alpha = 0.7) +
  labs(y = "Mean emergence time (DAS)", x = "Sowing date", tag="(a)",color = "Ecotype") +
  scale_color_manual(values = c("#B22222", "#3B4992FF") ) +
  scale_x_discrete(labels=c("S1", "S2","S3"))+
  scale_y_continuous(limits = c(0, 35), breaks = seq(0,35,2),expand = c(0, 0)) +
  theme_bw() +
  theme(
    axis.title.x = element_blank(),
    axis.text.x = element_text(size = 12, colour = "black"),
    axis.text.y = element_text(size = 12, colour = "black"),
    axis.title.y = element_text(size = 14, face = "bold",colour = "black"),
    plot.tag=element_text(face = "bold"),
    legend.title = element_text(face = "bold",size=14), 
    legend.text=element_text( size=12),
    legend.position = "bottom",
  ) +
  
  # Add horizontal lines and p-values manually
  geom_segment(
    data = data.frame(
      Time_Point = time_points,
      x_start = 1:n_timepoints - dodge_width/2,
      x_end = 1:n_timepoints + dodge_width/2,
      y = y_positions
    ),
    aes(x = x_start, xend = x_end, y = y, yend = y),
    color = "black",
    inherit.aes = FALSE
  ) +
  geom_text(
    data = data.frame(
      Time_Point = time_points,
      x = 1:n_timepoints,
      y = y_positions + 1,  # Slightly above the line
      label = paste("p =", stat_test$p.format, stat_test$p_annot)
    ),
    aes(x = x, y = y, label = label),
    color = "black",
    size = 3.5,
    inherit.aes = FALSE
  )
# Display the plot
print(Fig_gr)

###Germination Percentage
# Perform pairwise comparisons
stat_test <- compare_means(
  Germination_Perc ~ Type,
  group.by = "Time_Point",
  data = Blues_ID_T,
  method = "wilcox.test"
)

# Print results 
print(stat_test)

#  Manually calculate positions for lines and p-values

# 1. Define x-axis positions for dodged boxplots
dodge_width <- 0.8
time_points <- levels(Blues_ID_T$Time_Point)
n_timepoints <- length(time_points)

# Calculate x-coordinates for Highland (left) and Coastal (right) within each Time_Point
x_positions <- data.frame(
  Time_Point = rep(time_points, each = 2),
  Type = rep(c( "Coastal", "Highland"), times = n_timepoints),
  x = rep(1:n_timepoints, each = 2) + c(-dodge_width/2, dodge_width/2)
)

# 2. Define y_positions for horizontal lines (above the data)
y_max <- 90  # Based on your y-axis limit
y_start <- y_max - 2
y_step <- 2  # Spacing between lines
y_positions <- seq(y_start, y_start - (n_timepoints - 1) * y_step, by = -y_step)

# 3. Prepare p-value annotations with significance stars
stat_test$p_annot <- ifelse(
  stat_test$p < 0.001, "***",
  ifelse(stat_test$p < 0.01, "**",
         ifelse(stat_test$p < 0.05, "*", "ns"))
)

# Ensure stat_test is ordered to match Time_Point levels
stat_test <- stat_test[order(stat_test$Time_Point), ]

# 4. Create the plot
Fig_gp <- ggplot(Blues_ID_T, aes(x = Time_Point, y = Germination_Perc, color = Type)) +
  geom_boxplot(position = position_dodge(dodge_width)) +
  stat_summary(fun = mean, geom = "crossbar",linetype="dotted",width=.75,position = position_dodge(dodge_width)) +
  geom_jitter(position = position_jitterdodge(jitter.width = 0.3, dodge.width = dodge_width), alpha = 0.7) +
  labs(y = "Emergence percentage (%)", x = "Sowing date",tag="(b)", color = "Ecotype") +
  scale_color_manual(values = c("#B22222", "#3B4992FF") ) +
  scale_x_discrete(labels=c("S1", "S2","S3"))+
  scale_y_continuous(limits = c(0, 100), breaks = seq(0,100,10),expand = c(0, 0)) +
  theme_bw() +
  theme(
    axis.title.x = element_blank(),
    axis.text.x = element_text(size = 12, colour = "black"),
    axis.text.y = element_text(size = 12, colour = "black"),
    axis.title.y = element_text(size = 14, face = "bold"),
    plot.tag=element_text(face = "bold"),
    legend.title = element_text(face = "bold",size=14), 
    legend.text=element_text( size=12),
    legend.position = "bottom",
  ) +
  
  # Add horizontal lines and p-values manually
  geom_segment(
    data = data.frame(
      Time_Point = time_points,
      x_start = 1:n_timepoints - dodge_width/2,
      x_end = 1:n_timepoints + dodge_width/2,
      y = y_positions
    ),
    aes(x = x_start, xend = x_end, y = y, yend = y),
    color = "black",
    inherit.aes = FALSE
  ) +
  geom_text(
    data = data.frame(
      Time_Point = time_points,
      x = 1:n_timepoints,
      y = y_positions + 2.5,  # Slightly above the line
      label = paste("p =", stat_test$p.format, stat_test$p_annot)
    ),
    aes(x = x, y = y, label = label),
    color = "black",
    size = 3.5,
    inherit.aes = FALSE
  )
# Display the plot
print(Fig_gp)

###Panicle emergence
# Perform pairwise comparisons
stat_test <- compare_means(
  Panicle_Emergence ~ Type,
  group.by = "Time_Point",
  data = Blues_ID_T,
  method = "t.test"
)

# Print results 
print(stat_test)

#  Manually calculate positions for lines and p-values

# 1. Define x-axis positions for dodged boxplots
dodge_width <- 0.8
time_points <- levels(Blues_ID_T$Time_Point)
n_timepoints <- length(time_points)

# Calculate x-coordinates for Highland (left) and Coastal (right) within each Time_Point
x_positions <- data.frame(
  Time_Point = rep(time_points, each = 2),
  Type = rep(c( "Coastal", "Highland"), times = n_timepoints),
  x = rep(1:n_timepoints, each = 2) + c(-dodge_width/2, dodge_width/2)
)


# 2. Define y_positions for horizontal lines (above the data)
y_max <- 140  # Based on your y-axis limit
y_start <- y_max - 10
y_step <- 2  # Spacing between lines
y_positions <- seq(y_start, y_start - (n_timepoints - 1) * y_step, by = -y_step)

# 3. Prepare p-value annotations with significance stars
stat_test$p_annot <- ifelse(
  stat_test$p < 0.001, "***",
  ifelse(stat_test$p < 0.01, "**",
         ifelse(stat_test$p < 0.05, "*", "ns"))
)

# Ensure stat_test is ordered to match Time_Point levels
stat_test <- stat_test[order(stat_test$Time_Point), ]

# 4. Create the plot
Fig_pe <- ggplot(Blues_ID_T, aes(x = Time_Point, y =  Panicle_Emergence, color = Type)) +
  geom_boxplot(position = position_dodge(dodge_width)) +
  stat_summary(fun = mean, geom = "crossbar",linetype="dotted",width=.75,position = position_dodge(dodge_width)) +
  geom_jitter(position = position_jitterdodge(jitter.width = 0.3, dodge.width = dodge_width), alpha = 0.7) +
  labs(y = " Panicle emergence (DAS)", x = "Sowing date",tag="(c)", color = "Ecotype") +
  scale_color_manual(values = c("#B22222", "#3B4992FF") ) +
  scale_x_discrete(labels=c("S1", "S2","S3"))+
  scale_y_continuous(limits = c(0, 150), breaks = seq(0,150,20),expand = c(0, 0)) +
  theme_bw() +
  theme(
    axis.title.x = element_blank(),
    axis.text.x = element_text(size = 12, colour = "black"),
    axis.text.y = element_text(size = 12, colour = "black"),
    axis.title.y = element_text(size = 14, face = "bold"),
    plot.tag=element_text(face = "bold"),
    legend.title = element_text(face = "bold",size=14), 
    legend.text=element_text( size=12),
    legend.position = "bottom",
  ) +
  
  # Add horizontal lines and p-values manually
  geom_segment(
    data = data.frame(
      Time_Point = time_points,
      x_start = 1:n_timepoints - dodge_width/2,
      x_end = 1:n_timepoints + dodge_width/2,
      y = y_positions
    ),
    aes(x = x_start, xend = x_end, y = y, yend = y),
    color = "black",
    inherit.aes = FALSE
  ) +
  geom_text(
    data = data.frame(
      Time_Point = time_points,
      x = 1:n_timepoints,
      y = y_positions + 5,  # Slightly above the line
      label = paste("p =", stat_test$p.format, stat_test$p_annot)
    ),
    aes(x = x, y = y, label = label),
    color = "black",
    size = 3.5,
    inherit.aes = FALSE
  )
# Display the plot
print(Fig_pe)

###Days to flowering
# Perform pairwise comparisons
stat_test <- compare_means(
  Flowering ~ Type,
  group.by = "Time_Point",
  data = Blues_ID_T,
  method = "t.test"
)

# Print results 
print(stat_test)

#  Manually calculate positions for lines and p-values

# 1. Define x-axis positions for dodged boxplots
dodge_width <- 0.8
time_points <- levels(Blues_ID_T$Time_Point)
n_timepoints <- length(time_points)

# Calculate x-coordinates for Highland (left) and Coastal (right) within each Time_Point
x_positions <- data.frame(
  Time_Point = rep(time_points, each = 2),
  Type = rep(c( "Coastal", "Highland"), times = n_timepoints),
  x = rep(1:n_timepoints, each = 2) + c(-dodge_width/2, dodge_width/2)
)

# 2. Define y_positions for horizontal lines (above the data)
y_max <- 160  # Based on your y-axis limit
y_start <- y_max - 10
y_step <- 2  # Spacing between lines
y_positions <- seq(y_start, y_start - (n_timepoints - 1) * y_step, by = -y_step)

# 3. Prepare p-value annotations with significance stars
stat_test$p_annot <- ifelse(
  stat_test$p < 0.001, "***",
  ifelse(stat_test$p < 0.01, "**",
         ifelse(stat_test$p < 0.05, "*", "ns"))
)

# Ensure stat_test is ordered to match Time_Point levels
stat_test <- stat_test[order(stat_test$Time_Point), ]

# 4. Create the plot
Fig_f <- ggplot(Blues_ID_T, aes(x = Time_Point, y =  Flowering, color = Type)) +
  geom_boxplot(position = position_dodge(dodge_width)) +
  stat_summary(fun = mean, geom = "crossbar",linetype="dotted",width=.75,position = position_dodge(dodge_width)) +
  geom_jitter(position = position_jitterdodge(jitter.width = 0.3, dodge.width = dodge_width), alpha = 0.7) +
  labs(y = "Days to flowering (DAS)", x = "Sowing date", tag="(d)", color = "Ecotype") +
  scale_color_manual(values = c("#B22222", "#3B4992FF") ) +
  scale_x_discrete(labels=c("S1", "S2","S3"))+
  scale_y_continuous(limits = c(0, 170), breaks = seq(0,170,20),expand = c(0, 0)) +
  theme_bw() +
  theme(
    axis.title.x = element_blank(),
    axis.text.x = element_text(size = 12, colour = "black"),
    axis.text.y = element_text(size = 12, colour = "black"),
    axis.title.y = element_text(size = 14, face = "bold"),
    plot.tag=element_text(face = "bold"),
    legend.title = element_text(face = "bold",size=14), 
    legend.text=element_text( size=12),
    legend.position = "bottom",
  ) +
  
  # Add horizontal lines and p-values manually
  geom_segment(
    data = data.frame(
      Time_Point = time_points,
      x_start = 1:n_timepoints - dodge_width/2,
      x_end = 1:n_timepoints + dodge_width/2,
      y = y_positions
    ),
    aes(x = x_start, xend = x_end, y = y, yend = y),
    color = "black",
    inherit.aes = FALSE
  ) +
  geom_text(
    data = data.frame(
      Time_Point = time_points,
      x = 1:n_timepoints,
      y = y_positions + 5,  # Slightly above the line
      label = paste("p =", stat_test$p.format, stat_test$p_annot)
    ),
    aes(x = x, y = y, label = label),
    color = "black",
    size = 3.5,
    inherit.aes = FALSE
  )
# Display the plot
print(Fig_f)

###Days to Maturity
# Perform pairwise comparisons
stat_test <- compare_means(
  Maturity ~ Type,
  group.by = "Time_Point",
  data = Blues_ID_T,
  method = "t.test"
)

# Print results 
print(stat_test)

#  Manually calculate positions for lines and p-values

# 1. Define x-axis positions for dodged boxplots
dodge_width <- 0.8
time_points <- levels(Blues_ID_T$Time_Point)
n_timepoints <- length(time_points)

# Calculate x-coordinates for Highland (left) and Coastal (right) within each Time_Point
x_positions <- data.frame(
  Time_Point = rep(time_points, each = 2),
  Type = rep(c( "Coastal", "Highland"), times = n_timepoints),
  x = rep(1:n_timepoints, each = 2) + c(-dodge_width/2, dodge_width/2)
)


# 2. Define y_positions for horizontal lines (above the data)
y_max <- 255  # Based on your y-axis limit
y_start <- y_max - 10
y_step <- 2  # Spacing between lines
y_positions <- seq(y_start, y_start - (n_timepoints - 1) * y_step, by = -y_step)

# 3. Prepare p-value annotations with significance stars
stat_test$p_annot <- ifelse(
  stat_test$p < 0.001, "***",
  ifelse(stat_test$p < 0.01, "**",
         ifelse(stat_test$p < 0.05, "*", "ns"))
)

# Ensure stat_test is ordered to match Time_Point levels
stat_test <- stat_test[order(stat_test$Time_Point), ]

# 4. Create the plot
Fig_m <- ggplot(Blues_ID_T, aes(x = Time_Point, y =  Maturity, color = Type)) +
  geom_boxplot(position = position_dodge(dodge_width)) +
  stat_summary(fun = mean, geom = "crossbar",linetype="dotted",width=.75,position = position_dodge(dodge_width)) +
  geom_jitter(position = position_jitterdodge(jitter.width = 0.3, dodge.width = dodge_width), alpha = 0.7) +
  labs(y = "Days to maturity (DAS)", x = "Sowing date", tag="(e)", color = "Ecotype") +
  scale_color_manual(values = c("#B22222", "#3B4992FF") ) +
  scale_x_discrete(labels=c("S1", "S2","S3"))+
  scale_y_continuous(limits = c(100, 260), breaks = seq(0,260,20),expand = c(0, 0)) +
  theme_bw() +
  theme(
    axis.title.x = element_blank(),
    axis.text.x = element_text(size = 12, colour = "black"),
    axis.text.y = element_text(size = 12, colour = "black"),
    axis.title.y = element_text(size = 14, face = "bold"),
    plot.tag=element_text(face = "bold"),
    legend.title = element_text(face = "bold",size=14), 
    legend.text=element_text( size=12),
    legend.position = "bottom",
  ) +
  
  # Add horizontal lines and p-values manually
  geom_segment(
    data = data.frame(
      Time_Point = time_points,
      x_start = 1:n_timepoints - dodge_width/2,
      x_end = 1:n_timepoints + dodge_width/2,
      y = y_positions
    ),
    aes(x = x_start, xend = x_end, y = y, yend = y),
    color = "black",
    inherit.aes = FALSE
  ) +
  geom_text(
    data = data.frame(
      Time_Point = time_points,
      x = 1:n_timepoints,
      y = y_positions + 5,  # Slightly above the line
      label = paste("p =", stat_test$p.format, stat_test$p_annot)
    ),
    aes(x = x, y = y, label = label),
    color = "black",
    size = 3.5,
    inherit.aes = FALSE
  )
# Display the plot
print(Fig_m)

###Plant height
# Perform pairwise comparisons
stat_test <- compare_means(
  Plant_Height ~ Type,
  group.by = "Time_Point",
  data = Blues_ID_T,
  method = "t.test"
)

# Print results 
print(stat_test)

#  Manually calculate positions for lines and p-values

# 1. Define x-axis positions for dodged boxplots
dodge_width <- 0.8
time_points <- levels(Blues_ID_T$Time_Point)
n_timepoints <- length(time_points)

# Calculate x-coordinates for Highland (left) and Coastal (right) within each Time_Point
x_positions <- data.frame(
  Time_Point = rep(time_points, each = 2),
  Type = rep(c( "Coastal", "Highland"), times = n_timepoints),
  x = rep(1:n_timepoints, each = 2) + c(-dodge_width/2, dodge_width/2)
)

# 2. Define y_positions for horizontal lines (above the data)
y_max <- 250  # Based on your y-axis limit
y_start <- y_max - 20
y_step <- 2  # Spacing between lines
y_positions <- seq(y_start, y_start - (n_timepoints - 1) * y_step, by = -y_step)

# 3. Prepare p-value annotations with significance stars
stat_test$p_annot <- ifelse(
  stat_test$p < 0.001, "***",
  ifelse(stat_test$p < 0.01, "**",
         ifelse(stat_test$p < 0.05, "*", "ns"))
)

# Ensure stat_test is ordered to match Time_Point levels
stat_test <- stat_test[order(stat_test$Time_Point), ]

# 4. Create the plot
Fig_ph <- ggplot(Blues_ID_T, aes(x = Time_Point, y =  Plant_Height, color = Type)) +
  geom_boxplot(position = position_dodge(dodge_width)) +
  stat_summary(fun = mean, geom = "crossbar",linetype="dotted",width=.75,position = position_dodge(dodge_width)) +
  geom_jitter(position = position_jitterdodge(jitter.width = 0.3, dodge.width = dodge_width), alpha = 0.7) +
  labs(y = "Plant height (cm)", x = "Sowing date",tag="(g)", color = "Ecotype") +
  scale_color_manual(values = c("#B22222", "#3B4992FF") ) +
  scale_x_discrete(labels=c("S1", "S2","S3"))+
  scale_y_continuous(limits = c(0, 260), breaks = seq(0,260,20),expand = c(0, 0)) +
  theme_bw() +
  theme(
    axis.title.x = element_text(size = 14, face = "bold"),
    axis.text.x = element_text(size = 12, colour = "black"),
    axis.text.y = element_text(size = 12, colour = "black"),
    axis.title.y = element_text(size = 14,face = "bold"),
    plot.tag=element_text(face = "bold"),
    legend.title = element_text(face = "bold",size=14), 
    legend.text=element_text( size=12),
    legend.position = "bottom",
  ) +
  
  # Add horizontal lines and p-values manually
  geom_segment(
    data = data.frame(
      Time_Point = time_points,
      x_start = 1:n_timepoints - dodge_width/2,
      x_end = 1:n_timepoints + dodge_width/2,
      y = y_positions
    ),
    aes(x = x_start, xend = x_end, y = y, yend = y),
    color = "black",
    inherit.aes = FALSE
  ) +
  geom_text(
    data = data.frame(
      Time_Point = time_points,
      x = 1:n_timepoints,
      y = y_positions + 10,  # Slightly above the line
      label = paste("p =", stat_test$p.format, stat_test$p_annot)
    ),
    aes(x = x, y = y, label = label),
    color = "black",
    size = 3.5,
    inherit.aes = FALSE
  )
# Display the plot
print(Fig_ph)

###Panicle length
# Perform pairwise comparisons
stat_test <- compare_means(
  Panicle_Length ~ Type,
  group.by = "Time_Point",
  data = Blues_ID_T,
  method = "t.test"
)

# Print results 
print(stat_test)

#  Manually calculate positions for lines and p-values

# 1. Define x-axis positions for dodged boxplots
dodge_width <- 0.8
time_points <- levels(Blues_ID_T$Time_Point)
n_timepoints <- length(time_points)

# Calculate x-coordinates for Highland (left) and Coastal (right) within each Time_Point
x_positions <- data.frame(
  Time_Point = rep(time_points, each = 2),
  Type = rep(c( "Coastal", "Highland"), times = n_timepoints),
  x = rep(1:n_timepoints, each = 2) + c(-dodge_width/2, dodge_width/2)
)

# 2. Define y_positions for horizontal lines (above the data)
y_max <- 85  # Based on your y-axis limit
y_start <- y_max - 2
y_step <- 2  # Spacing between lines
y_positions <- seq(y_start, y_start - (n_timepoints - 1) * y_step, by = -y_step)

# 3. Prepare p-value annotations with significance stars
stat_test$p_annot <- ifelse(
  stat_test$p < 0.001, "***",
  ifelse(stat_test$p < 0.01, "**",
         ifelse(stat_test$p < 0.05, "*", "ns"))
)

# Ensure stat_test is ordered to match Time_Point levels
stat_test <- stat_test[order(stat_test$Time_Point), ]

# 4. Create the plot
Fig_pl <- ggplot(Blues_ID_T, aes(x = Time_Point, y =  Panicle_Length, color = Type)) +
  geom_boxplot(position = position_dodge(dodge_width)) +
  stat_summary(fun = mean, geom = "crossbar",linetype="dotted",width=.75,position = position_dodge(dodge_width)) +
  geom_jitter(position = position_jitterdodge(jitter.width = 0.3, dodge.width = dodge_width), alpha = 0.7) +
  labs(y = "Panicle length (cm)", x = "Sowing date", tag="(h)",color = "Ecotype") +
  scale_color_manual(values = c("#B22222", "#3B4992FF") ) +
  scale_x_discrete(labels=c("S1", "S2","S3"))+
  scale_y_continuous(limits = c(0, 100), breaks = seq(0,100,10),expand = c(0, 0)) +
  theme_bw() +
  theme(
    axis.title.x = element_text(size = 14, face = "bold"),
    axis.text.x = element_text(size = 12, colour = "black"),
    axis.text.y = element_text(size = 12, colour = "black"),
    axis.title.y = element_text(size = 14,face = "bold"),
    plot.tag=element_text(face = "bold"),
    legend.title = element_text(face = "bold",size=14), 
    legend.text=element_text( size=12),
    legend.position = "bottom",
  ) +
  
  # Add horizontal lines and p-values manually
  geom_segment(
    data = data.frame(
      Time_Point = time_points,
      x_start = 1:n_timepoints - dodge_width/2,
      x_end = 1:n_timepoints + dodge_width/2,
      y = y_positions
    ),
    aes(x = x_start, xend = x_end, y = y, yend = y),
    color = "black",
    inherit.aes = FALSE
  ) +
  geom_text(
    data = data.frame(
      Time_Point = time_points,
      x = 1:n_timepoints,
      y = y_positions + 2.5,  # Slightly above the line
      label = paste("p =", stat_test$p.format, stat_test$p_annot)
    ),
    aes(x = x, y = y, label = label),
    color = "black",
    size = 3.5,
    inherit.aes = FALSE
  )
# Display the plot
print(Fig_pl)

###Panicle width
# Perform pairwise comparisons
stat_test <- compare_means(
  Panicle_Width ~ Type,
  group.by = "Time_Point",
  data = Blues_ID_T,
  method = "t.test"
)

# Print results 
print(stat_test)

#  Manually calculate positions for lines and p-values

# 1. Define x-axis positions for dodged boxplots
dodge_width <- 0.8
time_points <- levels(Blues_ID_T$Time_Point)
n_timepoints <- length(time_points)

# Calculate x-coordinates for Highland (left) and Coastal (right) within each Time_Point
x_positions <- data.frame(
  Time_Point = rep(time_points, each = 2),
  Type = rep(c( "Coastal", "Highland"), times = n_timepoints),
  x = rep(1:n_timepoints, each = 2) + c(-dodge_width/2, dodge_width/2)
)

# 2. Define y_positions for horizontal lines (above the data)
y_max <- 14  # Based on your y-axis limit
y_start <- y_max - 1
y_step <- 1  # Spacing between lines
y_positions <- seq(y_start, y_start - (n_timepoints - 1) * y_step, by = -y_step)

# 3. Prepare p-value annotations with significance stars
stat_test$p_annot <- ifelse(
  stat_test$p < 0.001, "***",
  ifelse(stat_test$p < 0.01, "**",
         ifelse(stat_test$p < 0.05, "*", "ns"))
)

# Ensure stat_test is ordered to match Time_Point levels
stat_test <- stat_test[order(stat_test$Time_Point), ]

# 4. Create the plot
Fig_pw <- ggplot(Blues_ID_T, aes(x = Time_Point, y =  Panicle_Width, color = Type)) +
  geom_boxplot(position = position_dodge(dodge_width)) +
  stat_summary(fun = mean, geom = "crossbar",linetype="dotted",width=.75,position = position_dodge(dodge_width)) +
  geom_jitter(position = position_jitterdodge(jitter.width = 0.3, dodge.width = dodge_width), alpha = 0.7) +
  labs(y = "Panicle width (cm)", x = "Sowing date",tag="(i)", color = "Ecotype") +
  scale_color_manual(values = c("#B22222", "#3B4992FF") ) +
  scale_x_discrete(labels=c("S1", "S2","S3"))+
  scale_y_continuous(limits = c(0, 15), breaks = seq(0,15,1),expand = c(0, 0)) +
  theme_bw() +
  theme(
    axis.title.x = element_text(size = 14, face = "bold"),
    axis.text.x = element_text(size = 12, colour = "black"),
    axis.text.y = element_text(size = 12, colour = "black"),
    axis.title.y = element_text(size = 14,face = "bold"),
    plot.tag=element_text(face = "bold"),
    legend.title = element_text(face = "bold",size=14), 
    legend.text=element_text( size=12),
    legend.position = "bottom",
  ) +
  
  # Add horizontal lines and p-values manually
  geom_segment(
    data = data.frame(
      Time_Point = time_points,
      x_start = 1:n_timepoints - dodge_width/2,
      x_end = 1:n_timepoints + dodge_width/2,
      y = y_positions
    ),
    aes(x = x_start, xend = x_end, y = y, yend = y),
    color = "black",
    inherit.aes = FALSE
  ) +
  geom_text(
    data = data.frame(
      Time_Point = time_points,
      x = 1:n_timepoints,
      y = y_positions + 0.5,  # Slightly above the line
      label = paste("p =", stat_test$p.format, stat_test$p_annot)
    ),
    aes(x = x, y = y, label = label),
    color = "black",
    size = 3.5,
    inherit.aes = FALSE
  )
# Display the plot
print(Fig_pw)

###Seed yield 
# Perform pairwise comparisons
stat_test <- compare_means(
  Seed_Yield ~ Type,
  group.by = "Time_Point",
  data = Blues_ID_T,
  method = "t.test"
)

# Print results 
print(stat_test)

#  Manually calculate positions for lines and p-values

# 1. Define x-axis positions for dodged boxplots
dodge_width <- 0.8
time_points <- levels(Blues_ID_T$Time_Point)
n_timepoints <- length(time_points)

# Calculate x-coordinates for Highland (left) and Coastal (right) within each Time_Point
x_positions <- data.frame(
  Time_Point = rep(time_points, each = 2),
  Type = rep(c( "Coastal", "Highland"), times = n_timepoints),
  x = rep(1:n_timepoints, each = 2) + c(-dodge_width/2, dodge_width/2)
)

# 2. Define y_positions for horizontal lines (above the data)
y_max <- 250  # Based on your y-axis limit
y_start <- y_max - 15
y_step <- 2  # Spacing between lines
y_positions <- seq(y_start, y_start - (n_timepoints - 1) * y_step, by = -y_step)

# 3. Prepare p-value annotations with significance stars
stat_test$p_annot <- ifelse(
  stat_test$p < 0.001, "***",
  ifelse(stat_test$p < 0.01, "**",
         ifelse(stat_test$p < 0.05, "*", "ns"))
)

# Ensure stat_test is ordered to match Time_Point levels
stat_test <- stat_test[order(stat_test$Time_Point), ]

# 4. Create the plot
Fig_sy <- ggplot(Blues_ID_T, aes(x = Time_Point, y =  Seed_Yield, color = Type)) +
  geom_boxplot(position = position_dodge(dodge_width)) +
  stat_summary(fun = mean, geom = "crossbar",linetype="dotted",width=.75,position = position_dodge(dodge_width)) +
  geom_jitter(position = position_jitterdodge(jitter.width = 0.3, dodge.width = dodge_width), alpha = 0.7) +
  labs(y = "Seed Yield (g)", x = "Sowing date", tag="(f)",color = "Ecotype") +
  scale_color_manual(values = c("#B22222", "#3B4992FF") ) +
  scale_x_discrete(labels=c("S1", "S2","S3"))+
  scale_y_continuous(limits = c(0, 260), breaks = seq(0,260,20),expand = c(0, 0)) +
  theme_bw() +
  theme(
    axis.title.x = element_blank(),
    axis.text.x = element_text(size = 12, colour = "black"),
    axis.text.y = element_text(size = 12, colour = "black"),
    axis.title.y = element_text(size = 14,face = "bold"),
    plot.tag=element_text(face = "bold"),
    legend.title = element_text(face = "bold",size=14), 
    legend.text=element_text( size=12),
    legend.position = "bottom",
  ) +
  
  # Add horizontal lines and p-values manually
  geom_segment(
    data = data.frame(
      Time_Point = time_points,
      x_start = 1:n_timepoints - dodge_width/2,
      x_end = 1:n_timepoints + dodge_width/2,
      y = y_positions
    ),
    aes(x = x_start, xend = x_end, y = y, yend = y),
    color = "black",
    inherit.aes = FALSE
  ) +
  geom_text(
    data = data.frame(
      Time_Point = time_points,
      x = 1:n_timepoints,
      y = y_positions + 10,  # Slightly above the line
      label = paste("p =", stat_test$p.format, stat_test$p_annot)
    ),
    aes(x = x, y = y, label = label),
    color = "black",
    size = 3.5,
    inherit.aes = FALSE
  )
# Display the plot
print(Fig_sy)

Fig_traits<- ggarrange(Fig_gr,Fig_gp,Fig_pe,
                       Fig_f,Fig_m,Fig_sy,
                       Fig_ph,Fig_pl,Fig_pw, 
                       ncol = 3, nrow = 3, 
                       common.legend = TRUE, 
                       legend = "bottom")
Fig_traits#Fig4
ggsave(plot = Fig_traits, width=12, height=12, dpi = 600, filename = "Fig_4.png")

##Figure_S4 Growth stages within ecotypes-----------------------------------------
stages_data<- Blues_ID_T[c(1:3,5,6,7,16,18)]
stages_graph<- stages_data%>%
  mutate(vegetative_phase= Panicle_Emergence-Germination_DAS,
         Reproductive_phase1= Flowering-Panicle_Emergence,
         Reproductive_phase2= Maturity-Flowering)
stages_graph<-stages_graph%>%
  relocate(vegetative_phase, .before= Germination_DAS) %>%
  relocate(Reproductive_phase1, .before= vegetative_phase) %>%
  relocate(Reproductive_phase2, .before= Reproductive_phase1)

stages_graph<-gather(stages_graph, Growth_stage, DAS, Reproductive_phase2:Germination_DAS,
                     factor_key = TRUE)

stack_t1 <- stages_graph %>%
  filter(Time_Point == "T1") 
stack_t2 <- stages_graph %>%
  filter(Time_Point == "T2") 
stack_t3 <- stages_graph %>%
  filter(Time_Point == "T3") 


fig_stack1<-ggplot(stack_t1, aes(fill=Growth_stage, x= Accession, y=DAS))+
  geom_bar(width = 0.9, position="stack",stat="identity")+
  labs(tag= "(a)")+
  scale_y_continuous(limits = c(0,240),
                     breaks = seq(0,240,20))+
  scale_fill_manual(values=wes_palette("Zissou1"), name = "Growth stage", 
                    labels = c("Maturity", "Flowering",
                               "Panicle Emergence","Seedling Emergence"))+
  theme_bw()+facet_grid(~Type, scales="free_x", space = "free")+
  theme(strip.text.x = element_text(size = 16,face="bold" ),
        plot.tag=element_text(face="bold", size= 14),
        axis.text.x = element_text( angle=90, colour="black"),
        axis.text.y = element_text( colour="black"),
        axis.title.x = element_blank(),
        axis.title.y = element_blank(),
        legend.title = element_text(face = "bold",size=14), 
        legend.text=element_text( size=12),
        legend.position = "bottom",
        #plot.margin = unit(c(0,1,1,0),"cm")
  )
fig_stack1


fig_stack2<-ggplot(stack_t2, aes(fill=Growth_stage, x= Accession, y=DAS))+
  geom_bar(width = 0.9, position="stack",stat="identity")+
  labs(tag= "(b)")+
  scale_y_continuous(limits = c(0,240),
                     breaks = seq(0,240,20))+
  scale_fill_manual(values=wes_palette("Zissou1"), name = "Growth stage", 
                    labels = c("Maturity", "Flowering",
                               "Panicle Emergence","Seedling Emergence"))+
  theme_bw()+facet_grid(~Type, scales="free_x", space = "free")+
  theme(strip.text.x = element_blank(),
        plot.tag=element_text(face="bold"),
        axis.text.x = element_text(angle=90, colour="black"),
        axis.text.y = element_text( colour="black"),
        axis.title.x = element_blank(),
        axis.title.y = element_blank(),
        legend.title = element_text(face = "bold",size=14), 
        legend.text=element_text( size=12),
        legend.position = "bottom",
        #plot.margin = unit(c(0,1,1,0),"cm")
  )

fig_stack2

fig_stack3<-ggplot(stack_t3, aes(fill=Growth_stage, x= Accession, y=DAS))+
  geom_bar(width = 0.9, position="stack",stat="identity")+
  labs(tag= "(c)",x="Accessions")+
  scale_y_continuous(limits = c(0,240),
                     breaks = seq(0,240,20))+
  scale_fill_manual(values=wes_palette("Zissou1"), name = "Growth stage", 
                    labels = c("Maturity", "Flowering",
                               "Panicle Emergence","Seedling Emergence"))+
  theme_bw()+facet_grid(~Type, scales="free_x", space = "free")+
  theme(strip.text.x = element_blank(),
        plot.tag=element_text(face="bold"),
        axis.text.x = element_text( angle=90, colour="black"),
        axis.text.y = element_text( colour="black"),
        axis.title.x = element_text( size = 16, face = "bold"),
        axis.title.y = element_blank(),
        legend.title = element_text(face = "bold",size=14), 
        legend.text=element_text( size=12),
        legend.position = "bottom",
        #plot.margin = unit(c(0,1,1,0),"cm")
  )
fig_stack3  

Fig_S4<- ggarrange(fig_stack1,fig_stack2,fig_stack3, nrow=3,common.legend = TRUE,
                  legend="bottom"
)

Fig_S4<-annotate_figure(Fig_3,left = text_grob("DAS",size = 14, face="bold",vjust = 1,rot=90))
Fig_S4 ## Figure S4

ggsave(plot = Fig_S4,width= 15.77, height= 9.50, dpi = 600, filename = "Fig_S4.png")

##Heritability-------------------------------------------------------------------

#Heritability values are under section 4.1

##Figure_S5 Correlations-------------------------------------------------------------
install.packages("gridGraphics")
library("gridGraphics")
Corr_ID<- Blues_ID%>% 
  dplyr::select(1:10)%>%
  rename(MET=Germination_DAS, EP= Germination_Perc, DTP= Panicle_Emergence,
         DTF= Flowering,DTM= Maturity, SY=Seed_Yield, PH=Plant_Height,PL=Panicle_Length,
         PW= Panicle_Width)

Corr_ID_T<- Blues_ID_T%>% 
  rename(MET=Germination_DAS, EP= Germination_Perc, DTP= Panicle_Emergence,
         DTF= Flowering,DTM= Maturity, SY=Seed_Yield, PH=Plant_Height,PL=Panicle_Length,
         PW= Panicle_Width)
Corr_T1<-Corr_ID_T%>%
  filter(Time_Point=="T1")%>%
  dplyr::select(c(1:11))
Corr_T2<-Corr_ID_T%>%
  filter(Time_Point=="T2")%>%
  dplyr::select(c(1:11))
Corr_T3<-Corr_ID_T%>%
  filter(Time_Point=="T3")%>%
  dplyr::select(c(1:11))

png("correlation_plot_1.png", width = 800, height = 600)
chart.Correlation(Corr_T1[, 3:11],method= "pearson", histogram=FALSE, pch=19)
dev.off()

png("correlation_plot_2.png", width = 800, height = 600)
chart.Correlation(Corr_T2[, 3:11],method= "pearson", histogram=FALSE, pch=19)
dev.off()

png("correlation_plot_3.png", width = 800, height = 600)
chart.Correlation(Corr_T3[, 3:11],method= "pearson", histogram=FALSE, pch=19)
dev.off()

png("correlation_plot_4.png", width = 800, height = 600)
chart.Correlation(Corr_ID[, 2:10], method= "pearson",histogram=FALSE, pch=19)
dev.off()

library(gridExtra)
library(png)
img1 <- readPNG("correlation_plot_1.png")
img2 <- readPNG("correlation_plot_2.png")
img3 <- readPNG("correlation_plot_3.png")
img4 <- readPNG("correlation_plot_4.png")

# Now, arrange the plots using ggarrange
correlation<-grid.arrange(rasterGrob(img1), rasterGrob(img2), rasterGrob(img3), rasterGrob(img4),
                          ncol = 2, nrow = 2)
ggsave(correlation, filename= "Figure_S5_correlation.png")

##Figure_5a and Figure_S6 Phenotypic PCA---------------------------------------------------------

library("FactoMineR")
library("factoextra")
str(Blues_ID)
Blues_ID$Accession<-as.factor(as.character(Blues_ID$Accession))
Blues_ID$Accession<-as.factor(as.character(Blues_ID$Accession))
Blues_ID_pca<- Blues_ID%>%
  dplyr::select(15,2:10,17)%>%
  rename(c(Mean_emergence_time= Germination_DAS,
           Emergence_percentage= Germination_Perc,
  ))
rownames(Blues_ID_pca) <- Blues_ID_pca$Accession
Blues_ID_pca$Type<-as.factor(as.character(Blues_ID_pca$Type))
Blues_ID_pca


res.pca <- PCA(Blues_ID_pca[,2:10], graph = FALSE)

write.csv(res.pca$var$coord, "phenotypic_pca.csv", na="NA")# Table 9

#Variances of the principal components

eigenvalues <- res.pca$eig
head(eigenvalues[, 1:2])

#Make a scree plot using base graphic

barplot(eigenvalues[, 2], names.arg=1:nrow(eigenvalues), 
        main = "Variances",
        xlab = "Principal Components",
        ylab = "Percentage of variances",
        col ="steelblue")

# Add connected line segments to the plot

lines(x = 1:nrow(eigenvalues), eigenvalues[, 2], 
      type="b", pch=19, col = "red")

#Make the scree plot using the package factoextra

fviz_screeplot(res.pca, ncp=10)
# Assign rownames
rownames(Blues_ID_pca) <- Blues_ID_pca$Accession

# Select numeric columns for PCA
numeric_data <- Blues_ID_pca[, sapply(Blues_ID_pca, is.numeric)]

# PCA
res.pca <- prcomp(numeric_data, scale. = TRUE)
#Figure 9
fviz_pca_biplot(res.pca,habillage = Blues_ID_pca$Type,
                col.var = "black", repel= TRUE, 
                label = c("ind","var"), lab.cex=2.5) +
  labs(x="PC 1 (34.3%)",y="PC 2 (19.6%)")+
  scale_color_manual(values = c("#B22222","#3B4992FF"))+
  theme_bw()+
  theme(axis.text.x = element_text( size=12),
        axis.text.y = element_text( size=12),
        axis.title.x = element_text( size = 16, face = "bold"),
        axis.title.y = element_text( size = 16, face = "bold"),
        legend.title = element_text( face="bold",size = 14),
        legend.text = element_text(size=12))


dimdesc(res.pca, axes=c(1,2))

#Figure 10
fviz_pca_biplot(res.pca,habillage = Blues_ID_pca$Type,axes = c(3,4),
                col.var = "black", repel= TRUE, 
                label = c("ind","var"), lab.cex=2) +
  labs(x="PC 3 (18%)",y="PC 4 (9.1%)")+
  scale_color_manual(values = c("#B22222","#3B4992FF"))+
  theme_bw()+
  theme(axis.text.x = element_text( size=12),
        axis.text.y = element_text( size=12),
        axis.title.x = element_text( size = 16, face = "bold"),
        axis.title.y = element_text( size = 16, face = "bold"),
        legend.title = element_text( face="bold",size = 14),
        legend.text = element_text(size=12))


# Load libraries

library(FactoMineR)
library(factoextra)
library(dplyr)
library(ggplot2)


# 1️⃣ Prepare the dataset


# Convert Accession and Type to factors
Blues_ID <- Blues_ID %>%
  mutate(
    Accession = as.factor(as.character(Accession)),
    Type = as.factor(as.character(Type))
  )

# Select columns for PCA
# Example: Accession, Type, and numeric traits in columns 2:10
Blues_ID_pca <- Blues_ID %>%
  dplyr::select(Accession, Type, 2:10)
Blues_ID_pca<- Blues_ID%>%
  dplyr::select(15,2:10,17)%>%
  rename(c(Mean_emergence_time= Germination_DAS,
           Emergence_percentage= Germination_Perc,
  ))

# Keep only numeric columns for PCA and convert to data.frame
numeric_data <- Blues_ID_pca %>%
  dplyr::select(where(is.numeric)) %>%
  as.data.frame()


# 2️⃣ Handle missing/infinite values

complete_rows <- complete.cases(numeric_data)
numeric_data <- numeric_data[complete_rows, ]

# Assign rownames matching numeric_data
rownames(numeric_data) <- Blues_ID_pca$Accession[complete_rows]


# 3️⃣ Perform PCA

res.pca <- PCA(numeric_data, graph = FALSE)


# 4️⃣ Scree plot

eigenvalues <- res.pca$eig

# Base R scree plot
barplot(eigenvalues[,2], names.arg = 1:nrow(eigenvalues),
        main = "Variance Explained by Principal Components",
        xlab = "Principal Components",
        ylab = "Percentage of variance",
        col = "steelblue")
lines(x = 1:nrow(eigenvalues), eigenvalues[,2], type="b", pch=19, col="red")

# Scree plot with factoextra
fviz_screeplot(res.pca, ncp = 10)


# 5️⃣ PCA biplots


# PC1 vs PC2
p1<-fviz_pca_biplot(res.pca,
                habillage = Blues_ID_pca$Type[complete_rows],  # color by Type
                col.var = "black",
                repel = TRUE,
                label = c("ind","var"), 
                lab.cex = 2.5) +
  labs(x = paste0("PC 1 (", round(eigenvalues[1,2],1), "%)"),
       y = paste0("PC 2 (", round(eigenvalues[2,2],1), "%)")) +
  scale_color_manual(values = c("#B22222","#3B4992FF")) +
  theme_bw() +
  theme(axis.text = element_text(size=12),
        axis.title = element_text(size=16, face="bold"),
        legend.title = element_text(size=14, face="bold"),
        legend.text = element_text(size=12))
# Save as PNG
ggsave("PCA_PC1_PC2.png", plot = p1, width = 12, height = 9, dpi = 300)

# PC3 vs PC4
p2<-fviz_pca_biplot(res.pca,
                habillage = Blues_ID_pca$Type[complete_rows],
                axes = c(3,4),
                col.var = "black",
                repel = TRUE,
                label = c("ind","var"),
                lab.cex = 2) +
  labs(x = paste0("PC 3 (", round(eigenvalues[3,2],1), "%)"),
       y = paste0("PC 4 (", round(eigenvalues[4,2],1), "%)")) +
  scale_color_manual(values = c("#B22222","#3B4992FF")) +
  theme_bw() +
  theme(axis.text = element_text(size=12),
        axis.title = element_text(size=16, face="bold"),
        legend.title = element_text(size=14, face="bold"),
        legend.text = element_text(size=12))

ggsave("PCA_PC3_PC4.png", plot = p2, width = 12, height = 9, dpi = 300)


# 6️⃣ Dimensional description

dimdesc(res.pca, axes = c(1,2))

# Round variable coordinates to 2 decimal places
var_coord_rounded <- round(res.pca$var$coord, 3)

# Save to CSV
write.csv(var_coord_rounded, "phenotypic_pca.csv", na = "NA", row.names = TRUE)
##Figure_5b Multi-trait stability index----------------------------------------------

data_PerPlot<-read.csv("data_PerPlot.csv",header=TRUE)
data_PerPlot <- data_PerPlot[,-1]

mtsi_inputdata <- data_PerPlot %>% filter(!ID %in% c("A13", "A22","A25","A30","A37"))#accessions with incomplete data filtered out
write.csv(mtsi_inputdata, "mtsi_inputdata.csv")

mtsi_inputdata<- read.csv("mtsi_inputdata.csv",header=TRUE)

waasby_model<- 
  waasb(mtsi_inputdata,
        env = Time_Point,
        gen = Accession,
        rep = Rep,
        resp = c( Germination_Perc,Plant_Height,Panicle_Width, 
                  Germination_DAS,Maturity,Flowering,Seed_Yield),
        wresp = 50,
        mresp=c("h,l,h,l,l,l,h"))

mtsi_index<- mtsi(waasby_model,index="waasby")
plot(mtsi_index)

plot(
  mtsi_index,
  SI = 15,
  type = "index",
  position = "fill",
  genotypes = "all",
  title = TRUE,
  radar = TRUE,
  arrange.label = TRUE,
  x.lab = NULL,
  y.lab = NULL,
  size.point = 2.5,
  size.line = 0.6,
  size.text =1.5,
  width.bar = 0.75,
  n.dodge = 1,
  check.overlap = TRUE,
  invert = TRUE,
  col.sel = "red",
  col.nonsel = "black",
  legend.position = "bottom",
)


mtsi_values<-mtsi_index$MTSI
mtsi_values<-mtsi_values%>%
  rename(Accession=Genotype)
mtsi<-merge(mtsi_values, origin_data,by="Accession", all.x=TRUE)
mtsi$MTSI<-round(mtsi$MTSI, digits = 3)
write.csv(mtsi, "mtsi_values.csv", na="NA")
mtsi<-read.csv("mtsi_values.csv",header=TRUE)# Supplementary Table 2

#LABORATORY EXPERIMENT---------------------------------------------------------

data_lab<-read_csv("Lab_germination.csv")# Laboratory germination data loading
Healthy_seedlings <- read_csv("Healthy_Seedlings.csv")# Healthy seedlings dataset uploading
origin_data<-read.csv("acc.csv",header = TRUE)#merging with the accession data
origin_data$ID <- gsub("H21_", "", origin_data$ID)
#1.Calculating germination percentage --------------------------------
##Germination_Perc
perc_lab <- data_lab %>%
  dplyr::group_by(ID,Rep,Treatment) %>%
  dplyr::summarise(Germination_Perc = sum(!is.na(Germination_DAS))/n()*100)

##Healthy_seedlings_Perc
Healthy_Seedlings<-Healthy_seedlings %>%
  dplyr::mutate(Healthy_seedlings_Perc=Healthy_seedlings/10*100 )

write.csv(Healthy_Seedlings, "Healthy_Seedlings.csv")

# 2.Creating average data----------------------------------------------

avg_data_lab <- data_lab %>%
  group_by(ID, Rep, Treatment) %>%
  summarise_at(vars(Germination_DAS),
               .funs = mean,
               na.rm = TRUE)

# 3.Merging datasets------------------------------------------------------

avg_labdata <- merge(avg_data_lab, perc_lab, by = c("ID", "Rep", "Treatment"))
avg_labdata$Germination_Perc <- 
  ifelse(avg_labdata$Germination_Perc == 0 , NA, avg_labdata$Germination_Perc)

avg_labdata <- merge(avg_labdata,origin_data, by = c("ID"))

avg_labdata<-merge(avg_labdata,Healthy_Seedlings,by=c("ID","Rep","Treatment"))

avg_labdata<-avg_labdata%>%relocate(Healthy_seedlings,Healthy_seedlings_Perc, .after=Germination_Perc)
write.csv (avg_labdata, "avg_labdata.csv")# Final average data

avg_labdata["Healthy_seedlings_Perc"][avg_labdata["Healthy_seedlings_Perc"] == 0] <- NA
avg_labdata$Germination_DAS[is.nan(avg_labdata$Germination_DAS)]<-NA

# 4.Rounding and factoring-----------------------------------------------------------------

avg_labdata$Germination_DAS<-round(avg_labdata$Germination_DAS,digits=1)

# 5.Factoring ------------------------------------------------------
data_lab$ID <- as.factor(as.character(data_lab$ID))
data_lab$Treatment <- as.factor(as.character(data_lab$Treatment))
data_lab$Rep <- as.factor(as.character(data_lab$Rep))

avg_labdata$ID <- as.factor(as.character(avg_labdata$ID))
avg_labdata$Treatment <- as.factor(as.character(avg_labdata$Treatment))
avg_labdata$Rep <- as.factor(as.character(avg_labdata$Rep))
str(avg_labdata)
hist(avg_labdata$Germination_DAS)

# 5.BLUEs Calculation---------------------------------------------------------------------------------
traits <- names(avg_labdata[c(4,5,7)])

an <- ID <- ID_T <- ID_R_T <-list(traits)


for (i in traits)
{
  mod_blue <- asreml(as.formula(paste(i,"~ Rep + Treatment+ ID+ Treatment*ID")),
                     random = ~Rep:Treatment,
                     data = avg_labdata)  
  
  pred_blues_ID <- predict(mod_blue, classify = "ID")
  pred_blues_IDxT <- predict(mod_blue, classify = "ID:Treatment")
  pred_blues_IDxRxT <- predict(mod_blue, classify = "ID:Rep:Treatment")
  
  blues_ID <- pred_blues_ID$pvals
  blues_ID <- blues_ID %>%
    dplyr::select(c(1:2))
  
  blues_IDxT <- pred_blues_IDxT$pvals
  blues_IDxT <- blues_IDxT %>%
    dplyr::select(c(1:3))
  
  blues_IDxRxT <- pred_blues_IDxRxT$pvals
  blues_IDxRxT <- blues_IDxRxT %>%
    dplyr::select(c(1:4))
  
  #Variance <- as.data.frame(summary(mod_blue)$varcomp)
  anova <- as.data.frame(wald(mod_blue))
  
  an[[paste(i)]] <- anova
  ID[[paste(i)]] <- blues_ID
  ID_T[[paste(i)]] <- blues_IDxT
  ID_R_T[[paste(i)]] <- blues_IDxRxT
  
}

anv <- do.call(rbind.data.frame, an)
anv <- anv[-1,]
anv$Factor <- rownames(anv)
anv <- anv %>%
  remove_rownames()%>%
  relocate(Factor, .before = Df) %>%
  separate(Factor, c("Trait", "Factor"), sep = "([.])", extra = "merge")

ID <- ID[-1]

Blues_lab_ID <- cbind(ID$Germination_DAS, ID$Germination_Perc$predicted.value,ID$Healthy_seedlings_Perc$predicted.value)
names(Blues_lab_ID)[2:4] <- traits

Blues_lab_ID <- merge(Blues_lab_ID, origin_data, by = "ID")

ID_T <- ID_T[-1]

Blues_lab_ID_T <- cbind(ID_T$Germination_DAS,  ID_T$Germination_Perc$predicted.value,ID_T$Healthy_seedlings_Perc$predicted.value)

names(Blues_lab_ID_T)[3:5] <- traits

Blues_lab_ID_T <- merge(Blues_lab_ID_T, origin_data, by = "ID")

ID_R_T <- ID_R_T[-1]

Blues_lab_ID_R_T <- cbind(ID_R_T$Germination_DAS,ID_R_T$Germination_Perc$predicted.value,ID_R_T$Healthy_seedlings_Perc$predicted.value)

names(Blues_lab_ID_R_T)[4:6] <- traits

Blues_lab_ID_R_T <- merge(Blues_lab_ID_R_T, origin_data, by = "ID")

Blues_lab_ID_R_T$Germination_Perc <- ifelse(Blues_lab_ID_R_T$Germination_Perc <= 0, NA, Blues_lab_ID_R_T$Germination_Perc)

#Output files...............................................................

write.csv(Blues_lab_ID_R_T, "Blues_lab_ID_R_T.csv",na="NA")
write.csv(Blues_lab_ID_T, "Blues_lab_ID_T.csv",na="NA")
write.csv(Blues_lab_ID, "Blues_lab_ID.csv",na="NA")

###5.1 Extracting Wald test results (Table_S5)-----------------------------------------------
str(anv)
anv$`Pr(Chisq)`<-as.numeric(anv$`Pr(Chisq)`)
anv$`Pr(Chisq)` <- round(anv$`Pr(Chisq)`, digits = 3)

stars <- ifelse(anv$`Pr(Chisq)` < .0001, "****",
                ifelse(anv$`Pr(Chisq)` < .001, "***",
                       ifelse(anv$`Pr(Chisq)` < .01, "**",
                              ifelse(anv$`Pr(Chisq)` < .05, "*", ""))))


test_anv <- anv
test_anv$`Pr(Chisq)` <- paste(test_anv$`Pr(Chisq)`, stars, "")
write.csv(test_anv, "Wald_results.csv", na="NA" ) # Table 4

#IMAGE ANALYSIS-------------------------------------------------------

# 1. Data reading and editing------------------------------------------------------
data <- read.table("image_anaylsis.txt", header=T)
head(data)

##count the differences in seedlings between days
df <- data %>% 
  #filter(day <= 3)  %>%  # max until day 3
  filter(count_seedlings <= 10)  %>%  # max count_seedling of 10
  arrange(accession, condition, rep, day) %>% 
  group_by(accession, condition, rep) %>% 
  mutate(seedling_increase = c(first(count_seedlings), diff(count_seedlings))) # calculate differences between days

## prepare new data frame with plant name
lab_germ_data <- data.frame(Genotype=as.character(),
                            Treatment=as.character(),
                            Plant=as.character(),
                            DAS=as.numeric(),
                            Rep=as.character())

## loop over the data frame, create name for "plants" and put it in a new data frame
for (i in 1:nrow(df)) {
  my_row <- df[i, ]
  
  if ( my_row$seedling_increase > 0) { # if the seedling increase is less than 0 we ignore that line (no new "plants" to be added)
    
    start_seed <- my_row$count_seedlings - my_row$seedling_increase # the difference between count_seedling and seedling_increase
    seeds_name <- as.vector(paste0("P", seq(start_seed+1, my_row$count_seedlings, by=1))) # create "plant" name
    n <- length(seeds_name)
    
    lab_germ_data <- rbind(lab_germ_data, 
                           data.frame(Genotype=rep(my_row$accession, n),
                                      Treatment=rep(my_row$condition, n),
                                      Plant=seeds_name,
                                      DAS=rep(my_row$day, n),
                                      Rep=rep(my_row$rep, n)))
  }
}

##removing the duplicated rows

image_analysis_data <- lab_germ_data %>%
  group_by(Genotype, Treatment, Rep) %>%
  filter(!duplicated(Plant))

## Data editing
david_data <- read.csv("Lab_germ.csv", header = TRUE)

david_data <- david_data[,-1]

str(image_analysis_data)

image_analysis_data <- as.data.frame(image_analysis_data)

image_analysis_data$DAS <- as.numeric(as.character(image_analysis_data$DAS))

image_analysis_data[image_analysis_data == "cold"] <- "K"

david_data[david_data == "R1"] <- "1"

david_data[david_data == "R2"] <- "2"

david_data[david_data == "R3"] <- "3"

david_data$Rep <- as.integer(as.character(david_data$Rep))

lab_image_data <- merge(david_data, image_analysis_data, by = c("Genotype", "Rep", "Treatment", "Plant"), 
                        suffix= c("_lab","_image"),
                        all.x = TRUE)

lab_image_data<- lab_image_data[,-5]
str(lab_image_data)

lab_image_data<-lab_image_data%>% rename("DAS"="DAS_image")
lab_image_data<-lab_image_data%>% rename("Germination_DAS"="DAS")
lab_image_data<-lab_image_data%>%rename("ID"="Genotype")

origin_data<-read.csv("acc.csv",header = TRUE)
origin_data$ID <- gsub("H21_", "", origin_data$ID)
lab_image_data <- merge(lab_image_data,origin_data, by = c("ID"))
write.csv(lab_image_data,"lab_image_data.csv")

#2.Calculating germination percentage --------------------------------
##Germination_Perc
perc_lab <- lab_image_data %>%
  dplyr::group_by(ID,Rep,Treatment) %>%
  dplyr::summarise(Germination_Perc = sum(!is.na(Germination_DAS))/n()*100)

# 3.Creating average data----------------------------------------------

avg_data_image <- lab_image_data %>%
  group_by(ID, Rep, Treatment) %>%
  summarise_at(vars(Germination_DAS),
               .funs = mean,
               na.rm = TRUE)

# 4.Merging datasets------------------------------------------------------

avg_imagedata <- merge(avg_data_image, perc_lab, by = c("ID", "Rep", "Treatment"))
avg_imagedata$Germination_Perc <- 
  ifelse(avg_imagedata$Germination_Perc == 0 , NA, avg_imagedata$Germination_Perc)

avg_imagedata <- merge(avg_imagedata,origin_data, by = c("ID"))

##4.1 merging mumax dataset (growth rate) with average dataset----------------------------------------

# read data
data_K <- read.table("growth_rates_fit_cold_days1-15.txt",sep=",",header=T)
data_RT <- read.table("growth_rates_fit_RT_days1-3.txt", sep=",",header=T)

data_K$Treatment <- "K"
data_RT$Treatment <- "RT"
image_mumax_data <- rbind(data_K, data_RT)

image_mumax_data<-image_mumax_data%>%rename("ID"="Accession")
image_mumax_data<-image_mumax_data%>%rename("Rep"="Replicate")

avg_imagedata<-merge(avg_imagedata, image_mumax_data, by = c("ID", "Rep", "Treatment"), 
                     all.x = TRUE)
avg_imagedata<-avg_imagedata[,-12]
avg_imagedata<-avg_imagedata[,-13]
avg_imagedata<-avg_imagedata%>%relocate(mumax, .after=Germination_Perc)

write.csv(avg_imagedata, "avg_imagedata.csv")

# 5.Rounding-----------------------------------------------------------------

avg_imagedata$Germination_DAS<-round(avg_imagedata$Germination_DAS,digits=1)

# 6.Factoring ------------------------------------------------------
lab_image_data$ID <- as.factor(as.character(lab_image_data$ID))
lab_image_data$Treatment <- as.factor(as.character(lab_image_data$Treatment))
lab_image_data$Rep <- as.factor(as.character(lab_image_data$Rep))

avg_imagedata$ID <- as.factor(as.character(avg_imagedata$ID))
avg_imagedata$Treatment <- as.factor(as.character(avg_imagedata$Treatment))

avg_imagedata$Rep <- as.factor(as.character(avg_imagedata$Rep))
str(avg_imagedata)
hist(avg_imagedata$Germination_DAS)

# 7.BLUEs Calculation---------------------------------------------------------------------------------
traits <- names(avg_imagedata[c(4,5,6)])

an <- ID <- ID_T <- ID_R_T <-list(traits)


for (i in traits)
{
  mod_blue <- asreml(as.formula(paste(i,"~ Rep + ID + Treatment + ID*Treatment")),
                     random = ~Rep:ID,
                     data = avg_imagedata)  
  
  pred_blues_ID <- predict(mod_blue, classify = "ID")
  pred_blues_IDxT <- predict(mod_blue, classify = "ID:Treatment")
  pred_blues_IDxRxT <- predict(mod_blue, classify = "ID:Rep:Treatment")
  
  blues_ID <- pred_blues_ID$pvals
  blues_ID <- blues_ID %>%
    dplyr::select(c(1:2))
  
  blues_IDxT <- pred_blues_IDxT$pvals
  blues_IDxT <- blues_IDxT %>%
    dplyr::select(c(1:3))
  
  blues_IDxRxT <- pred_blues_IDxRxT$pvals
  blues_IDxRxT <- blues_IDxRxT %>%
    dplyr::select(c(1:4))
  
  #Variance <- as.data.frame(summary(mod_blue)$varcomp)
  anova <- as.data.frame(wald(mod_blue))
  
  an[[paste(i)]] <- anova
  ID[[paste(i)]] <- blues_ID
  ID_T[[paste(i)]] <- blues_IDxT
  ID_R_T[[paste(i)]] <- blues_IDxRxT
  
}

anv <- do.call(rbind.data.frame, an)
anv <- anv[-1,]
anv$Factor <- rownames(anv)
anv <- anv %>%
  remove_rownames()%>%
  relocate(Factor, .before = Df) %>%
  separate(Factor, c("Trait", "Factor"), sep = "([.])", extra = "merge")

ID <- ID[-1]

Blues_image_ID <- cbind(ID$Germination_DAS,ID$Germination_Perc$predicted.value,ID$mumax$predicted.value)
names(Blues_image_ID)[2:4] <- traits

Blues_image_ID <- merge(Blues_image_ID, origin_data, by = "ID")

ID_T <- ID_T[-1]

Blues_image_ID_T <- cbind(ID_T$Germination_DAS, ID_T$Germination_Perc$predicted.value,ID_T$mumax$predicted.value)

names(Blues_image_ID_T)[3:5] <- traits

Blues_image_ID_T <- merge(Blues_image_ID_T, origin_data, by = "ID")

ID_R_T <- ID_R_T[-1]

Blues_image_ID_R_T <- cbind(ID_R_T$Germination_DAS,ID_R_T$Germination_Perc$predicted.value,ID_R_T$mumax$predicted.value)

names(Blues_image_ID_R_T)[4:6] <- traits

Blues_image_ID_R_T <- merge(Blues_image_ID_R_T, origin_data, by = "ID")

Blues_image_ID_R_T$Germination_Perc <- ifelse(Blues_image_ID_R_T$Germination_Perc <= 0, NA, Blues_image_ID_R_T$Germination_Perc)

###7.1 Extracting Wald test results (Table_S5)-----------------------------------------------
str(anv)
anv$`Pr(Chisq)`<-as.numeric(anv$`Pr(Chisq)`)
anv$`Pr(Chisq)` <- round(anv$`Pr(Chisq)`, digits = 3)

stars <- ifelse(anv$`Pr(Chisq)` < .0001, "****",
                ifelse(anv$`Pr(Chisq)` < .001, "***",
                       ifelse(anv$`Pr(Chisq)` < .01, "**",
                              ifelse(anv$`Pr(Chisq)` < .05, "*", ""))))


test_anv <- anv
test_anv$`Pr(Chisq)` <- paste(test_anv$`Pr(Chisq)`, stars, "")
write.csv(test_anv, "Waldimage_results.csv", na="NA" ) 

#COMPARISONS BETWEEN LABORATORY MANUAL DATA AND IMAGE ANALYSIS DATA------------------------------
#1. Test comparisons between ecotypes under two treatments (Table 3)--------------------------------------------------
#Blues_lab_ID_T <- read_csv("Blues_lab_ID_T.csv")  
traits <- c("Germination_DAS", "Germination_Perc", "Healthy_seedlings_Perc")  # Replace with actual trait names
t.test_lab_type <- map(traits, function(trait) {
  compare_means(
    reformulate("Type", trait),  # Dynamically create formula
    group.by = "Treatment",
    data = Blues_lab_ID_T,
    method = "t.test"
  ) %>%
    mutate(Trait = trait)  # Add trait name to results
})

# Combine all results into a single data frame
t.test_lab_type <- bind_rows(t.test_lab_type)
write.csv(t.test_lab_type,"lab_ecotype_t.test.csv")


#Blues_image_ID_T <- read_csv("Blues_image_ID_T.csv") read the data if required

traits <- c("Germination_DAS", "Germination_Perc", "mumax")  # Replace with actual trait names
t.test_image_type <- map(traits, function(trait) {
  compare_means(
    reformulate("Type", trait),  # Dynamically create formula
    group.by = "Treatment",
    data = Blues_image_ID_T,
    method = "t.test"
  ) %>%
    mutate(Trait = trait)  # Add trait name to results
})

# Combine all results into a single data frame
t.test_image_type <- bind_rows(t.test_image_type)
write.csv(t.test_image_type,"image_ecotype_t.test.csv")
print(t.test_image_type)

#Figure_6 Differences of estimated means (BLUEs) in the laboratory assay of 60 quinoa accessions under the treatments cold and room temperature treatments---------------------------------------------------------------------
##Manuallaboratory ------------------------------------------------
Blues_lab_ID_T$Treatment <- as.character(Blues_lab_ID_T$Treatment)
Blues_lab_ID_T$Treatment[Blues_lab_ID_T$Treatment == "K"] <- "Cold"
Blues_lab_ID_T$Treatment[Blues_lab_ID_T$Treatment == "RT"] <- "Room temperature"
my_comparisons = list( c("Cold", "Room temperature") )
Fig_GR_manual <- ggplot(Blues_lab_ID_T,aes(x=Treatment, y = Germination_DAS,color=Treatment))+
  geom_boxplot(position=position_dodge(0.9))+
  stat_summary(fun=mean, geom="crossbar",linetype="dotted",width=.75,position=position_dodge(0.9))+
  geom_jitter(position = position_jitterdodge(jitter.width = 0.3), alpha = 0.7)+
  labs(y = "Mean germination time (DAS)", x = "Treatment", tag="(a)", color="Treatment")+
  stat_compare_means(comparisons = my_comparisons, label="p.signif",method="t.test", label.y = c(16, 5), na=TRUE)+
  scale_color_manual(values = c("deepskyblue", "dodgerblue4"),
                     labels=c("Image"="Image analysis"))+
  scale_y_continuous(limits = c(0,20),
                     breaks = seq(0,20,2), expand = c(0,0))+
  theme_bw()+
  theme(axis.title = element_text(size = 12, face = "bold"),
        axis.text = element_text(size=12, colour="black"),
        legend.position = "none",
        plot.tag = element_text(face="bold"),
        legend.title = element_text(face = "bold",size=14), 
        legend.text=element_text(size=12))

Fig_GR_manual#Fig 6a
ggsave(plot = Fig_GR_manual,  dpi = 600, filename = "Fig_GR_manual.png")


Fig_GP_manual <- ggplot(Blues_lab_ID_T,aes(x=Treatment, y = Germination_Perc,color=Treatment))+
  geom_boxplot(position=position_dodge(0.9))+
  stat_summary(fun=mean, geom="crossbar",linetype="dotted",width=.75,position=position_dodge(0.9))+
  geom_jitter(position = position_jitterdodge(jitter.width = 0.3), alpha = 0.7)+
  labs(y = "Germination percentage (%)", x = "Treatment", tag="(b)", color="Treatment")+
  stat_compare_means(comparisons = my_comparisons, label="p.signif",method="t.test", label.y = c(105), na=TRUE)+
  scale_color_manual(values = c("deepskyblue", "dodgerblue4"),
                     labels=c("Image"="Image analysis"))+
  scale_y_continuous(limits = c(0,120),
                     breaks = seq(0,120,10), expand = c(0,0))+
  theme_bw()+
  theme(axis.title = element_text(size = 12, face = "bold"),
        axis.text = element_text(size=12, colour="black"),
        legend.position = "none",
        plot.tag = element_text(face="bold"),
        legend.title = element_text(face = "bold",size=14), 
        legend.text=element_text(size=12))

Fig_GP_manual#Fig 6b
ggsave(plot = Fig_GP_manual,  dpi = 600, filename = "Fig_GP_manual.png")


Fig_HGP_manual <- ggplot(Blues_lab_ID_T,aes(x=Treatment, y = Healthy_seedlings_Perc,color=Treatment))+
  geom_boxplot(position=position_dodge(0.9))+
  stat_summary(fun=mean, geom="crossbar",linetype="dotted",width=.75,position=position_dodge(0.9))+
  geom_jitter(position = position_jitterdodge(jitter.width = 0.3), alpha = 0.7)+
  labs(y = "Viable germination percentage (%)", x = "Treatment", tag="(c)", color="Treatment")+
  stat_compare_means(comparisons = my_comparisons, label="p.signif",method="t.test", label.y = c(105), na=TRUE)+
  scale_color_manual(values = c("deepskyblue", "dodgerblue4"),
                     labels=c("Image"="Image analysis"))+
  scale_y_continuous(limits = c(0,120),
                     breaks = seq(0,120,10), expand = c(0,0))+
  theme_bw()+
  theme(axis.title = element_text(size = 12, face = "bold"),
        axis.text = element_text(size=12, colour="black"),
        legend.position = "none",
        plot.tag = element_text(face="bold"),
        legend.title = element_text(face = "bold",size=14), 
        legend.text=element_text( size=12))

Fig_HGP_manual#Fig 6c
ggsave(plot = Fig_GP_manual,  dpi = 600, filename = "Fig_HGP_manual.png")
plot_manual<- ggarrange(Fig_GR_manual, Fig_GP_manual, Fig_HGP_manual,ncol =3)
plot_manual

side_title<-textGrob("Manual scoring",gp=gpar(fontsize=18, fontface="bold"),rot=90, just="center")
plot_manual<-grid.arrange(side_title,plot_manual,ncol=2, widths=c(0.5,10))


top_title<-textGrob("Manual scoring",gp=gpar(fontsize=18, fontface="bold"))

plot_manual+annotation_custom(top_title,xmin=-Inf,xmax = Inf,ymin=-Inf,ymax = Inf)+
  theme(plot.margin = margin(t=10))
plot_manual_top
##Image----------------------------------------------------------------------------
#Blues_image_ID_T<-read_csv("Blues_image_ID_T.csv")
Blues_image_ID_T$Treatment <- as.character(Blues_image_ID_T$Treatment)
Blues_image_ID_T$Treatment[Blues_image_ID_T$Treatment == "K"] <- "Cold"
Blues_image_ID_T$Treatment[Blues_image_ID_T$Treatment == "RT"] <- "Room temperature"

my_comparisons = list( c("Cold", "Room temperature") )
Fig_GR_image <- ggplot(Blues_image_ID_T,aes(x=Treatment, y = Germination_DAS,color=Treatment))+
  geom_boxplot(position=position_dodge(0.9))+
  stat_summary(fun=mean, geom="crossbar",linetype="dotted",width=.75,position=position_dodge(0.9))+
  geom_jitter(position = position_jitterdodge(jitter.width = 0.3), alpha = 0.7)+
  labs(y = "Mean germination time (DAS)", x = "Treatment", tag="(d)", color="Treatment")+
  stat_compare_means(comparisons = my_comparisons, label="p.signif",method="t.test", label.y = c(16, 5), na=TRUE)+
  scale_color_manual(values = c("deepskyblue", "dodgerblue4"),
                     labels=c("Image"="Image analysis"))+
  scale_y_continuous(limits = c(0,20),
                     breaks = seq(0,20,2), expand = c(0,0))+
  theme_bw()+
  theme(axis.title = element_text(size = 12, face = "bold"),
        axis.text = element_text(size=12, colour="black"),
        legend.position = "none",
        plot.tag = element_text(face="bold"),
        legend.title = element_text(face = "bold",size=14), 
        legend.text=element_text( size=12))

Fig_GR_image#Fig 6d
ggsave(plot = Fig_GR_image,  dpi = 600, filename = "Fig_GR_image.png")


Fig_GP_image <- ggplot(Blues_image_ID_T,aes(x=Treatment, y = Germination_Perc,color=Treatment))+
  geom_boxplot(position=position_dodge(0.9))+
  stat_summary(fun=mean, geom="crossbar",linetype="dotted",width=.75,position=position_dodge(0.9))+
  geom_jitter(position = position_jitterdodge(jitter.width = 0.3), alpha = 0.7)+
  labs(y = "Germination percentage (%)", x = "Treatment", tag="(e)", color="Treatment")+
  stat_compare_means(comparisons = my_comparisons, label="p.signif",method="t.test", label.y = c(105), na=TRUE)+
  scale_color_manual(values = c("deepskyblue", "dodgerblue4"),
                     labels=c("Image"="Image analysis"))+
  scale_y_continuous(limits = c(0,120),
                     breaks = seq(0,120,10), expand = c(0,0))+
  theme_bw()+
  theme(axis.title = element_text(size = 12, face = "bold"),
        axis.text = element_text(size=12, colour="black"),
        legend.position = "none",
        plot.tag = element_text(face="bold"),
        legend.title = element_text(face = "bold",size=14), 
        legend.text=element_text( size=12))

Fig_GP_image#Fig 6e
ggsave(plot = Fig_GP_image,  dpi = 600, filename = "Fig_GP_image.png")


Fig_growth_image <- ggplot(Blues_image_ID_T,aes(x=Treatment, y = mumax,color=Treatment))+
  geom_boxplot(position=position_dodge(0.9))+
  stat_summary(fun=mean, geom="crossbar",linetype="dotted",width=.75,position=position_dodge(0.9))+
  geom_jitter(position = position_jitterdodge(jitter.width = 0.3), alpha = 0.7)+
  labs(y = "Growth rate", x = "Treatment", tag="(f)", color="Treatment")+
  stat_compare_means(comparisons = my_comparisons, label="p.signif",method="t.test", label.y = c(0.9), na=TRUE)+
  scale_color_manual(values = c("deepskyblue", "dodgerblue4"),
                     labels=c("Image"="Image analysis"))+
  scale_y_continuous(limits = c(0,1),
                     breaks = seq(0,1,0.1), expand = c(0,0))+
  theme_bw()+
  theme(axis.title = element_text(size = 12, face = "bold"),
        axis.text = element_text(size=12, colour="black"),
        legend.position = "none",
        plot.tag = element_text(face="bold"),
        legend.title = element_text(face = "bold",size=14), 
        legend.text=element_text(size=12))

Fig_growth_image#Fig 6f
ggsave(plot =Fig_growth_image,  dpi = 600, filename = "Fig_growth_image")

plot_image<-ggarrange(Fig_GR_image,Fig_GP_image,Fig_growth_image, ncol=3)
side_title<-textGrob("Image analysis",gp=gpar(fontsize=18, fontface="bold"),rot=90, just="center")
plot_image<-grid.arrange(side_title,plot_image,ncol=2, widths=c(0.5,10))


##Correlation plots between manual scoring (y-axis) and image analysis data (x-axis)  for the trait ------------------------------------------------------------------

Blues_lab_ID<-read.csv("Blues_lab_ID.csv", header=TRUE)
Blues_lab_ID<-Blues_lab_ID[-1]
Blues_image_ID<-read.csv("Blues_image_ID.csv", header=TRUE)
Blues_image_ID<-Blues_image_ID[-1]

Blues_lab <- Blues_lab_ID%>%
  rename(Germination_lab = Germination_DAS,
         Perc_lab = Healthy_seedlings_Perc)


Blues_image <- Blues_image_ID%>%
  select(c(1,2,3)) %>%
  rename(Germination_image = Germination_DAS,
         Perc_image = Germination_Perc)

plot_data_blues <- merge(Blues_lab , Blues_image, by = "ID", all.x = TRUE)

#Germination_DAS................................................
p1 <- ggplot(plot_data_blues, aes(y=Germination_lab, x=Germination_image))+
  geom_point(aes(color = Type), size=2) +
  stat_cor(method = "pearson", size = 4, fontface = "bold", cor.coef.name = c("r"), geom = "label")+
  stat_smooth(method = lm, color = "black",se = FALSE) +
  labs(tag="(g)",y = " Manual scoring (DAS)", x = " Image analysis (DAS)", color = "Ecotype")+ 
  ggtitle("Germination days after sowing")+
  scale_y_continuous(limits = c(2,9),
                     breaks = seq(2,9,1))+
  scale_x_continuous(limits = c(2,7),
                     breaks = seq(2,9,1))+
  scale_colour_manual(values = c("#B22222","#3B4992FF"))+
  scale_shape_manual(values = c(16,17))+
  theme_bw()+
  theme(axis.title = element_text(size = 14, face = "bold"),
        axis.text.x = element_text(size=12, colour="black"),
        axis.text.y = element_text(size=12, colour="black"),
        plot.tag = element_text(face="bold"),
        legend.title = element_text(face = "bold", size=14), 
        legend.text = element_text( size=12),
        plot.title = element_text (hjust=0.5, size=14,face="bold"))
p1#Fig_6g
ggsave(plot = p1, width =8 , height = 6 , dpi = 600, filename = "Fig1.png")
#Germination_Perc..............................
p2 <- ggplot(plot_data_blues, aes(y=Perc_lab, x=Perc_image))+
  geom_point(aes(color = Type), size=2) +
  stat_cor(method = "pearson", size = 4, fontface = "bold", cor.coef.name = c("r"), geom = "label")+
  stat_smooth(method = lm, color = "black",se = FALSE) +
  labs(tag="(h)",y = " Manual scoring (%)", x = "Image analysis (%)", color = "Ecotype")+
  ggtitle("Germination percentage (%)")+
  scale_y_continuous(limits = c(10,100),
                     breaks = seq(10,100,10))+
  scale_x_continuous(limits = c(40,100),
                     breaks = seq(40,100,10))+
  scale_colour_manual(values = c("#B22222","#3B4992FF"))+
  scale_shape_manual(values = c(16,17))+
  theme_bw()+
  theme(axis.title = element_text(size = 14, face = "bold"),
        axis.text.x = element_text(size=12, colour="black"),
        axis.text.y = element_text(size=12, colour="black"),
        plot.tag = element_text(face="bold"),
        legend.title = element_text(face = "bold", size=14), 
        legend.text = element_text(size=12),
        plot.title = element_text (hjust=0.5, size=14,face="bold"))
p2#Fig_6h
ggsave(plot = p2, width =8 , height = 6 , dpi = 600, filename = "Fig2.png")

Fig_corr<-ggarrange(p1,p2,ncol =2,common.legend=TRUE,legend ="bottom")

side_title<-textGrob("Correlations",gp=gpar(fontsize=18, fontface="bold"),rot=90, just="center")
Fig_corr<-grid.arrange(side_title,Fig_corr,ncol=2, widths=c(0.5,14))

ggsave(plot = Fig_corr, width = 15 , height = 8, dpi = 600, filename = "Fig_6g-h.png")


Fig_6<-ggarrange(plot_manual,plot_image,Fig_corr,nrow=3)
Fig_6
ggsave(Fig_6, dpi=600, filename="Fig_6.png",width=12, height = 12)
#Figure_S5 Boxplots field and lab comparisons------------------------------------------------
Blues_ID_T<-read.csv("Blues_ID_T.csv")
Blues_ID_T <- Blues_ID_T[,-1]
Blues_field <- Blues_ID_T %>%
  select(c(1,2,3,4,18))%>%
  rename(Treatment= Time_Point)

Blues_lab_ID_T<-read.csv("Blues_lab_ID_T.csv")
Blues_lab_ID_T <- Blues_lab_ID_T[,-1]
Blues_lab <- Blues_lab_ID_T %>%
  select(c(1,2,3,5,8)) %>%
  rename(Germination_Perc = Healthy_seedlings_Perc)

Blues_image_ID_T<-read.csv("Blues_image_ID_T.csv")
Blues_image_ID_T  <-Blues_image_ID_T [,-1]
Blues_image <- Blues_image_ID_T %>%
  select(c(1,2,3,4,8))%>%
  mutate(Treatment = recode(Treatment, 
                            "K"= "K_1",
                            "RT"="RT_1")) 


Boxplot_fvsl<-bind_rows(Blues_field,Blues_lab, Blues_image)
Boxplot_fvsl$Treatment <- as.character(Boxplot_fvsl$Treatment)
Boxplot_fvsl<- Boxplot_fvsl %>%
  mutate(Treatment = recode(Treatment, 
                            "T1" = "S1 cold stress (Field)", 
                            "T2" = "S2 (Field)", 
                            "T3" = "S3 (Field)",
                            "K"= "Cold stress (lab)",
                            "RT"="Control (lab)",
                            "K_1"= "Cold stress (image)",
                            "RT_1"="Control (image)"))


#### Germination_DAS------------------------------------------------
Boxplot_fvsl$Type <- factor(Boxplot_fvsl$Type, levels = c("Coastal","Highland"))
Boxplot_fvsl$Treatment <- factor(Boxplot_fvsl$Treatment)
#unique(Boxplot_fvsl$Treatment)
#Boxplot_fvsl <- Boxplot_fvsl %>%
 # mutate(Treatment = factor(Treatment, levels = unique(Treatment))) %>%
 # mutate(Treatment = factor(Treatment, levels = c(
  #  "S1 cold stress (Field)", "S2 (Field)", "S3 (Field)", 
  #  "Cold stress (lab)",  "Control (lab)", 
  #  "Cold stress (image)", "Control (image)"
  #)))

###Germination DAS
# Perform pairwise comparisons
stat_test <- compare_means(
  Germination_DAS ~ Type,
  group.by = "Treatment",
  data = Boxplot_fvsl,
  method = "t.test"
  #method="wilcox.test"
)

# Print results 
print(stat_test)

#  Manually calculate positions for lines and p-values

# Define x-axis positions for dodged boxplots
dodge_width <- 0.8
treatments <- levels(Boxplot_fvsl$Treatment)
n_treatments <- length(treatments)

# Calculate x-coordinates for Highland (left) and Coastal (right) within each Time_Point
x_positions <- data.frame(
  Treatment = rep(treatments, each = 2),
  Type = rep(c( "Coastal", "Highland"), times = n_treatments),
  x = rep(1:n_treatments, each = 2) + c(-dodge_width/2, dodge_width/2)
)

# Define y_positions for horizontal lines (above the data)
y_max <- 32  # Based on your y-axis limit
y_start <- y_max - 2
y_step <- 2  # Spacing between lines
y_positions <- seq(y_start, y_start - (n_treatments - 1) * y_step, by = -y_step)

#Prepare p-value annotations with significance stars
stat_test$p_annot <- ifelse(
  stat_test$p < 0.001, "***",
  ifelse(stat_test$p < 0.01, "**",
         ifelse(stat_test$p < 0.05, "*", "ns"))
)

# Ensure stat_test is ordered to match Time_Point levels
stat_test <- stat_test[order(stat_test$ Treatment), ]


# Create the plot
Fig_gr_compare <- ggplot(Boxplot_fvsl, aes(x =  Treatment, y = Germination_DAS, color = Type)) +
  geom_boxplot(position = position_dodge(dodge_width)) +
  stat_summary(fun = mean, geom = "crossbar",linetype="dotted",width=.75,position = position_dodge(dodge_width)) +
  geom_jitter(position = position_jitterdodge(jitter.width = 0.3, dodge.width = dodge_width), alpha = 0.7) +
  labs(y = "Mean germination time (DAS)", x = "Treatment", tag="(a)",color = "Ecotype") +
  scale_x_discrete(labels=c("S1 cold stress (Field SE)", "S2 (Field SE)","S3 (Field SE)", "Cold stress (lab)","Control(lab)","Cold stress (image)","Control (image)"))+
  scale_color_manual(values = c("#B22222", "#3B4992FF") ) +
  scale_y_continuous(limits = c(0, 35), breaks = seq(0,35,2),expand = c(0, 0)) +
  theme_bw() +
  theme(
    axis.title.x = element_blank(),
    plot.tag=element_text(face = "bold"),
    axis.text.x = element_text(size = 12, colour = "black"),
    axis.text.y = element_text(size = 12, colour = "black"),
    axis.title.y = element_text(size = 14, face = "bold",colour = "black"),
    legend.title = element_text(face = "bold",size=14), 
    legend.text=element_text( size=12),
    legend.position = "bottom"
  ) +
  
  # Add horizontal lines and p-values manually
  geom_segment(
    data = data.frame(
      Treatment = treatments,
      x_start = 1:n_treatments - dodge_width/2,
      x_end = 1:n_treatments + dodge_width/2,
      y = y_positions
    ),
    aes(x = x_start, xend = x_end, y = y, yend = y),
    color = "black",
    inherit.aes = FALSE
  ) +
  geom_text(
    data = data.frame(
      Treatment = treatments,
      x = 1:n_treatments,
      y = y_positions + 1,  # Slightly above the line
      label = paste("p =", stat_test$p.format, stat_test$p_annot)
    ),
    aes(x = x, y = y, label = label),
    color = "black",
    size = 3.5,
    inherit.aes = FALSE
  )
# Display the plot
print(Fig_gr_compare)

###Germination Percentage
# Perform pairwise comparisons
# Perform pairwise comparisons
stat_test <- compare_means(
  Germination_Perc ~ Type,
  group.by = "Treatment",
  data = Boxplot_fvsl,
  method = "t.test"
  #method="wilcox.test"
)

# Print results 
print(stat_test)

##Germination_Perc--------------------------------------------
#  Manually calculate positions for lines and p-values

# Define x-axis positions for dodged boxplots
dodge_width <- 0.8
treatments <- levels(Boxplot_fvsl$Treatment)
n_treatments <- length(treatments)

# Calculate x-coordinates for Highland (left) and Coastal (right) within each Time_Point
x_positions <- data.frame(
  Treatment = rep(treatments, each = 2),
  Type = rep(c( "Coastal", "Highland"), times = n_treatments),
  x = rep(1:n_treatments, each = 2) + c(-dodge_width/2, dodge_width/2)
)

# Define y_positions for horizontal lines (above the data)
y_max <- 110  # Based on your y-axis limit
y_start <- y_max - 2
y_step <- 0  # Spacing between lines
y_positions <- seq(y_start, y_start - (n_treatments - 1) * y_step, by = -y_step)

#  Prepare p-value annotations with significance stars
stat_test$p_annot <- ifelse(
  stat_test$p < 0.001, "***",
  ifelse(stat_test$p < 0.01, "**",
         ifelse(stat_test$p < 0.05, "*", "ns"))
)

# Ensure stat_test is ordered to match Time_Point levels
stat_test <- stat_test[order(stat_test$ Treatment), ]


#  Create the plot
Fig_gp_compare <- ggplot(Boxplot_fvsl, aes(x =  Treatment, y = Germination_Perc, color = Type)) +
  geom_boxplot(position = position_dodge(dodge_width)) +
  stat_summary(fun = mean, geom = "crossbar",linetype="dotted",width=.75,position = position_dodge(dodge_width)) +
  geom_jitter(position = position_jitterdodge(jitter.width = 0.3, dodge.width = dodge_width), alpha = 0.7) +
  labs(y = "Germination Percentage (%)", x = "Treatment", tag="(b)",color = "Ecotype") +
  scale_x_discrete(labels=c("S1 cold stress (Field EP)", "S2 (Field EP)","S3 (Field EP)", "Cold stress (lab)","Control(lab)","Cold stress (image)","Control (image)"))+
  scale_color_manual(values = c("#B22222", "#3B4992FF") ) +
  scale_y_continuous(limits = c(0, 120), breaks = seq(0,120,10),expand = c(0, 0)) +
  theme_bw() +
  theme(
    #axis.title.x = element_blank(),
    axis.text.x = element_text(size = 12, colour = "black"),
    axis.text.y = element_text(size = 12, colour = "black"),
    axis.title.y = element_text(size = 14, face = "bold",colour = "black"),
    axis.title.x  = element_text(size = 14, face = "bold",colour = "black"),
    plot.tag=element_text(face = "bold"),
    legend.title = element_text(face = "bold",size=14), 
    legend.text=element_text( size=12),
    legend.position = "bottom"
  ) +
  
  # Add horizontal lines and p-values manually
  geom_segment(
    data = data.frame(
      Treatment = treatments,
      x_start = 1:n_treatments - dodge_width/2,
      x_end = 1:n_treatments + dodge_width/2,
      y = y_positions
    ),
    aes(x = x_start, xend = x_end, y = y, yend = y),
    color = "black",
    inherit.aes = FALSE
  ) +
  geom_text(
    data = data.frame(
      Treatment = treatments,
      x = 1:n_treatments,
      y = y_positions + 3,  # Slightly above the line
      label = paste("p =", stat_test$p.format, stat_test$p_annot)
    ),
    aes(x = x, y = y, label = label),
    color = "black",
    size = 3.5,
    inherit.aes = FALSE
  )
# Display the plot
print(Fig_gp_compare)

Fig_comparison<-ggarrange(Fig_gr_compare,Fig_gp_compare,nrow=2,common.legend=TRUE,legend ="bottom")
Fig_comparison

ggsave(plot = Fig_comparison, width = 17,height = 11, dpi = 600, filename = "Fig_comparison.png")

##Correlations between percentage field and lab---------------------------------------------
Blues_ID_T <- read_csv("Blues_ID_T.csv")
Blues_ID_T <- Blues_ID_T[,-1]
Blues_field1 <- Blues_ID_T %>%
  filter(Time_Point == "T1") %>%
  select(c(1,3,4,18)) %>%
  rename(Germination_field = Germination_DAS,
         Perc_field = Germination_Perc)

Blues_lab_ID_T <- read_csv("Blues_lab_ID_T.csv")
Blues_lab_ID_T <- Blues_lab_ID_T[,-1]
Blues_lab1 <- Blues_lab_ID_T %>%
  filter(Treatment == "K") %>%
  select(c(1:3,4,5)) %>%
  rename(Germination_lab = Germination_DAS,
         Viable_Perc_lab = Healthy_seedlings_Perc,
         Perc_lab=Germination_Perc
  )

plot_data_blues <- merge(Blues_field1, Blues_lab1, by = "ID", all.x = TRUE)


corr_fvsl <- ggplot(plot_data_blues, aes(Perc_lab, Perc_field))+
  geom_point(aes(color = Type), size=2) +
  stat_cor(method = "pearson", size = 4, fontface = "bold", cor.coef.name = c("r"), geom = "label")+
  stat_smooth(method = lm, color = "black",se = FALSE) +
  labs(y = "Emergence percentage (Field) (%)", x = "Germination percentage (Lab) (%)", tag= "(c)",color = "Ecotype", 
       shape = "Ecotype")+ 
  scale_y_continuous(limits = c(10,60),
                     breaks = seq(10,60,10))+
  scale_x_continuous(limits = c(50,100),
                     breaks = seq(50,100,10))+
  scale_colour_manual(values = c("#B22222","#3B4992FF"))+
  scale_shape_manual(values = c(16,17))+
  theme_bw()+
  theme(axis.title = element_text(size = 14, face = "bold"),
        axis.text.x = element_text(size=12, colour="black"),
        axis.text.y = element_text(size=12, colour="black"),
        plot.tag = element_text(face="bold"),
        legend.title = element_text(face = "bold", size=14), 
        legend.text = element_text( size=12),
        plot.title = element_text (hjust=0.5, size=14,face="bold"))
corr_fvsl

corr_fvsvl <- ggplot(plot_data_blues, aes(Viable_Perc_lab, Perc_field))+
  geom_point(aes(color = Type), size=2) +
  stat_cor(method = "pearson", size = 4, fontface = "bold", cor.coef.name = c("r"), geom = "label")+
  stat_smooth(method = lm, color = "black",se = FALSE) +
  labs(y = "Emergence percentage (Field) (%)", x = "Viable germination percentage (Lab) (%)", tag= "(d)",color = "Ecotype", 
       shape = "Ecotype")+ 
  scale_y_continuous(limits = c(10,60),
                     breaks = seq(10,60,10))+
  scale_x_continuous(limits = c(10,100),
                     breaks = seq(10,100,10))+
  scale_colour_manual(values = c("#B22222","#3B4992FF"))+
  scale_shape_manual(values = c(16,17))+
  theme_bw()+
  theme(axis.title = element_text(size = 14, face = "bold"),
        axis.text.x = element_text(size=12, colour="black"),
        axis.text.y = element_text(size=12, colour="black"),
        plot.tag = element_text(face="bold"),
        legend.title = element_text(face = "bold", size=14), 
        legend.text = element_text( size=12),
        plot.title = element_text (hjust=0.5, size=14,face="bold"))
corr_fvsvl

Fig_corr_compare<-ggarrange(corr_fvsl,corr_fvsvl,ncol=2,common.legend=TRUE,legend ="bottom")
Fig_corr_compare

ggsave(plot = Fig_corr_compare, width = 15,height = 8, dpi = 600, filename = "Fig_corr_compare.png")


# Figure_S7 Cumulative germination---------------------------------------------------------------
Cummulative_cold<-read_ods("Cummulative.ods", sheet = "Cold")
Cummulative_RT<-read_ods("Cummulative.ods", sheet = "RT")
Cummulative_cold$Total_Cumulative_Germination<-rowSums(Cummulative_cold[,-1])
Cummulative_RT$Total_Cumulative_Germination<-rowSums(Cummulative_RT[,-1])


RT_1<-ggplot(Cummulative_RT,aes(x=DAS, y=Total_Cumulative_Germination))+
  geom_line(color="skyblue")+
  geom_point()+
  labs(x="Days after sowing (DAS)", y="Cumulative Gerimation Count (RT)", tag="(a)")+
  scale_y_continuous(limits = c(0,600),
                     breaks = seq(0,600,100), expand = c(0,0))+
  scale_x_continuous(limits = c(0,31),
                     breaks = seq(0,30,1), expand = c(0,0))+
  theme_bw()+
  theme(axis.title.x = element_text(size = 14, face = "bold"),
        axis.text.x = element_text(size=12, colour="black"),
        axis.text.y = element_text(size=12, colour="black"),
        plot.tag=element_text(face="bold"),
        axis.title.y = element_text(size = 14, face = "bold"),
  )

RT_1 
Cold_1<-ggplot(Cummulative_cold,aes(x=DAS, y=Total_Cumulative_Germination))+
  geom_line(color="skyblue")+
  geom_point()+
  labs(x="Days after sowing (DAS)", y="Cumulative Gerimation Count (Cold)", tag="(b)")+
  scale_y_continuous(limits = c(0,600),
                     breaks = seq(0,600,100), expand = c(0,0))+
  scale_x_continuous(limits = c(0,31),
                     breaks = seq(0,30,1), expand = c(0,0))+
  theme_bw()+
  theme(axis.title.x = element_text(size = 14, face = "bold"),
        axis.text.x = element_text(size=12, colour="black"),
        axis.text.y = element_text(size=12, colour="black"),
        plot.tag=element_text(face="bold"),
        axis.title.y = element_text(size = 14, face = "bold"),
  )
Cold_1

Cumulative_line<-ggarrange(RT_1, Cold_1, nrow=2)
Cumulative_line
ggsave(plot = Cumulative_line, width = 17,height = 11, dpi = 600, filename = "Cumulative_line.png")

#SEED SIZE analysis (MARViN measurements and image analysis)---------------------------------------

#1. Data preparation ---------------------------------------------------------------

Marvin_data<- readODS::read_ods("marvin-data-processed.ods")
#image_data<-read.delim("seedsize_image.txt",header=TRUE,sep=",")
subset_dataset <- Marvin_data[complete.cases(Marvin_data$ID), ]
subset_dataset1 <- subset_dataset[,c(5,13,14,15,16,17,18,20)]
subset_dataset1<-subset_dataset1%>%relocate(ID,line_name,Experiment, .before=TKW_g)
subset_dataset1 <- subset_dataset1[subset_dataset1$ID != "A60", ]
subset_KAUST<-subset_dataset1%>%filter(Experiment == "KAUST")
subset_KAUST <- subset_KAUST[complete.cases(subset_KAUST$ID), ]
subset_dataset2 <- subset_dataset1[subset_dataset1$Experiment != "KAUST", ]


write.csv(subset_dataset2, "subset_dataset.csv")
write.csv(subset_KAUST, "subset_KAUST.csv")
## 1.1 Relative values-------------------------------------------------------
###KAUST
str(subset_KAUST)
subset_KAUST$TKW_g <- as.numeric(as.character(subset_KAUST$TKW_g))
tkw_mean <-mean(subset_KAUST$TKW_g)
subset_KAUST$TKW_relative_value <-  subset_KAUST$TKW_g - tkw_mean 

area_mean <-mean(subset_KAUST$area_mean)
subset_KAUST$area_relative_value <-  subset_KAUST$area_mean -area_mean 
subset_KAUST$Data<-"KAUST"
###Others
str(subset_dataset2)
subset_dataset2$TKW_g <- as.numeric(as.character(subset_dataset2$TKW_g))
tkw_mean1 <-mean(subset_dataset2$TKW_g, na.rm = TRUE)
subset_dataset2$TKW_relative_value <-  subset_dataset2$TKW_g -tkw_mean1 
subset_dataset2$Data<-"Others"

area_mean1 <-mean(subset_dataset2$area_mean)
subset_dataset2$area_relative_value <-  subset_dataset2$area_mean-area_mean1 


relative_values<-rbind(subset_dataset2,subset_KAUST)

origin_data_saponin<-read.csv("acc_seedsize.csv",header = TRUE)
origin_data_saponin$ID <- gsub("H21_", "", origin_data_saponin$ID)
origin_data_image <- origin_data_saponin[origin_data_saponin$ID != "A60", ]
relative_values<- merge(relative_values,origin_data_saponin, by = c("ID"))

## Average data marvin---------------------------------------------------
### Marvin 
avg_data_marvin <- relative_values %>%
  group_by(ID, Experiment, generation, Data) %>%
  summarise_at(vars(TKW_relative_value, area_relative_value),
               .funs = mean,
               na.rm = TRUE)
avg_data_marvin<- merge(avg_data_marvin,origin_data_saponin, by = c("ID"))
str(avg_data_marvin)
write.csv(avg_data_marvin,"avg_data_marvin")

##1.2 Image Analysis ---------------------------------------------------

data_corrected2<-read.csv("corrected-output-8C.csv")
data_corrected2<-data_corrected2%>%select(4,5,12,14)
data_corrected2<-data_corrected2%>%rename("Rep"="imgname_rep",
                                          "ID"="imgname_accession")
data_corrected2<-data_corrected2%>%filter(class == "seed")
avg_data_corrected2 <- data_corrected2%>%
  group_by(ID,Rep) %>%
  summarise_at(vars(area_cm2),
               .funs = mean,
               na.rm = TRUE)
avg_data_corrected2<- merge(avg_data_corrected2,origin_data_saponin, by = c("ID"))
avg_data_corrected2$area_mm2<-avg_data_corrected2$area_cm2*100

avg_data_corrected2$ID <- as.factor(as.character(avg_data_corrected2$ID))
avg_data_corrected2$Rep <- as.factor(as.character(avg_data_corrected2$Rep))

## 1.3 Subset (generation 1) Heidfeldhof --------------------------------------------
Heidfeldhof_marvin<-subset_dataset1%>%filter(Experiment == "Heidfeldhof2018")

# 2. Seed size Relative values Marvin -------
##2.1 BLUPs_relative values -----------------------------------------------------------------
str(avg_data_marvin)

avg_data_marvin$ID <- as.factor(as.character(avg_data_marvin$ID))
avg_data_marvin$Experiment <- as.factor(as.character(avg_data_marvin$Experiment))
avg_data_marvin$Type <- as.factor(as.character(avg_data_marvin$Type))
traits <- names(avg_data_marvin[c(5,6)])
avg_data_marvin$Experiment

traits

var <- ID <-list(traits)


for (i in traits)
{
  mod_blup <- asreml(as.formula(paste(i,"~ 1+ Experiment ")),
                     random= ~ID,
                     data = avg_data_marvin)  
  
  pred_blups_ID <- predict(mod_blup, classify = "ID")
  
  bp_ID <- pred_blups_ID$pvals
  bp_ID <- bp_ID %>%
    dplyr::select(c(1:2))
  
  Variance <- as.data.frame(summary(mod_blup)$varcomp)
  Variance <- Variance %>%
    mutate(avsed = pred_blups_ID$avsed,
           h2 = c(1-(((pred_blups_ID$avsed)^2)/(2*(Variance[1,1])))))
  
  var[[paste(i)]] <- Variance
  ID[[paste(i)]] <- bp_ID
  
}

Ran_var <- do.call(rbind.data.frame, var)
Ran_var <- Ran_var[-1,]

Ran_var$Factor <- rownames(Ran_var)
Ran_var <- Ran_var %>%
  remove_rownames()%>%
  relocate(Factor, .before = component) %>%
  separate(Factor, c("Trait", "Factor"), sep = "([.])", extra = "merge")%>%
  mutate_if(is.numeric, round, digits = 2)

Var_h2 <- Ran_var %>%
  distinct(Trait, .keep_all = TRUE)

ID <- ID[-1]

Blups_ID <- cbind(ID$TKW_relative_value, ID$area_relative_value$predicted.value)

names(Blups_ID)[2:3] <- traits

traits
Blups_ID <- merge(Blups_ID, origin_data_saponin, by = "ID")

write.csv(Blups_ID,"Blups_image_ID.csv", na="NA")


### 2.1.1 Heritability ----------------------------------------------------------
Var_h2$component<-as.numeric(as.character(Var_h2$component))
Var_h2$avsed<-as.numeric(as.character(Var_h2$avsed))
Var_h2$h2<-as.numeric(as.character(Var_h2$h2))

Var_h2<- Var_h2%>%
  mutate_if(is.numeric,round,digits=2)

## 2.2 BLUEs_relative_values--------------------------------------------------------------------------

traits <- names(avg_data_marvin[c(5,6)])
traits

an <- ID <- ID_E <-list(traits)


for (i in traits)
{
  mod_blue <- asreml(as.formula(paste(i,"~ 1+ ID+ Experiment")),
                     data = avg_data_marvin)    
  
  pred_blues_ID <- predict(mod_blue, classify = "ID")
  pred_blues_IDxE <- predict(mod_blue, classify = "ID:Experiment")
  
  blues_ID <- pred_blues_ID$pvals
  blues_ID <- blues_ID %>%
    dplyr::select(c(1:2))
  
  
  blues_IDxE<- pred_blues_IDxE$pvals
  blues_IDxE<- blues_IDxE %>%
    dplyr::select(c(1:3))
  
  #Variance <- as.data.frame(summary(mod_blue)$varcomp)
  anova <- as.data.frame(wald(mod_blue))
  
  an[[paste(i)]] <- anova
  ID[[paste(i)]] <- blues_ID
  ID_E[[paste(i)]] <- blues_IDxE
  
}

anv <- do.call(rbind.data.frame, an)
anv <- anv[-1,]
anv$Factor <- rownames(anv)
anv <- anv %>%
  remove_rownames()%>%
  relocate(Factor, .before = Df) %>%
  separate(Factor, c("Trait", "Factor"), sep = "([.])", extra = "merge")

ID <- ID[-1]

Blues_marvin_ID <- cbind(ID$TKW_relative_value, ID$area_relative_value$predicted.value)
names(Blues_marvin_ID)[2:3] <- traits

Blues_marvin_ID <- merge(Blues_marvin_ID, origin_data_saponin, by = "ID")

write.csv(Blues_marvin_ID,"Blues_marvin_ID.csv")

#ID_E <- ID_E[-1]

#Blues_marvin_ID_E <- cbind(ID_E$TKW_relative_value, ID_E$area_relative_value$predicted.value)

#names(Blues_marvin_ID_E)[3:6] <- traits

#Blues_marvin_ID_E <- merge(Blues_marvin_ID_E, origin_data_saponin, by = "ID")

# Figure 7a Boxplot ecotype---------------------------------------------------------------------
Fig_area_Marvin <- ggplot(Blues_marvin_ID,aes(x=Type, y = area_relative_value,color=Type))+
  geom_boxplot()+
  stat_summary(fun=mean, geom="crossbar", linetype="dotted", width=.75)+
  geom_point(position = position_jitterdodge(jitter.width = 0.2), alpha = 0.7)+
  labs(y = "Seed size (area mm2)", x = "Ecotype",tag = "(a)",color = "Ecotype")+
  ggtitle("MARViN measurements (relative values)")+
  scale_color_manual(values = c("#B22222","#3B4992FF"))+ 
  #scale_fill_manual(values = wes_palette("Darjeeling1", n = 5))+
  scale_y_continuous(limits = c(-1,1),
                     breaks = seq(-1,1.0,0.5), expand = c(0,0))+
  theme_bw()+
  theme(axis.title.x = element_text(size = 14, face = "bold"),
        axis.text.x = element_text(size=12, colour="black"),
        axis.text.y = element_text(size=12, colour="black"),
        plot.tag=element_text(face="bold"),
        axis.title.y = element_text(size = 14, face = "bold"),
        legend.position = "bottom",
        legend.title=element_text(face="bold",size=14), 
        legend.text=element_text(size=12),
        plot.title = element_text (hjust=0.5, size=18,face="bold"))
Fig_area_Marvin
compare_means(data=Blues_marvin_ID,area_relative_value~Type)
test<-t.test (data=Blues_marvin_ID, area_relative_value~Type)
Fig_area_Marvin<- Fig_area_Marvin+ stat_compare_means (comparisons =list(c("Coastal", "Highland")), 
                                                       label="p.signif",method="t.test", ref.group="Coastal")

Fig_area_Marvin
ggsave(plot = Fig_area_Marvin, width =8 , height = 6 , dpi = 600, filename = "Fig_area_Marvin.png")

stat_test<-compare_means(area_relative_value~Type,
                         data=Blues_marvin_ID,
                         method="t.test")
print(stat_test)
aggregate(area_relative_value~Type,data=Blues_marvin_ID,FUN=mean)

# Figure 7b Saponin---------------------------------
Fig_area_Marvin_Saponin <- ggplot(Blues_marvin_ID,aes(x=Saponin, y = area_relative_value,color=Saponin))+
  geom_boxplot(aes(middle=mean(area_relative_value)))+
  geom_point(position = position_jitterdodge(jitter.width = 0.2), alpha = 0.7)+
  labs(y = "Seed_size(area mm2)", x = "Saponin", tag="(b)")+
  ggtitle("MARViN measurements (relative values)")+
  scale_color_manual(values = c("Bitter" = "#3B4992FF", "Sweet" = "#E69F00"))+ 
  #scale_fill_manual(values = wes_palette("Darjeeling1", n = 5))+
  scale_y_continuous(limits = c(-1,1),
                     breaks = seq(-1,1.0,0.5), expand = c(0,0))+
  theme_bw()+
  theme(axis.title.x = element_text(size = 14, face = "bold"),
        axis.text.x = element_text(size=12, colour="black"),
        axis.text.y = element_text(size=12, colour="black"),
        plot.tag=element_text(face="bold"),
        axis.title.y = element_text(size = 14, face = "bold"),
        legend.position = "bottom",
        legend.title=element_text(size=12), 
        legend.text=element_text(size=10),
        plot.title = element_text (hjust=0.5, size=18,face="bold"))
Fig_area_Marvin_Saponin
compare_means(data=Blues_marvin_ID,area_relative_value~Saponin)
test<-t.test (data=Blues_marvin_ID, area_relative_value~Saponin)
Fig_area_Marvin_Saponin<- Fig_area_Marvin_Saponin+ stat_compare_means (comparisons =list(c("Bitter", "Sweet")), 
                                                                       label="p.signif",method="t.test", ref.group="Bitter")

Fig_area_Marvin_Saponin
ggsave(plot = Fig_area_Marvin_Saponin, width =8 , height = 6 , dpi = 600, filename = "Fig_area_Marvin_Saponin.png")
Fig_area_comparisons<-ggarrange(Fig_area_Marvin,Fig_area_Marvin_Saponin, ncol=2,legend ="bottom")
ggsave(plot = Fig_area_comparisons, width =16 , height = 6 , dpi = 600, filename = "Fig_area_comparisons.png")

# 3.Marvin seed size for Generation 1 (Heidfeldhof)------------------------------------------------------------------
## 3.1 Blups ------------------------------
Heidfeldhof_marvin$ID <- as.factor(as.character(Heidfeldhof_marvin$ID))

traits <- names(Heidfeldhof_marvin[c(5)])
traits

an <- ID <- list(traits)

for (i in traits)
{
  mod_blup <- asreml(as.formula(paste(i,"~ 1 ")),
                     random= ~ID,
                     data =Heidfeldhof_marvin )  
  
  pred_blups_ID <- predict(mod_blup, classify = "ID")
  
  bp_ID <- pred_blups_ID$pvals
  bp_ID <- bp_ID %>%
    dplyr::select(c(1:2))
  
  Variance <- as.data.frame(summary(mod_blup)$varcomp)
  Variance <- Variance %>%
    mutate(avsed = pred_blups_ID$avsed,
           h2 = c(1-(((pred_blups_ID$avsed)^2)/(2*(Variance[1,1])))))
  
  var[[paste(i)]] <- Variance
  ID[[paste(i)]] <- bp_ID
  
}

Ran_var <- do.call(rbind.data.frame, var)
Ran_var <- Ran_var[-1,]

Ran_var$Factor <- rownames(Ran_var)
Ran_var <- Ran_var %>%
  remove_rownames()%>%
  relocate(Factor, .before = component) %>%
  separate(Factor, c("Trait", "Factor"), sep = "([.])", extra = "merge")%>%
  mutate_if(is.numeric, round, digits = 2)

Var_h2_1<- Ran_var %>%
  distinct(Trait, .keep_all = TRUE)

ID <- ID[-1]

Blups_ID_1 <- cbind( ID$area_mean$predicted.value)

names(Blups_ID_1)[2] <- traits

traits
#Blups_ID_1 <- merge(Blups_ID_1, origin_data_image, by = "ID")

#write.csv(Blups_ID_1,"Blups_ID_1.csv", na="NA")


### Heritability ----------------------------------------------------------
Var_h2_1 $component<-as.numeric(as.character(Var_h2_1 $component))
Var_h2_1 $avsed<-as.numeric(as.character(Var_h2_1 $avsed))
Var_h2_1 $h2<-as.numeric(as.character(Var_h2_1 $h2))

Var_h2_1 <- Var_h2_1 %>%
  mutate_if(is.numeric,round,digits=2)

write.csv(Var_h2_1, file="Heritability.csv")

## 3.2  BLUEs-----------------------------------------
Heidfeldhof_marvin$ID <- as.factor(as.character(Heidfeldhof_marvin$ID))

traits <- names(Heidfeldhof_marvin[c(5)])
traits

an <- ID <- list(traits)


for (i in traits)
  
{
  mod_blue <- asreml(as.formula(paste(i,"~ 1+ ID")),
                     data = Heidfeldhof_marvin)    
  
  pred_blues_ID <- predict(mod_blue, classify = "ID")
  
  blues_ID <- pred_blues_ID$pvals
  blues_ID <- blues_ID %>%
    dplyr::select(c(1:2))
  
  
  #Variance <- as.data.frame(summary(mod_blue)$varcomp)
  anova <- as.data.frame(wald(mod_blue))
  
  an[[paste(i)]] <- anova
  ID[[paste(i)]] <- blues_ID
  
}

anv <- do.call(rbind.data.frame, an)
anv <- anv[-1,]
anv$Factor <- rownames(anv)
anv <- anv %>%
  remove_rownames()%>%
  relocate(Factor, .before = Df) %>%
  separate(Factor, c("Trait", "Factor"), sep = "([.])", extra = "merge")

Blues_Heidfeldhof_marvin_ID <- cbind(ID$area_mean)

names(Blues_Heidfeldhof_marvin_ID )[2] <- traits

Blues_gen1_marvin_ID <- merge(Blues_Heidfeldhof_marvin_ID, origin_data_image, by = "ID")
write.csv(Blues_gen1_marvin_ID,"Blues_gen1_marvin_ID.csv")

# Figure 7c Boxplot----------------------------------------------
marvin1 <- ggplot(Blues_gen1_marvin_ID,aes(x=Type, y = area_mean,color=Type))+
  geom_boxplot()+
  stat_summary(fun=mean, geom="crossbar", linetype="dotted", width=.75)+
  geom_point(position = position_jitterdodge(jitter.width = 0.2), alpha = 0.7)+
  labs(y = "Seed size(area mm2)", x = "Ecotype", tag="(b)",color = "Ecotype")+
  ggtitle("MARViN measurements")+
  scale_color_manual(values = c("#B22222","#3B4992FF"))+ 
  #scale_fill_manual(values = wes_palette("Darjeeling1", n = 5))+
  scale_y_continuous(limits = c(0.5,3),
                     breaks = seq(0.5,3,0.5), expand = c(0,0))+
  theme_bw()+
  theme(axis.title.x = element_text(size = 14, face = "bold"),
        axis.text.x = element_text(size=12, colour="black"),
        axis.text.y = element_text(size=12, colour="black"),
        plot.tag=element_text(face="bold"),
        axis.title.y = element_text(size = 14, face = "bold"),
        legend.position = "bottom",
        legend.title=element_text(face="bold",size=14), 
        legend.text=element_text(size=12),
        plot.title = element_text (hjust=0.5, size=18,face="bold"))
compare_means(data=Blues_gen1_marvin_ID,area_mean~Type)
test<-t.test (data=Blues_gen1_marvin_ID, area_mean~Type)
marvin1<- marvin1+ stat_compare_means (comparisons =list(c("Coastal", "Highland")), 
                                       label="p.signif",method="t.test", ref.group="Coastal")

marvin1
ggsave(plot = marvin1 , width =8 , height = 6 , dpi = 600, filename = "marvin1.png")


#4. Seed size from Image analysis----------
## 4.1 Blups and heritability---------------------------------------------------------------------
traits <- names(avg_data_corrected2[c(4)])
traits

an <- ID <- ID_R <-list(traits)


for (i in traits)
{
  mod_blup <- asreml(as.formula(paste(i,"~ 1+ Rep ")),
                     random= ~ID,
                     data = avg_data_corrected2)  
  
  pred_blups_ID <- predict(mod_blup, classify = "ID")
  
  bp_ID <- pred_blups_ID$pvals
  bp_ID <- bp_ID %>%
    dplyr::select(c(1:2))
  
  Variance <- as.data.frame(summary(mod_blup)$varcomp)
  Variance <- Variance %>%
    mutate(avsed = pred_blups_ID$avsed,
           h2 = c(1-(((pred_blups_ID$avsed)^2)/(2*(Variance[1,1])))))
  
  var[[paste(i)]] <- Variance
  ID[[paste(i)]] <- bp_ID
  
}

Ran_var <- do.call(rbind.data.frame, var)
Ran_var <- Ran_var[-1,]

Ran_var$Factor <- rownames(Ran_var)
Ran_var <- Ran_var %>%
  remove_rownames()%>%
  relocate(Factor, .before = component) %>%
  separate(Factor, c("Trait", "Factor"), sep = "([.])", extra = "merge")%>%
  mutate_if(is.numeric, round, digits = 2)

Var_h2_image_c <- Ran_var %>%
  distinct(Trait, .keep_all = TRUE)

ID <- ID[-1]

Blups_image_c_ID <- cbind( ID$area_mm2$predicted.value)

names(Blups_image_c_ID)[2] <- traits

traits
B#lups_image_c_ID <- merge(Blups_image_c_ID, origin_data_image, by = "ID")

#write.csv(Blups_image_c_ID,"Blups_image_c_ID.csv", na="NA")


### Heritability ----------------------------------------------------------
Var_h2_image_c $component<-as.numeric(as.character(Var_h2_image_c $component))
Var_h2_image_c $avsed<-as.numeric(as.character(Var_h2_image_c $avsed))
Var_h2_image_c $h2<-as.numeric(as.character(Var_h2_image_c $h2))

Var_h2_image_c <- Var_h2_image_c %>%
  mutate_if(is.numeric,round,digits=2)

## 4.2 BLUEs-----------------------------------------
traits <- names(avg_data_corrected2[c(4)])
traits

an <- ID <- list(traits)


for (i in traits)
  
{
  mod_blue <- asreml(as.formula(paste(i,"~ 1+ Rep+ ID")),
                     data = avg_data_corrected2)    
  
  pred_blues_ID <- predict(mod_blue, classify = "ID")
  
  blues_ID <- pred_blues_ID$pvals
  blues_ID <- blues_ID %>%
    dplyr::select(c(1:2))
  
  
  #Variance <- as.data.frame(summary(mod_blue)$varcomp)
  anova <- as.data.frame(wald(mod_blue))
  
  an[[paste(i)]] <- anova
  ID[[paste(i)]] <- blues_ID
  
}

anv <- do.call(rbind.data.frame, an)
anv <- anv[-1,]
anv$Factor <- rownames(anv)
anv <- anv %>%
  remove_rownames()%>%
  relocate(Factor, .before = Df) %>%
  separate(Factor, c("Trait", "Factor"), sep = "([.])", extra = "merge")

Blues_avg_data_corrected2 <- cbind(ID$area_mm2)

names(Blues_avg_data_corrected2 )[2] <- traits

Blues_image_seedsize_ID <- merge(Blues_avg_data_corrected2, origin_data_image, by = "ID")
write.csv(Blues_image_seedsize_ID, "Blues_image_seedsize_ID .csv")
#Figure 7d Boxplot------------------------------------------
Fig_corrected2 <- ggplot(Blues_image_seedsize_ID,aes(x=Type, y = area_mm2,color=Type))+
  geom_boxplot()+
  ggtitle("Image analysis")+
  stat_summary(fun=mean, geom="crossbar", linetype="dotted", width=.75)+
  geom_point(position = position_jitterdodge(jitter.width = 0.2), alpha = 0.7)+
  labs(y = "Seed size(area mm2)", x = "Ecotype", tag="(c)",color = "Ecotype")+
  scale_color_manual(values = c("#B22222","#3B4992FF"))+ 
  #scale_fill_manual(values = wes_palette("Darjeeling1", n = 5))+
  scale_y_continuous(limits = c(3,7),
                     breaks = seq(3,7,0.5), expand = c(0,0))+
  theme_bw()+
  theme(axis.title.x = element_text(size = 14, face = "bold"),
        axis.text.x = element_text(size=12, colour="black"),
        axis.text.y = element_text(size=12, colour="black"),
        plot.tag=element_text(face="bold"),
        axis.title.y = element_text(size = 14, face = "bold"),
        legend.position = "bottom",
        legend.title=element_text(face="bold",size=14), 
        legend.text=element_text(size=12),
        plot.title = element_text (hjust=0.5, size=18,face="bold"))
Fig_corrected2
compare_means(data=Blues_image_seedsize_ID,area_mm2~Type)
test<-t.test (data=Blues_image_seedsize_ID, area_mm2~Type)
Fig_image_seedsize<- Fig_corrected2+ stat_compare_means (comparisons =list(c("Coastal", "Highland")), 
                                               label="p.signif",method="t.test", ref.group="Coastal")
Fig_image_seedsize
ggsave(plot = Fig_corrected2, width =8 , height = 6 , dpi = 600, filename = "Fig_corrected_0.95.png")

seed_size<-ggarrange(Fig_area_Marvin, Fig_area_Marvin_Saponin, marvin1,Fig_image_seedsize, ncol=2,nrow=2)
seed_size #Figure 7a-d
ggsave(plot =seed_size, width =12 , height = 9 ,dpi = 600, filename = "Seed_size.png")

#Figure 7e Correlations------------------------------------------------------------------

Blues_avg_image_seedsize <-Blues_image_seedsize_ID%>%
  rename(area_mean_image=area_mm2)
#Blues_avg_image_seedsize <- Blues_avg_image_seedsize[Blues_image_seedsize$ID != "A60", ]

Blues_gen1_marvin <-Blues_gen1_marvin_ID%>%
  select(c(1,2)) %>%
  rename(area_mean_marvin=area_mean)

plot_marvin_blues <- merge( Blues_avg_image_seedsize, Blues_gen1_marvin , by = "ID", all.x = TRUE)

plot_marvin_corr<-plot_marvin_blues[,c(1,2,3,5,9,10)]

Corr_seedsize <- ggplot(plot_marvin_corr, aes(area_mean_marvin, area_mean_image))+
  geom_point(aes(color = Type), size = 3 ) +
  stat_cor(method = "pearson", size = 4, fontface = "bold", cor.coef.name = c("r"), geom = "label")+
  stat_smooth(method = lm, color = "black",se = FALSE) +
  labs(x = " MARViN Seed size (mm2)", y = " Image analysis Seed size (mm2)", color = "Ecotype", tag="(d)")+ 
  ggtitle("Correlation")+
  scale_y_continuous(limits = c(3.5,7),
                     breaks = seq(3.5,7.5,0.5))+
  scale_x_continuous(limits = c(1,2.7),
                     breaks = seq(1,2.7,0.5))+
  scale_colour_manual(values = c("#B22222","#3B4992FF"))+
  theme_bw()+
  theme(axis.title = element_text(size = 14, face = "bold"),
        axis.text.x = element_text(size=12, colour="black"),
        axis.text.y = element_text(size=12, colour="black"),
        plot.tag = element_text(face="bold"),
        legend.title=element_text(size=12), 
        legend.text=element_text(size=10),
        legend.position  = "bottom",
        plot.title = element_text (hjust=0.5, size=18,face="bold"))

Corr_seedsize# Figure 7e
ggsave(plot = Corr_seedsize, width =8 , height = 6 ,dpi = 600, filename = "Seed_size_corr_3.png")


#Figure_S4 Pearson’s correlation coefficients from laboratory germination experiment--------------------
Blues_lab_ID <- read_csv("Blues_lab_ID.csv")
Blues_lab_ID <- Blues_lab_ID[,-1]
Corr_lab_ID<- Blues_lab_ID%>% 
  dplyr::select(1,2,4)%>%
  rename("Mean germination time"=Germination_DAS, "Germination percentage"= Healthy_seedlings_Perc)

Blues_image_ID <- read_csv("Blues_image_ID.csv")
Blues_image_ID<-Blues_image_ID[,-1]
Corr_image_ID<-Blues_image_ID%>% 
  dplyr::select(1:4)%>%
  rename("Mean germination time"=Germination_DAS, "Germination percentage"= Germination_Perc,"Growth rate"=mumax)

Corr_seedvsger<-merge(Corr_image_ID,Blues_image_seedsize_ID,by="ID")
Corr_seedvsger<-merge(Corr_seedvsger,Corr_image_ID,by="ID")

Corr_seedvsger<- Corr_seedvsger%>% 
  rename("Seed size" = area_mm2)

Corr_seedvsger<- Corr_seedvsger%>% 
  dplyr::select(c(1:5))
str(Corr_seedvsger)

png("correlation_plot_seedvsgr.png",width=800, height=600)
chart.Correlation(Corr_seedvsger[,2:5],method="pearson",histogram=FALSE,pch=15)
dev.off()

library(gridExtra)
library(png)
img1<-readPNG("correlation_plot_seedvsgr.png")
img1
```
