## Supplementary material for "Phenotypic differentiation between highland and coastal quinoa under cold stress conditions": R Scripts: README.md

- **Paper.analysis.R** contain R codes for the anaylsis

---

- **acc.csv** – Contains information on the 60 accessions included in this study.  
- **fielddata_file1.csv** – Contains field experiment data for the traits *Germination*, *Days to panicle emergence*, *Plant height*, *Panicle length*, and *Panicle width* measured per plant.  
- **fielddata_file2.csv** – Contains field experiment data for the traits *Days to flowering*, *Days to maturity*, and *Seed yield*.  
- **Data_PerPlant.csv** and **Data_PerPlot.csv** – Contain field experiment data combined with growing degree days (GDD) information per plant and per plot, respectively.  
- **quinoa_field2021_miss0.80_pruned_plink2.vcf** – Genotype data file for 55 accessions.  
- **Temperature_field.csv** – Contains weather data recorded during the field experiment.  
- **Blues_ID.csv**, **Blues_ID_R_T.csv**, and **Blues_ID_T.csv** – Contain the estimated means for the field experiment data: per accession, per accession × sowing date × replication, and per accession × sowing date, respectively.  
- **mtsi_inputdata.csv** – Contains the input data used for the multi-trait stability index (MTSI) analysis.  

---

- **Lab_germinationdata.csv** and **Lab_germ.csv** – Contain raw laboratory germination data.  
- **Healthy_seedlings.csv** – Contains laboratory data on healthy seedling germination.  
- **avg_labdata.csv** – Contains average laboratory germination data.  
- **Blues_lab_ID.csv**, **Blues_lab_ID_R_T.csv**, and **Blues_lab_ID_T.csv** – Contain the estimated means for the laboratory germination data: per accession, per accession × treatment × replication, and per accession × treatment, respectively.  

---

- **image_analysis.txt** – Contains the output of image analysis for the laboratory germination data.  
- **growth_rates_fit_RT_days1-3.txt** and **growth_rates_fit_cold_days1-15.txt** – Contain seedling growth rate data derived from image analysis under respective conditions.  
- **Blues_image_ID.csv** and **Blues_image_ID_T.csv** – Contain the estimated means for the image analysis-based laboratory germination data: per accession and per accession × treatment, respectively.  
- **Cummulative-ods** – Contains cumulative germination data.  

---

- **acc_seedsize.csv** contain the saponin data for 60 accessions.
- **marvin-data-processed.ods** – Contains seed size measurements obtained using the MARViN device.  
- **corrected-output-8C.csv** – Contains seed size measurements obtained through image analysis.  
- **Blues_gen1_marvin_ID.csv** – Contains the estimated means for seed size area data for generation 1.  
- **Blues_marvin_ID.csv** and **Blues_image_seedsize.ID.csv** – Contain the estimated means of relative seed size values measured using the MARViN device and image analysis, respectively.  
