## Supplementary figures and images for "Phenotypic differentiation between highland and coastal quinoa under cold stress conditions"

### Fig_1.png

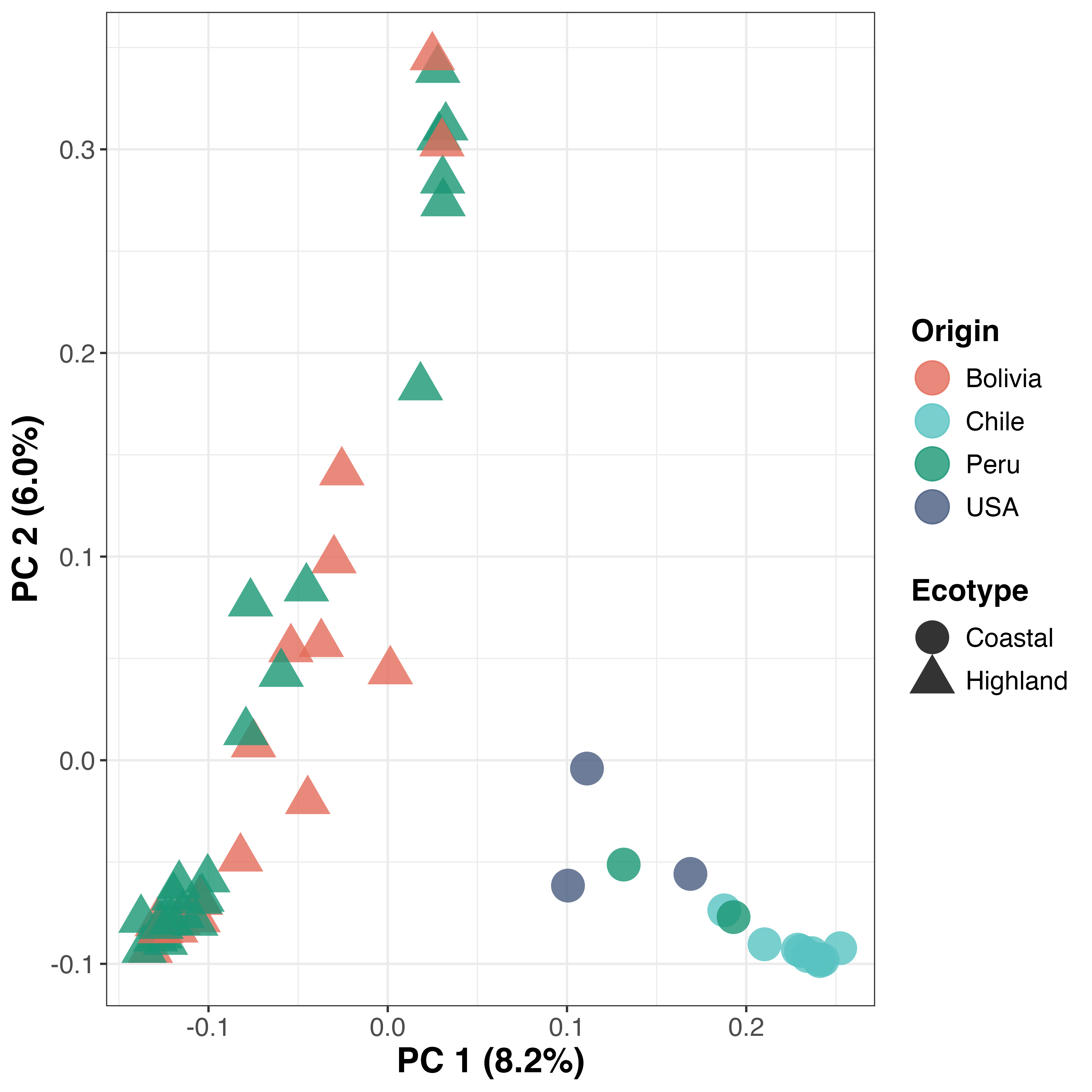
